# Cortical recycling in high-level visual cortex during childhood development

**DOI:** 10.1101/2020.07.18.209783

**Authors:** Marisa Nordt, Jesse Gomez, Vaidehi Natu, Alex A. Rezai, Dawn Finzi, Holly Kular, Kalanit Grill-Spector

## Abstract

Human ventral temporal cortex (VTC) contains category-selective regions that respond preferentially to ecologically-relevant categories such as faces, bodies, places, and words, which are causally involved in the perception of these categories. How do these regions develop during childhood? We used functional MRI to measure longitudinal development of category-selectivity in school-age children over 1 to 5 years. We discovered that from young childhood to the teens, face- and word-selective regions in VTC expand and become more category-selective, but limb-selective regions shrink and lose their preference for limbs. Critically, as a child develops, increases in face- and word-selectivity are directly linked to decreases in limb-selectivity, revealing that during childhood limb-selectivity in VTC is repurposed into word- and face-selectivity. These data provide evidence for cortical recycling during childhood development. This has important implications for understanding typical as well as atypical brain development and necessitates a rethinking of how cortical function develops during childhood.

## Main

A central question in neuroscience is how does cortical function develop? Ventral temporal cortex (VTC) is an excellent model system to address this question as it contains regions selective for ecological categories such as faces^1^, bodies^2^, places^3^, and words^4^ that are critical for human behavior and can be identified in each individual. When infants first open their eyes, they are inundated with faces, body-parts, their surrounding room, and objects. This visual input may begin to shape VTC representations in infancy and lead to the emergence of face representations in the first year of life^5-7^. However, experience with other categories, such as words, does not begin until later in childhood when children learn to read.

Two main theories regarding the development of category-selective regions have been proposed. The theory of functional refinement predicts that category-selective regions emerge from raw, general-purpose cortex^8,9^, which has some basic property^10^ that is present early in development such as foveal bias^11,12^. For example, a longitudinal study that tracked development of word-selectivity during the first year of school, found that word-selectivity emerged upon previously unspecialized cortex^9^. Similarly, developmental studies of face-selective regions reported that the growing part of face-selective regions showed less specificity in children than adults^8^ and also in younger than older infant monkeys^7^.

Alternatively, the theory of competition posits that rivalry for cortical resources may constrain development^13,14^. In particular, as both face and word recognition require fine visual acuity afforded by foveal vision, emerging representations for these categories may compete over foveal resources in lateral VTC^15^. This competition for foveal cortical territory may lead to recycling of cortex selective to one category earlier in childhood (e.g., faces) to be selective to another category (e.g., words) with later demands for reading^14,16^. For example, comparison of brains of illiterate or late literate adults to literate adults suggests that increased literacy in adulthood is associated with reduced responses to faces in the left fusiform gyrus^14,17^. Because reading acquisition is specific to humans, and the fusiform gyrus, which is the anatomical structure where face-selective regions reside, is hominoid-specific, addressing this intense theoretical debate necessitates longitudinal brain measurements in school-age children to examine the development of multiple category representations.

Here, we addressed this key gap in knowledge using longitudinal fMRI in 29 children (initially 5–12 years old) to measure development of cortical representations of multiple categories as well as their relationship. Children were scanned repeatedly over the course of 1 to 5 years (mean±SD: 3.75±1.5 years, Fig. S1A), with an average of 4.4±1.92 fMRI sessions per child and 128 included sessions overall (Methods). During fMRI, children viewed images from 10 categories spanning five domains: characters (pseudowords, numbers), body parts (headless bodies, limbs), faces (adult faces, child faces), objects (string instruments, cars), and places (houses, corridors, Fig. S1D). Analyses were performed in each individual’s native brain space, which allows precise tracking of the developing cortex in each participant and prevents blurring of responses from different categories due to normalization to a standard brain template^18^.

## Results

### How does category-selectivity develop in VTC?

To assess the development of category-selectivity in VTC, we first quantified the volume of category-selective activations in VTC as a function of age. Selectivity was computed by contrasting responses to each category vs. all other categories except the other category from the same domain (e.g., limbs vs. all other categories except bodies, t>3, voxel level). VTC was anatomically defined on the cortical surfaces of each child and divided into lateral and medial partitions (Fig. 1B). This division captures the center-periphery organization of VTC^12,15^, where lateral VTC represents the central visual field, and medial VTC represents the peripheral visual field^19^. To test for age-related changes in the volume of category-selective activations, we used linear mixed models (LMMs) with age as a fixed effect and participant as a random effect. LMMs with intercepts that varied across participants and a constant slope fit the data best in the majority of cases (Methods). Fig. 1A shows examples of linear mixed model fits for words, limbs, and faces (all LMMs in Fig. S2). Fig. 1C summarizes the LMM slopes and confidence intervals (CI) for all categories and VTC partitions. The slopes (β_age_) indicate the rate of change (mm^3^/month) of category-selective volume with age (Fig. 1C-colored bars). Significant development after FDR correction^20^ is indicated by asterisks in Fig. 1C (full statistics in Tables S1-S2).

**Fig. 1.**
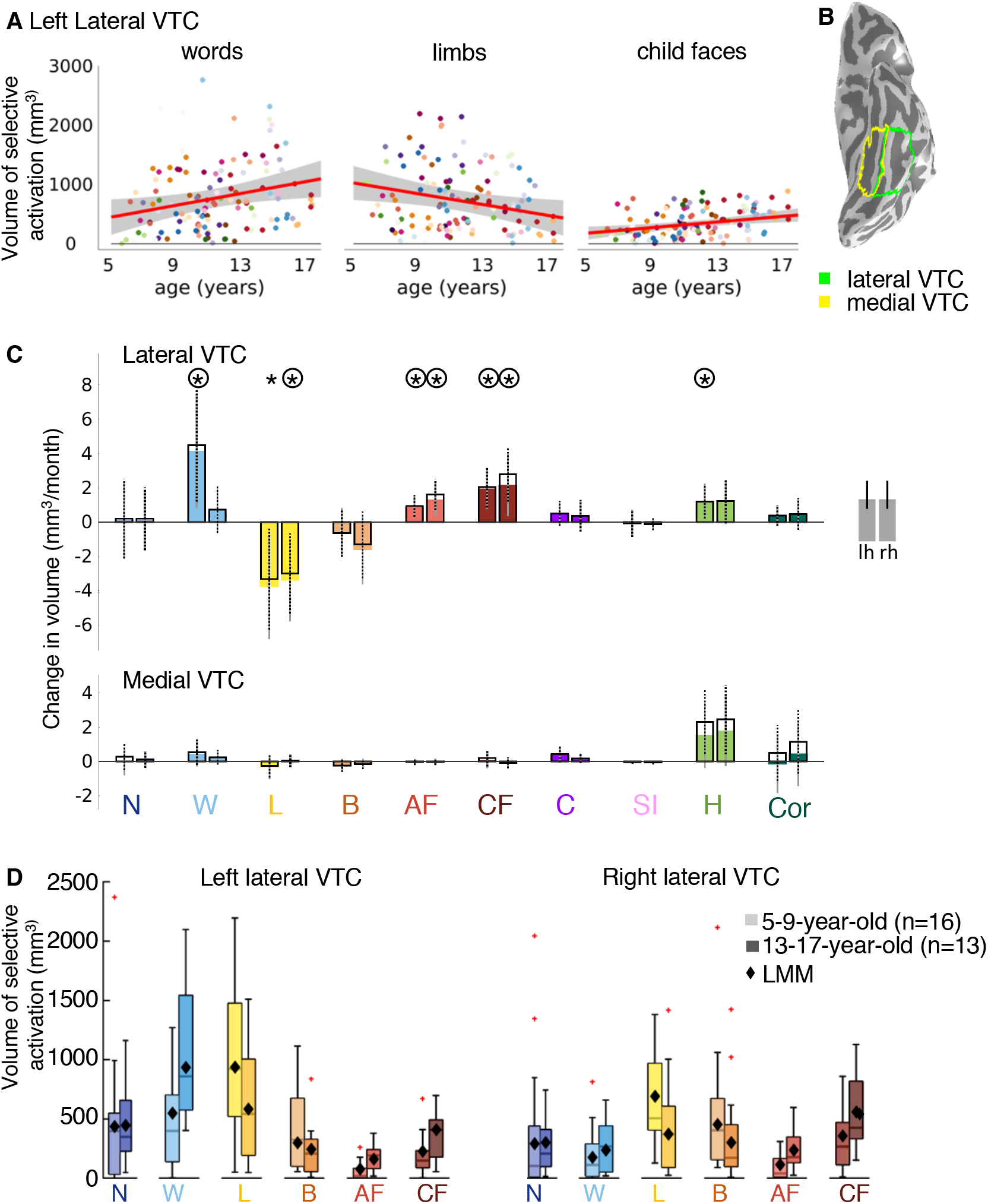
Developmental increases and decreases in category-selective activation in lateral VTC. (A) Volume of word-, limb- and child-face-selective activation by age. Each dot is a session and colored by participant. *Red line:* Linear mixed model (LMM) prediction of category-selective activation by age. Shaded gray: 95% confidence interval (CI). All scatterplots in Fig. S2. (B) Lateral and medial VTC on the inflated cortical surface of a 5-year-old. (C) LMM slopes indicating change in category-selective activation volume per month (n=128 sessions, 29 children). *Colored bars: s*lopes for models using age as a predictor. *Open bars with black outlines:* slopes for the age predictor for models including both age and time series signal-to-noise ratio (tSNR) as predictors. *Error bars:* 95% CI. Significant development after FDR-correction (p<0.05) is indicated by asterisks (colored bars) and circles (open bars). (D) Category-selective activation by age group. One session per child is included per boxplot. *Crosses:* outliers; *Diamonds:* mean volume predicted by LMM. All categories in Fig. S3. *Acronyms:* N: numbers; W: words; L: limbs; B: bodies; AF: adult faces; CF: child faces, C: Cars, SI: String instruments, H: Houses, Cor: Corridors.

Results reveal differential development of category-selectivity across VTC partitions and categories. Significant development of category-selectivity occurred in lateral, but not medial VTC (Fig. 1C, Fig. S2, Tables S1-S2). Surprisingly, there were both developmental increases and decreases in the volume of category-selective activation in lateral VTC: word- and face-selective activation increased, but limb-selective activation decreased (Fig. 1A,C, Fig. S2).

Interestingly, during childhood, the volume of word-selective activation significantly increased in left lateral VTC (Fig. 1C-light blue, β_age_=4.14 mm^3^/month (95%CI:0.81,7.48), t(126)=2.46, p_FDR_=0.044). However, there was no evidence for change in the volume of word-selective activation in the right hemisphere (Fig. 1C,D, Fig. S2, β_age_=0.66 mm^3^/month (−0.66,1.99), t(126)=0.99, p_FDR_=0.46), or of number-selective activations bilaterally (Fig. 1C, Fig. S2, Table S1, left: β_age_=0.11 mm^3^/month (−2.15, 2.38), t(126)=0.10, p_FDR_=0.92; right: β_age_=0.11 mm^3^/month (−1.69, 1.90), t(126)=0.12, p_FDR_=0.92). Examining the average volume of activations (Fig. 1D-boxplots, Fig. S3 for all categories) as well as LMM predictions of the mean volume (Fig 1D-diamonds), revealed that word-selectivity doubled on average from ~500 mm^3^ in 5–9-year-olds to ~1000 mm^3^ in 13–17-year-olds. Notably, as the volume of word-selective activation doubled, the volume of limb-selective activation halved (Fig. 1D-yellow). In fact, limb-selective activation significantly decreased over childhood in both hemispheres (Fig. 1C-yellow, left: β_age_=-3.79 mm^3^/month (−6.84,-0.74), t(126)=-2.46, p_FDR_=0.044; right: β_age_=-3.42 mm^3^/month (−5.79,-1.04), t(126)=-2.85, p_FDR_=0.04), but there was no evidence for change in body-selective activation (Fig 1C-orange, left: β_age_=-0.59 mm^3^/month (−1.98, 0.80), t(126)=-0.84, p_FDR_=0.50; right: β_age_=-1.63 mm^3^/month (−3.63, 0.38), t(126)=-1.61, p_FDR_=0.22). At the same time, the volume of face-selective activation significantly increased over childhood bilaterally for both adult faces (Fig. 1C-bright red: left: β_age_=0.89 mm^3^/month (0.26,1.52), t(126)=2.79, p_FDR_=0.04; right: β_age_=1.32 mm^3^/month (0.34,2.29), t(126)=2.68, p_FDR_=0.042) and child faces (Fig. 1C-dark red, left: β_age_=1.95 mm^3^/month (0.74,3.17), t(126)=3.18, p_FDR_=0.036; right: β_age_=2.16 mm^3^/month (0.35,3.97), t(126)=2.37, p_FDR_=0.049). Similar longitudinal development in lateral VTC was observed for other contrasts, such as contrasting responses by domain (Fig. S4) and for other metrics, such as the level of category-selectivity within constant-sized regions (Fig. S5).

A concern in pediatric imaging is that data quality may conflate developmental effects. Thus, we performed several controls to test if factors that may affect data quality contribute to our results. First, behavioral performance on the oddball task performed during scanning was overall high (median performance=91%, SD=18%), indicating that children were attending to the stimuli. Second, to test if developmental effects may be related to temporal signal-to-noise (tSNR) or motion during scan, we ran more LMMs, which included in addition to age, tSNR and average motion during scan as predictors. Motion was not a significant predictor of the volume of category-selective activation (Table S3). While tSNR was a significant predictor, it was independent from age. That is, LMMs that included tSNR as an additional predictor revealed the same pattern of results showing significant age-related increases in face- and word-selective activations as well as age-related decreases in limb-selective activations (Fig 1C-open bars, Tables S4, S5). Third, to test if developmental effects are driven by overall lower reliability of responses in younger children compared to older children, we assessed response reliability in V1 as a control ROI. Response reliability in V1 was overall high and not related to age (Fig. S6). Together, these analyses validate that age-related changes in category-selective activations reflect longitudinal development rather than differences in measurement quality across years.

### Is development anatomically specific?

To determine the anatomical specificity of the observed development, we defined limb-, word-, and face-selective regions of interest (ROIs) in each participant and session (Fig. 2A-C, t-value > 3 voxel-level) and examined them longitudinally. Category-selective ROIs remained largely within the same anatomical structures as ROIs expanded or shrank across childhood (Fig. 2A-C). We used LMMs to examine the effect of age on ROI volume (age: fixed effect, subject: random effect, change of ROI volume per month (β_age_) in Fig. 2D, full statistics in Table S6).

**Fig. 2.**
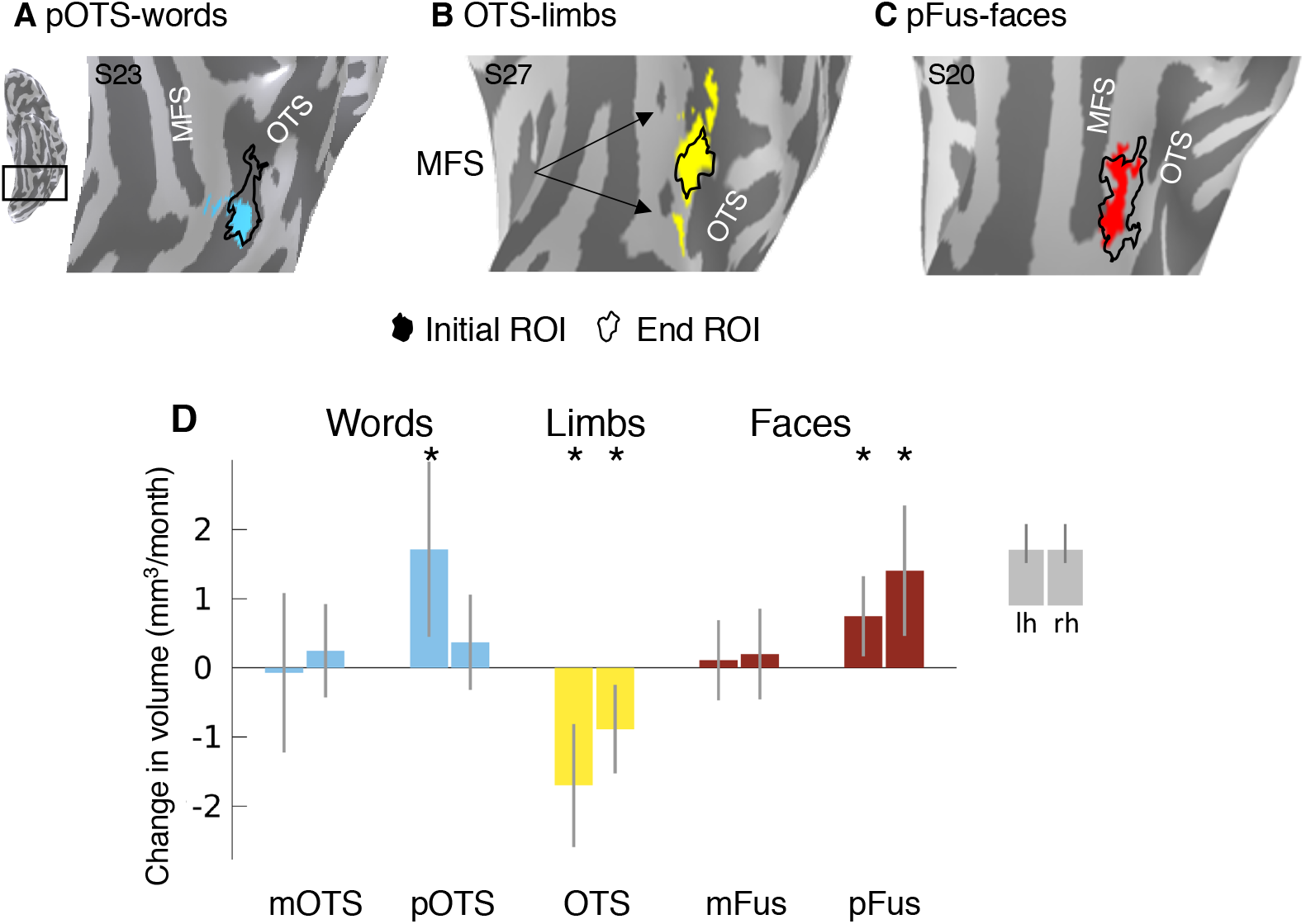
Development of category-selective ROIs. Initial ROIs (colored) and end ROIs (outline) in 3 example children: (A) left pOTS-words at age 10 and 15, (B) left OTS-limbs at age 11 and 13, (C) left pFus-faces at age 9 and 14. MFS: mid fusiform sulcus; OTS: occipito-temporal sulcus. (D) LMM slopes: change in volume of category-selective regions per month. *Error bars:* 95% CI; *Asterisks:* significant development, p<0.05, FDR-corrected. The number of sessions per ROI is as follows: left mOTS-words: n=103, right mOTS-words: n=35, left pOTS-words: n=119, right pOTS-words: n=73, left OTS-limbs: n=126, right OTS-limbs: n=126, left mFus-faces: n=107, right mFus-faces: n=102, left pFus-faces: n=120, right pFus-faces: n=98.

The growth of word- and face-selective regions was anatomically specific: posterior but not anterior ROIs significantly expanded (Fig. 2D). Activation for words grew significantly in the left posterior occipitotemporal sulcus (Fig 2D-pOTS-words, β_age_=1.71 mm^3^/month (0.45,2.98), t(117)=2.68, p_FDR_=0.02), but not in the mid occipitotemporal sulcus (Fig. 2D-mOTS-words, left: β_age_=-0.07 mm^3^/month (−1.22,1.08), t(101)=-0.12, p_FDR_=0.90, right: β_age_=0.25 mm^3^/month (−0.43,0.92), t(33)=0.74, p_FDR_=0.66). Similarly, bilateral activation for faces grew significantly in the posterior fusiform (Fig 2D-pFus-faces, left: β_age_=0.75 mm^3^/month (0.17,1.33), t(118)=2.55, p_FDR_=0.02, right: β_age_=1.41 mm^3^/month (0.46,2.35), t(96)=2.96, p_FDR_=0.019), but not in the mid fusiform (Fig 2D-mFus-faces, left: β_age_=0.11 mm^3^/month (−0.47,0.69), t(105)=0.38, p_FDR_=0.79, right: β_age_=0.20 mm^3^/month (−0.46,0.85), t(100)=0.60, p_FDR_=0.69). In contrast to the growth of pOTS-words and pFus-faces, OTS-limbs shrank significantly in both hemispheres (Fig. 2D-OTS-limbs, Table S6, left: β_age_=-1.70 mm^3^/month (−2.59,-0.81), t(124)=-3.78, p_FDR_=0.002, right: β_age_=-0.89 mm^3^/month (−1.53,-0.24), t(124)=-2.73, p_FDR_=0.02). The growth of face- and word-selective regions and the shrinking of limb-selective regions also holds for ROI surface area (Fig. S7), but it is less pronounced for ROIs defined from domain contrasts (Fig. S8). Together, these data show that the development of category selectivity is more prominent in posterior than anterior face and word-selective ROIs.

### What functional changes occur in the developing regions?

We next assessed the functional properties of the developing regions of category-selective ROIs in each participant. For words and faces, where selective voxels emerge during development, we call the difference between the end and initial ROIs the emerging pOTS-words and emerging pFus-faces. For limbs, we call the difference between the initial and end ROIs the waning OTS-limbs. We focus on the left hemisphere due to the left lateralization of the development of word-selectivity (right hemisphere in Fig. S9). LMMs were used to assess the effect of age on selectivity in emerging and waning ROIs (age: continuous fixed effect; participant: random effect). LMM slopes and their significance are shown in Fig. 3-left (stats in figure caption and Table S7). Since developing regions are not completely independent from the original ROIs, we repeated the analysis in independent ring-shaped ROIs centered on the initial functional ROIs, yielding similar findings (Fig. S10).

**Fig. 3.**
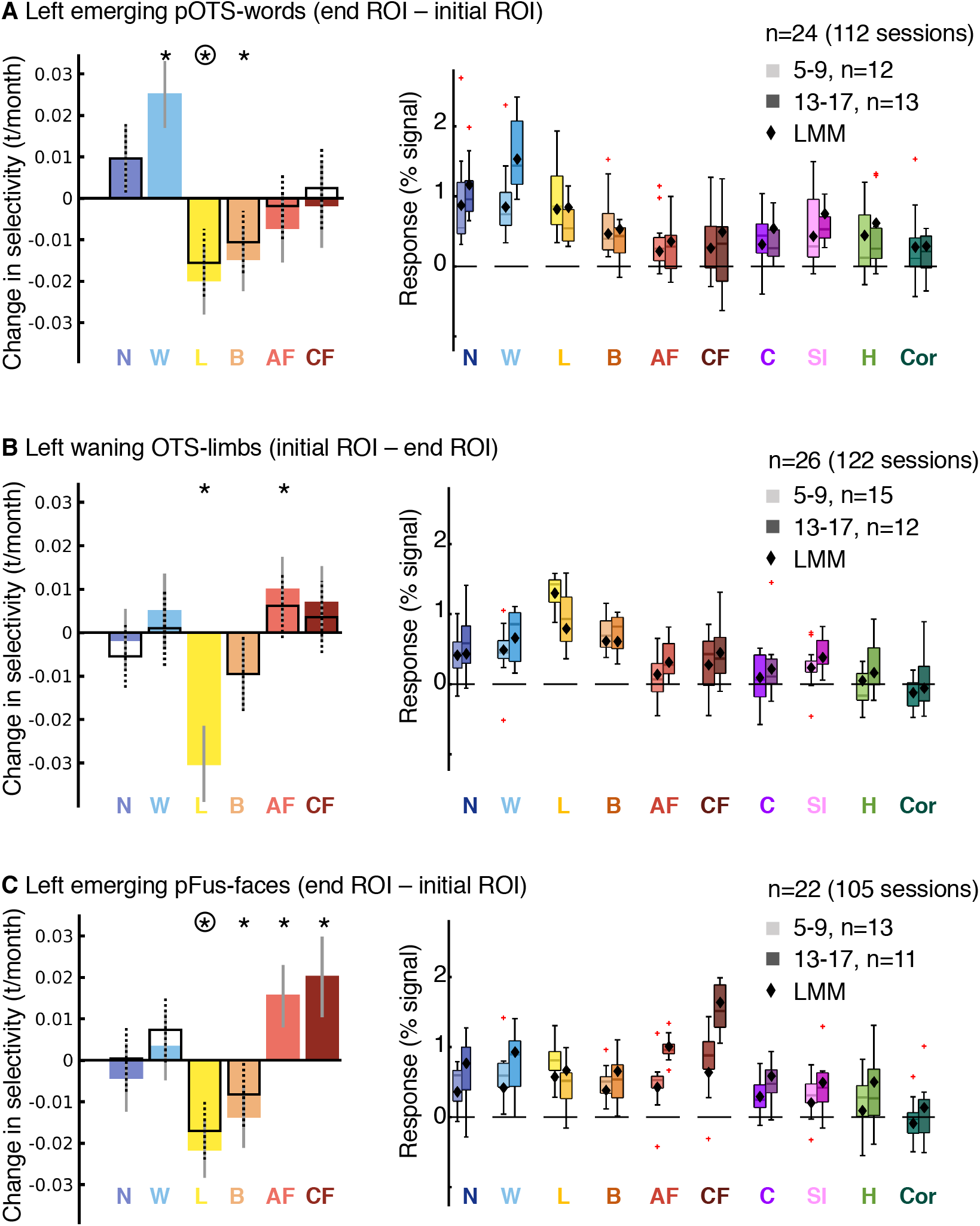
Age-related increases in word- and face-selectivity parallel decreases in limb-selectivity in the developing regions. (A-C) *Left: Colored bars:* LMM slopes indicating changes in selectivity by age. *Open bars:* LMM slopes for contrasts in which the ROI-defining category is not included. *Error bars:* 95% CI; Significant development after FDR-correction (p<0.05) is indicated by asterisks (colored bars) and circles (open bars). Note that there are no open bars or circles for the category defining the ROI. Statistics for colored bars: (A) Emerging pOTS-word, words: t(110)=6.17, p_FDR_<0.001; limbs: t(110)=-4.74, p_FDR_<0.001; bodies: t(110)=-3.68, p_FDR_=0.002. (B) Waning OTS-limbs, limbs: t(120)=-6.83, p_FDR_<0.001; adult faces: t(120)=2.59, p_FDR_= 0.04. (C) Emerging pFus-faces, adult faces: t(103)=4.09, p_FDR_<0.001; child faces: t(103)=4.11, p_FDR_<0.001; limbs: t(103)=-6.24, p_FDR_<0.001; bodies: t(103)=-3.57, p_FDR_=0.002. All ROIs and categories in Fig. S9. Full statistics in Tables S7-S8. *Right:* Response amplitudes for the 10 categories. One functional session per child is included per boxplot. *Diamonds*: estimated responses from LMMs. *Crosses:* outliers.

In the emerging left pOTS-words, selectivity to words significantly increased with age, as expected by the definition of the ROI (Fig. 3A-left blue bar). At the same time, selectivity to limbs and bodies significantly decreased (Fig. 3A-left, colored bars), but we found no evidence for changes in selectivity to other categories (Fig. S9, Table S7). As a control, we repeated these analyses excluding each region’s preferred category (e.g., excluding words in contrasts examining the emerging pOTS-words), finding similar results (Fig. 3A-left, open bars, Table S8).

To elucidate if the changes in selectivity in the emerging ROI were due to increased responses to the preferred category or decreased responses to nonpreferred categories, we used LMMs to quantify the response amplitude to each category as a function of age (age: continuous fixed effect; participant: random effect, change in %-signal/month (β_age_), Fig. S11, Table S9). Results show that developmental changes in word-selectivity were associated with significant increases in the responses to words (β_age_=0.008 %-signal/month (0.004,0.012), t(110)=3.82, p_FDR_=0.002) with no significant changes to other categories (Fig. S11, Table S9) except that responses to string instruments also significantly increased (β_age_=0.004 %-signal/month (0.001,0.006), t(110)=2.98, p_FDR_=0.025). Indeed, average responses to words in pOTS-words are higher in teens (13-17-year-olds) than children (5-9-year-olds; Fig. 3A-right-boxplots, Fig. S11, Table S9) in correspondence with LMM predictions (Fig. 3A-right-diamonds).

Similarly, in the emerging pFus-faces, selectivity to faces increased (Fig. 3C-left-red bars, Fig. S9). At the same time, selectivity to limbs significantly decreased bilaterally (Fig. 3C-left-yellow bar) and this development was significant in the left hemisphere also when faces were excluded from the contrast (Fig. 3C-left, open bars); selectivity to bodies only decreased in the left hemisphere (Fig. 3C-orange). We found no significant changes in selectivity to other categories (Fig. S9, Table S7). Increases in selectivity to faces were associated with significant increases in responses to faces (Fig. S11, Table S9, adult faces: β_age_=0.006 %-signal/month (0.004,0.009), t(103)=5.33, p_FDR_<0.001; child faces: β_age_=0.011 %-signal/month (0.007,0.015), t(103)=5.79, p_FDR_<0.001). Indeed, average responses to both adult and child faces were higher in teens than children in correspondence with LMM predictions (Fig. 3C-right). Additionally, responses to words and string instruments also significantly increased in the left emerging pFus-faces (words: β_age_=0.006 %-signal/month (0.002,0.01), t(103)=2.78, p_FDR_=0.04, string instruments: β_age_=0.003 %-signal/month (7.7×10^-4^,0.006), t(103)=2.62, p_FDR_=0.04), and there was a trend for an increase in responses to cars (Fig. 3C-right, Fig. S11, Table S9). Thus, developmental increases in word- and face-selectivity in emerging word and face ROIs, respectively, are driven by increased responses to their preferred category, rather than decreased responses to nonpreferred categories.

As OTS-limbs is located between pOTS-words and pFus-faces, we asked if increased responses to faces and words also occur in the waning OTS-limbs. We found that not only did responses to limbs significantly decrease with age (Fig. S11, β_age_=-0.005 %-signal/month (−0.008,-0.003), t(120)=-4.47, p_FDR_<0.001) but also responses to adult faces significantly increased with age (Fig. S11, β_age_=0.002 %-signal/month (4.6×10^-4^,0.003), t(120)=2.64, p_FDR_=0.043). This is an intriguing phenomenon in which this waning region responds more strongly to limbs in 5-9-years-olds than in 13-17-year-olds (Fig. 3B-right-yellow). There was no evidence for changes of responses to other categories (Fig. 3B-right, Fig. S11, Table S9-full statistics). These changes in responses amplitudes resulted in significant decreases in limb-selectivity (Fig. 3B-left, Fig. S9) as well as significant increases in face-selectivity (Fig. 3B-left, colored bars). There was no evidence for changes in selectivity to other categories including bodies (Fig. S9). Therefore, developmental decreases in limb-selectivity of the waning OTS-limbs reflect decreased responses to the preferred category together with increased responses to faces.

Given the profound developmental decreases in limb-selectivity in both emerging and waning ROIs, we tested if this is a general phenomenon across lateral VTC. Analyses of lateral VTC excluding voxels that were selective in the first session to categories showing development, revealed no significant decreases in limb-selectivity in the remainder of lateral VTC (Fig. S12). This suggests that developmental changes in limb-selectivity are most prominent in the emerging and waning ROIs.

### Are changes in face-, word-, and limb-selectivity linked?

While the theory of competition does not make predictions about limb representations, it predicts that development of face and word representations are linked as they compete over cortical territory that shows a foveal bias^13,14^. Thus, we tested if there is a quantitative relationship between selectivities to faces, words, and limbs in the emerging and waning ROIs. Model comparison of LMMs relating the selectivity to the preferred category to selectivity of the other categories revealed that in all developing ROIs, selectivity to the preferred category was better predicted by the selectivity to the other two categories rather than just one of them (likelihood ratio tests comparing a one-predictor-LMM with a two-predictor-LMM, left hemisphere: all χ^2^≥4.76, p≤0.029, Table S10). Moreover, in all developing ROIs, selectivity to the preferred category was significantly and negatively related to selectivity to the other two categories (β_category1_, β_category2_: fixed effects, subject: random effect, Fig. 4, left hemisphere; Fig. S13, right hemisphere; statistics in figure caption and full statistics in Table S11). E.g., in the emerging pOTS-words, higher word-selectivity is significantly linked with both lower face- and limb-selectivity (Fig. 4A, Table S11). The negative relationship between the preferred category of an ROI and the other two categories was also observed at the voxel level (Fig. S14).

**Fig. 4.**
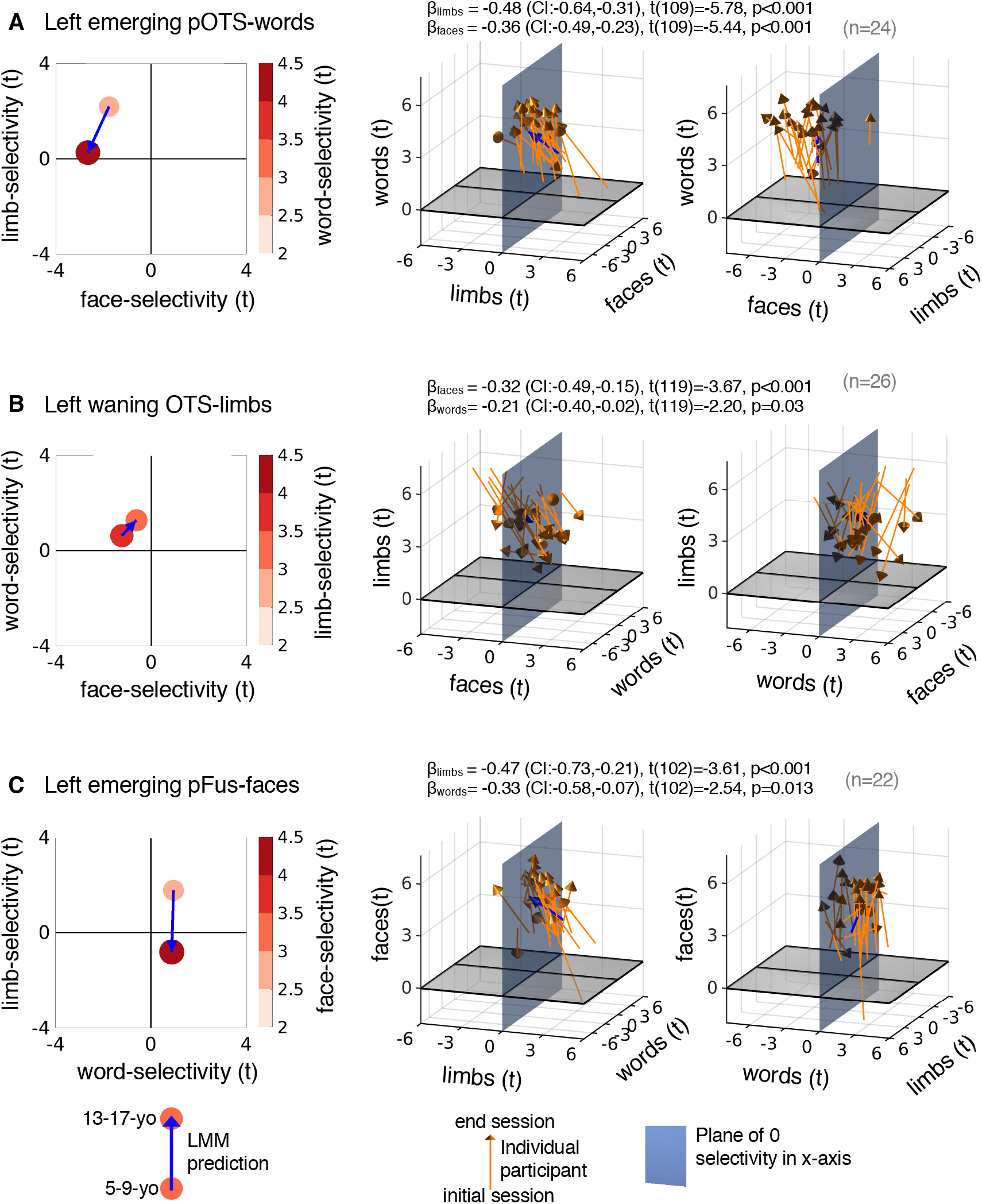
Developmental changes in word-, face-, and limb-selectivity are linked. *Left:* LMM prediction (circle) of category selectivity to words (A), limbs (B), and faces (C) vs. selectivity to the other two categories in 5-9-year-olds and 13-17-year-olds. *Middle:* Individual participant data visualized in 3D. In each panel the variable on the z-axis is related to the x- and y-variables, this relationship is quantified in the model; *βs*, 95%-CI, t-values, degrees of freedom, and p-values are shown at the top. Full statistics in Table S11; *Orange arrows:* Individual child data. *Blue arrows:* LMM prediction (same as left panel). *Right:* Rotated version of the plots in the middle column to enhance visibility of positive and negative values along the other horizontal axis. Right hemisphere in Fig. S13.

We visualized how selectivity to words, faces, and limbs changes in emerging and waning ROIs. Using the LMM, we related the selectivity to the preferred category with the selectivity to the other two categories for 5-9-year-olds and for 13-17-year-olds (Fig. 4-left, Table S12). In the emerging pOTS-words, 5-9-year-olds have positive selectivity to limbs and negative selectivity to faces (Fig 4A). By age 13-17, word-selectivity has increased while limb-selectivity has reduced to zero and face-selectivity has become even more negative (Fig. 4A-left). That is, after development, selectivity to words in pOTS-words has replaced the initial selectivity for limbs, not faces. Notably, this developmental pattern is visible in individual children from their initial age (Fig. 4A-middle & right, arrow bases) to their end age (Fig. 4A-middle & right, arrow heads). Similarly, in the emerging pFus-faces, 5-9-year-olds have positive limb-selectivity but no clear preference for words (Fig. 4C-right). By age 13-17, as face-selectivity has increased, limb-selectivity is lost and there is little change to word-selectivity (Fig. 4C Fig. S13, Table S12).

In the waning OTS-limbs, 5-9-year-olds exhibit largely negative face-selectivity and mild positive word-selectivity (Fig. 4B-left, Table S12). As limb-selectivity declines by age 13-17, both word- and face-selectivity increase (Fig. 4B-left). While limb-selectivity consistently declined across individuals, there was more variability in individual developmental trajectories compared to the emerging ROIs (Fig. 4B-middle, right). In some children limb-selectivity was replaced by word-selectivity and in others with face-selectivity.

## Discussion

Our longitudinal measurements in children using a large range of ecologically-relevant categories reveal new insights about the functional development of high-level visual cortex. We show that while face- and word-selective regions grow and become more selective, limb-selective regions shrink during childhood development and lose their selectivity to limbs. Importantly, the decrease in limb-selectivity is directly linked to the increase in word- and face-selectivity providing evidence for cortical recycling during childhood development. We discuss each of these findings and their implications below.

A surprising finding from our study is that childhood development is not only associated with growth of category-selective regions and increases in selectivity, but also involves loss of selectivity. That is, in addition to finding growth and increased selectivity of face- and word-selective regions in VTC as individual children develop, consistent with cross-sectional studies^8,12,16,21–24^, we find that limb-selective regions in VTC shrink during childhood development and lose their selectivity to limbs. This decrease in the amount of limb-selective volume is specific to limbs as the amount of body-selective volume remains largely unchanged (Fig. 1C), consistent with previous findings^9,22^. As such, our results show that young children’s VTC is actually more selective for limbs than it is later in childhood.

Our results provide striking empirical evidence for recycling^14^ of category-selectivity in high-level visual cortex during childhood. However, contrary to previous predictions^13,14^ that face-selectivity is recycled to word-selectivity during development, our results show that limb-selectivity is recycled to both word- and face-selectivity. This recycling occurs via a mechanism of decreasing responses to limbs and increasing responses to both faces and words at the ROI and voxel levels. Future research is needed to determine if this recycling also occurs at the single neuron level^25^.

Critically, our results require a rethinking of prevailing developmental theories that propose that cortical development involves sculpting of new representations upon general-purpose cortex^8,9^. First, in contrast to the prevailing view suggesting that children’s VTC is indistinctive^8-10,16,26^, the present data shows that young children’s’ VTC is more selective to limbs than it is later in childhood. Second, our data suggest that during childhood, cortical selectivity can change from one category to another.

These results generate new questions for future research. First, why do young children exhibit large VTC representations for limbs? Some clues to this question can be gleaned from behavioral studies that examined what infants and toddlers look at in natural settings. These studies discovered that young infants (≤ 6 months) look at faces more than hands, but older infants and toddlers (1-2 year-olds) look at hands more than faces^27-29^. Developmental psychologists have hypothesized that this change in the child’s “visual diet” may be related to multiple factors including the mobility of the child^29^, their motor dexterity at manipulating objects^27,29^, and communicative information in gesturing^30^. Based on this developmental literature and our findings, we speculate that much like baby teeth changing to permanent teeth, as children’s diet and size changes, cortical recycling in VTC may reflect adjustment to changing visual demands during childhood. Thus, we propose an intriguing new hypothesis that cortical recycling in VTC co-occurs with changes in the saliency and frequency of visual stimuli that are socially and communicatively relevant, moving from hands and gestures in toddlers to faces and words in school age children and teens. Future research can test this hypothesis by determining what children and teens look at together with computational modeling to test the effect of different visual diets on emerging VTC representations^31-33^.

Second, what are the behavioral implications of this cortical recycling? Prior research has found that developmental improvements in face recognition ability^8,34^ and face discriminability^35^ are linked with developmental increases in face selectivity^8,34^ and neural sensitivity to faces^35^ in VTC, respectively. Likewise, developmental improvements in reading proficiency are linked with developmental increases in neural selectivity to words^9,23,36^ and in the informativeness of distributed responses to words in VTC^12^. These studies suggest that the developmental decreases in responses to limbs in VTC may have behavioral ramifications, a hypothesis that can be tested in future developmental research combining neuroimaging and behavioral measurements.

The present findings have important implications for understanding typical^37^ and atypical^38-41^ brain development. First, these data fill a key gap in knowledge by quantifying the rate of the development of category-selectivity from young children to teens. Thus, they offer a foundation for using fMRI to assess developmental and learning deficiencies, especially those related to reading^42^ and social perception^38,39^. We acknowledge that a limitation of the current study is the variability in data acquisition with regard to the time interval between scans and number of scans per child. Thus, we emphasize the necessity for future longitudinal research with more regular age sampling spanning a large duration in both typical and atypical populations, which will allow further in-depth analyses of cortical recycling at finer spatial scales, such as on the level of individual voxels. Second, it will be important to determine if there is a critical period during development in which cortical recycling can occur^10,43^ and if this period is particularly protracted in lateral VTC, which overlaps foveal representations^11,15,19^. Third, childhood visual deprivation^40^ or brain lesions^41^ may affect cortical recycling and the emergence of category representations in VTC in atypical populations^44-47^. Future longitudinal studies measuring cortical development in children who have vastly different visual experience with faces, hands, and written words (e.g., congenital blind, sighted, and congenital deaf), will be important for determining how differing visual inputs and behavioral demands during childhood affect cortical recycling. Finally, an open question for future research is whether cortical recycling occurs in other brain systems in which earlier representations may be altered by schooling^48^ or prolonged childhood experience. Some examples include learning new languages after one’s mother tongue^49,50^ or learning complex math^51^ upon an earlier numerosity system^52,53^.

In sum, the surprising finding from our study is that contrary to the prevailing hypothesis that during childhood new cortical representations are formed upon unspecified cortex, we find evidence for cortical recycling in which limb representations are repurposed during childhood to represent faces and words. The discovery of cortical recycling is important because it not only provides a key advancement in understanding cortical development, but also necessitates a rethinking of how cortical function develops during childhood.

## Methods

### Statement on ethical regulations

This study was approved by the Institutional Review Board of Stanford University and complies with all relevant ethical regulations. Prior to the start of the study, parents gave written consent, and children gave written assent. For their participation children received $30 per hour for scanning, as well as a small toy.

### Participants

Children with normal or corrected-to-normal vision were recruited from local schools in the Bay Area. The diversity of the participants reflects the makeup of the Bay Area population. 62.5% of children were Caucasian, 20% were Asian, 5% were Native Hawaiian, 5% were Hispanic, and 7.5% were multiracial or from other racial/ethnic groups.

Prior to their first MRI session, children were trained in a scanner simulator to acclimate them to the scanner environment and to enhance quality of MRI data. In the simulator, children practiced laying still while watching a short movie and receiving live feedback on their motion. For subsequent scans, simulator training was repeated if necessary.

We collected data from 40 (26 female) children (onset age=5-12 years, M=8.66 years, SD=2.34 years, Fig. S1). We selected this age range because (i) it captures the phase in which children start learning to read and (ii) it covers a broad age range spanning childhood and adolescence in which previous studies have documented VTC development^9,21^.

Data from 4 children were excluded because they dropped out of the study after participating only once, and thus did not provide longitudinal data. Data from 7 children were excluded because their data did not pass inclusion criteria (see below). In the remaining 29 children, 29 functional sessions were excluded due to motion, 1 session due to a technical error during acquisition, and 1 session due to aliasing artifacts during acquisition. Therefore, data from 128 functional sessions of 29 neurotypical children (18 female, 11 male) are reported in this study (Fig. S1A,B has an overview of the included and excluded sessions). Initial ages of the included children ranged from 5 to 12 years (mean=9.19, SD=2.13). No statistical methods were used to pre-determine sample sizes, but our sample sizes are similar to those reported in previous cross-sectional publications^34,54^ and larger than previous longitudinal studies on VTC development^9^.

Participants were scanned using functional and structural MRI for 1 to 5 years. When possible, children participated in 1 to 2 functional scans per year. Additionally, children participated in 1 structural MRI session per year. Each child participated in at least 2 and up to 10 fMRI sessions (mean=4.41, SD=1.92) with the time interval between the first and last fMRI scan ranging from 10 months to 5 years (mean=45 months, SD=18 months, Fig. S1C). Functional and anatomical scans were typically conducted on different days to avoid fatigue.

### Magnetic resonance imaging

#### Structural imaging

Data were acquired at the Center for Cognitive Neurobiological Imaging at Stanford University on a 3 Tesla GE Discovery MR750 scanner (GE Medical Systems) using a phase-array 32 channel head coil. Whole brain anatomical scans were collected using quantitative MRI (qMRI^55^) with a spoiled gradient echo sequence using multiple flip angles (*α*=4°, 10°, 20°, 30°), TR=14ms and TE=2.4ms. The scan resolution was 0.8×0.8×1.0mm^3^ (later resampled to 1mm isotropic). For T1-calibration we acquired spin-echo inversion recovery scans with an echo-planar imaging read-out, spectral spatial fat suppression and a slab inversion pulse. These scans were acquired at TR=3s, inplane resolution=2×2mm^2^, slice thickness=4mm and 2x acceleration, echo time=minimum full.

#### Functional imaging

Functional data were collected using the same scanner and head coil as the structural images. Slices were oriented parallel to the parieto-occipital sulcus. The simultaneous multi-slice, one-shot T2* sensitive gradient echo EPI sequence was acquired with a multiplexing factor of 3 to acquire near whole brain coverage (48 slices), FOV=192mm, TR=1s, TE=30ms, and flip angle=76°. Resolution was 2.4 mm isotropic.

#### 10 category experiment

Participants completed three runs of a 10-category experiment^11,12,54^. During each run participants viewed images from five domains each comprising images of two categories including faces (adult faces, child faces), body parts (headless bodies, limbs), objects (cars, string instruments), places (corridors, houses) and characters (pseudowords, numbers). Following prior work on visual representations, we define a visual category as a set of exemplars sharing the same parts and configuration, e.g., limbs^56-59^ and domain as a grouping of one or more categories that share semantic association (whether or not they share visual features) and are thought to require distinct processing mechanisms^60^. E.g., houses and corridors are both places, but are visually dissimilar. Examples of stimuli are shown in Fig. S1D.

Images were presented in 4 s blocks, at a rate of 2 Hz and did not repeat across the course of the experiment. Image blocks were intermixed with gray luminance screen baseline blocks. Blocks were counterbalanced across categories and baseline. Stimuli were grayscale and contained a phase-scrambled background generated from randomly selected images. Participants were instructed to view the images while fixating on a central dot and to perform an oddball task. Participants pressed a button whenever an image comprising only the phase-scrambled background appeared. Due to occasional button box malfunction behavioral responses in the oddball task were recorded in 98 out of 128 sessions used in the analyses.

### Data analysis

Data analysis was performed in MATLAB version 2017b (The MathWorks, Inc.) and using the mrVista software package (https://github.com/vistalab/vistasoft/wiki/mrVista).

#### Inclusion criteria

In each functional session children participated in three runs of the 10 category-experiment. Criteria for inclusion of data were (i) at least 2 runs per session where within-run motion < 2 voxels and between-run motion < 3 voxels, and (ii) at least two fMRI sessions at least six months apart. Because only two of the three runs survived motion quality thresholds for several fMRI sessions, analyses include two runs per child per session to ensure equal amounts of data across participants and sessions. For sessions with all 3 runs passing motion quality criteria, 2 runs with lowest within-run motion were included.

#### Structural MRI data analysis and individual template creation

Quantitative whole brain images of each child and timepoint were processed with the mrQ pipeline (https://github.com/mezera/mrQ^55^) to generate synthetic T1 brain volumes. For each child, the synthetic T1 brain volumes from their multiple timepoints were used to generate a within-subject brain volume template. Each participant’s brain anatomical template was generated using the FreeSurfer Longitudinal pipeline (https://surfer.nmr.mgh.harvard.edu/fswiki/LongitudinalProcessing^61^) using FreeSurfer version 6.0. The gray-white matter segmentation of each participant’s within-subject brain template was manually edited to fix segmentation errors (e.g., holes and handles) to generate an accurate cortical surface reconstruction of each participant’s brain. The motivation for aligning the functional data to the within-subject-template were (i) to enable comparison of regions of interest (ROIs) from different timepoints in the same brain volume for each participant and to (ii) minimize potential biases which can occur from aligning longitudinal data to the anatomical volume from a single timepoint^61^. On average 2.48 (SD=0.69) synthetic T1s were used to generate the within-subject-template (min=2, max=5). Functional data from all sessions of a participant were aligned to their within-subject brain template. In 17 participants the last fMRI session that was included was conducted after the within-subject template had been created. These functional sessions were acquired on average 11± 2 months after acquisition of the last synthetic T1 that was included in the within-subject-template (excluding 2 participants whose last T1 could not be used because of technical error during acquisition and subject motion).

#### Definition of anatomical regions of interest (lateral and medial VTC ROIs)

On the inflated surface of each hemisphere in each participant, we defined anatomical regions of interest (ROIs) of the lateral and medial VTC (Fig. 1B) as in previous publications^12^. We first defined the VTC anatomically and then separated it to lateral and medial VTC. The posterior border of VTC was the posterior transverse collateral sulcus (ptCoS) and the anterior border was aligned to the posterior end of the hippocampus, which typically aligns with the anterior tip of the mid fusiform sulcus (MFS). The lateral border of VTC was the inferior temporal gyrus (ITG) and the medial border of VTC was the medial border of the collateral sulcus (CoS). Finally, VTC was divided into its lateral and medial partitions along the MFS (example in Fig. 1B).

#### fMRI data analysis

Functional data from each session were aligned to the individual within-subject template. Motion correction was performed both within and across functional runs. No spatial smoothing and no slice-timing correction were performed. Time courses were transformed into percentage signal change by dividing each timepoint of each voxel’s data by the average response across the entire run. To estimate the contribution of each of the 10 conditions (corresponding to the 10 image categories) a general linear model (GLM) was fit to each voxel by convolving the stimulus presentation design with the hemodynamic response function (as implemented in SPM, www.fil.ion.ucl.ac.uk/spm).

#### Definition of selectivity in lateral and medial VTC

We used a data-driven approach to examine the development of category selectivity in VTC. The motivation for this type of analysis was to (i) use an automated, observer-independent approach and (ii) use an approach that does not require clustered activations.

In each participant, we assessed selectivity to each category in anatomically defined lateral and medial VTC ROIs (**Fig. 1B**). Selectivity was defined as t-value > 3 (voxel-level) for the contrast of interest. We also performed a complementary threshold-independent analysis in constant-sized regions (see Control analysis below). Category contrasts (Fig. 1) were computed by contrasting responses to each category with all other categories from all other domains (i.e., words vs. all other categories except numbers). We used this approach for defining contrasts to ensure that all categories are contrasted to the same number of control categories and contrasts are not biased to any particular category. We also computed contrasts for domains (faces, body parts, characters, objects, places), see supplemental analyses (Fig. S4). For domain contrasts, responses to both categories of each domain were contrasted with stimuli from all other domains (i.e., characters vs. all others).

#### Definition of functional regions of interest

To examine the anatomical specificity of the observed development of selectivity (Fig. 1), we defined word-, limb-, and face-selective functional regions in each participant (see Fig. 2A-C). For the definition of functional ROIs, a threshold of a t-value > 3 (voxel-level) was used. All ROIs were defined in each participant’s native cortical surface generated from the within-subject brain template. Word-selective regions were defined as above-threshold clusters for the contrast words vs. all categories except numbers that straddled the occipito-temporal-sulcus (OTS). The more anterior cluster was defined as mOTS-words and the posterior cluster as pOTS-words. These clusters are also referred to as the visual word form area (VWFA 1 and 2, respectively^4^). The limb-selective region was defined as above-threshold cluster of voxels for the contrast limbs vs. all categories except bodies straddling the OTS and was labelled OTS-limbs. This cluster is also referred to as the fusiform body area (FBA^2^). Face-selective regions were defined using the contrast faces (adult and child) vs. all other categories, because we observed similar development for both face types in the analysis related to Fig. 1. Face-selective clusters were defined as above-threshold clusters on the lateral fusiform gyrus. The more anterior cluster typically aligns with the anterior tip of the MFS, and was defined as mFus-faces, while the more posterior cluster was defined as pFus-faces^19^. These two face-selective clusters are also referred to as the fusiform face area (FFA 1 and 2, respectively^1^). For supplemental analyses (Fig. S7) we also defined place-, character-, and combined body-part-selective regions. Place-selective regions were defined as above threshold clusters on the collateral sulcus that respond to houses and corridors vs. all other categories. This region is also referred to as parahippocampal place area (PPA^3^). Character-selective regions were defined as above-threshold clusters on the OTS that responded more strongly to words and numbers vs. all other categories. Similarly, a body-part-selective region was defined as above-threshold clusters on the OTS that responded more strongly to bodies and limbs than the other categories.

#### Emerging and waning ROIs

To characterize the selectivity and responses within VTC regions that changed with development, we defined in each participant’s VTC ROIs that: (i) either gained selectivity to faces or words or (ii) lost selectivity to limbs during childhood. To do so, we defined emerging face and word ROIs as well as waning limb ROIs. For words and faces, where selective voxels emerge during development, the emerging ROI is the region that was selective to faces (or words) in the end timepoint but was not selective to that category in the initial timepoint (Fig. 2A,C). We call the region that is the difference between the end and initial ROIs the emerging pOTS-words and emerging pFus-faces. For limbs, where selective voxels decline during development, we call the difference between the initial and end ROIs the waning OTS-limbs. That is, the waning OTS-limbs is the region that was within the ROI in the initial timepoint but not in the end timepoint (Fig. 2B). For a given participant, the initial ROI corresponds to the first fMRI session in which the ROI could be identified, and end ROI corresponds to the last fMRI session in which the ROI could be identified.

#### Control analysis of selectivity development in independent ring-shaped ROIs

The emerging and waning ROIs were defined from individual participant’s functional ROIs on their native brain anatomy and thus capture precisely the part of cortex that undergoes development. As these ROIs are not completely independent from the original ROIs, we sought to validate these results in an independent manner. Thus, we conducted a complementary analysis to evaluate responses in independent ring ROIs. First, we created two disk ROIs centered at the center coordinate of the initial functional ROI. One ROI was sized to match the surface area of the initial ROI, and the other, sized to match the corresponding end ROI. Then, we defined the area in between these two disk ROIs as the independent ring-shaped ROI.

Within each ring-shaped ROI, we measured responses to the 10 categories as well as selectivity to each category (Fig. S10). Importantly, this analysis replicated the significant decrease in limb-selectivity in the waning limb-ROIs, emerging left pOTS-words, and emerging left pFus-faces. In right pFus-faces, we found a similar trend for decreasing limb-selectivity (see figure legend of Fig. S10 for statistics). In left pOTS-words, we find a similar trend for increasing word-selectivity (Fig. S10A) and in right pFus-faces we find a similar trend for increasing face-selectivity (Fig. S10D,E). We note that a limitation of this approach is that while the ring analysis guarantees independence, it does not capture exactly the developing tissue, as the actual developing ROIs are not ring shaped (Fig. 2A-C, see example ROIs). Therefore, this approach is less accurate for assessing developmental changes in VTC.

#### Control analysis: estimating changes in selectivity across development in a constant number of lateral VTC voxels

In the analyses in Fig. 1, we evaluated category-selectivity by estimating the number of voxels within an anatomical ROI that significantly responded to one category (vs. all other categories except the other category from the same domain, threshold of t>3, voxel-level). While this threshold guarantees significant selectivity, ultimately the threshold value is a number decided by the experimenter.

To ensure that findings of developmental effects do not depend on the threshold used, we performed a complementary analysis of category-selectivity in lateral VTC across development which did not depend on a threshold. Across development, we kept the number of voxels within VTC constant and evaluated their mean selectivity (t-value) to a category. Specifically, at each timepoint we selected the 20% most selective voxels (i.e., the voxels with the highest t-values) for each category contrast and calculated their mean t-value (Fig. S5). The 20% most selective voxels are determined in each session independently, to avoid biasing the selection of voxels to a specific timepoint. Then we used LMMs to determine if the mean selectivity of these voxels changes over time. LMMs can be expressed as: t-value ~ age in months + (1| participant).

Results of this analysis largely replicate the main finding presented in Fig. 1C, and reveal (i) a significant increase in word-selectivity in the left hemisphere (see legend of Fig. S6 for statistics), (ii) a bilateral decrease in limb-selectivity, (iii) an increase in selectivity to adult faces in the right hemisphere, (iv) a bilateral increase in selectivity to child faces and (v) an increase in selectivity to houses in the left hemisphere.

#### Control analysis: Estimating the response reliability in V1 and testing if it is related to age

To assess the reliability of responses, we conducted a multivariate pattern analysis (MVPA^62^) in V1 ROIs defined by the Glasser atlas^63^ as well as lateral VTC ROIs. For each voxel, the response amplitude (beta value) for each category was estimated from the GLM of each run. These vectors of responses - also called multivoxel patterns (MVPs) - were generated independently for each of the two runs in each session. We then calculated correlations between pairs of MVPs (run1 to run2) to each category combination, resulting in 10×10 representational similarity matrices. We calculated the mean of reliability across all categories (mean of the on-diagonal correlations) as a measure for the reliability of responses, as it provides a measurement for the similarity of responses to the same category from one functional run to another. Next, we tested if there is a relationship between the reliability of responses and age. To this end, we ran linear mixed models predicting the response reliability by age using random intercept models with the grouping variable subject (response reliability ~age + (1|subject)). Results of this analysis are in Fig. S6.

#### Is the decrease in limb-selectivity uniform across lateral VTC?

As we found accumulating evidence for reductions in limb-selectivity in lateral VTC, an open question is whether the decrease in limb-selectivity occurs across the entire lateral VTC or whether it is restricted to the regions in which selectivity changes across development. To this end, we aimed to assess development of limb-selectivity in VTC excluding voxels that were selective to categories showing significant development (Fig. 1). Thus, we identified in each participant’s first session the voxels selective for these categories, excluded them from lateral VTC, and then measured limb-selectivity across development in the remaining lateral VTC voxels. Next, we used LMMs to test if there was a significant decrease in the mean selectivity to limbs in these remaining lateral VTC voxels (Fig. S12). While there was a trend for a decrease in selectivity in both hemispheres, the effects were not significant after correcting for multiple comparisons (left: slope=-0.0039 t/month, p=0.16; right: slope=-0.0036 t/month, p=0.16, FDR corrected). These results suggest that the decrease in limb-selectivity appears to be strongest in the developing category-selective regions.

#### Can cortical recycling be measured at the voxel level?

To assess if cortical recycling is also observable at the voxel level, we measured the selectivity for limbs, faces, and words in each session for each voxel of the waning and emerging parts of ROIs. Using this data, we examined the relationship between the voxel selectivity to the category that defines the ROI as a function of voxel selectivity to the two other categories (i.e., in emerging pOTS-words we related word-selectivity to limb- and face-selectivity: word-selectivity ~ limb-selectivity + face-selectivity +(1| session). To quantify the relationship, we ran a separate linear mixed model (LMM) for each participant and ROI. Thus, the number of independent measurements in each LMM is the number of voxels in the emerging/waning ROI and the grouping variable in the LMM is ‘session’. We then summarized the individual subject slopes from the LMM for each ROI in a boxplot (Fig. S14)

#### Statistical Analyses

Linear mixed models (LMMs) were used for statistical analyses because (i) the data has a hierarchical structure with sessions being nested within each participant, and (ii) sessions were unevenly distributed across time (Fig. S1A). Models were fitted using the ‘fitlme’ function in MATLAB version 2017b (The MathWorks, Inc.). In initial analyses the fit of random-intercept models, which allow intercepts to vary across participants, were compared with the fit of random-slope models, which allow both intercepts and slopes to vary across participants. Results revealed that a random-intercept model fitted the data best in the majority of cases. Thus, to enable comparability across analyses, LMMs with random intercepts were used throughout the analyses. Data distribution was assumed to be normal, but this was not formally tested.

LMMs related to Figure 1 can be expressed as: volume of selective activation for a category in mm^3^ ~ age in months + (1| participant), in which volume of selective activation is the response variable, age in months is a continuous predictor (fixed effect), and the term (1 | participant) indicates that random intercepts are used for the grouping variable participant. The slopes of the LMMs are plotted in Fig. 1C. Statistics were run on the complete data set including 128 sessions.

Boxplots in Fig. 1D showing subsets of the data are used for estimating volume of selective activation in different age groups and visualization purposes, but not to evaluate statistical significance. One session per child per age group is included in the boxplot. To confirm the validity of the grouping of the age groups for the boxplots in Fig. 1D (5-9yo and 13-17yo), we used the LMMs (shown in Fig. 1C) to predict the size of category-selective activation for the mean age in years of the participants in each of the two age groups. This predicted size is indicated as a black diamond in the boxplots. Estimated mean size (diamonds) from the LMM corresponded well with the medians of the boxplots, thus, validating the grouping of the participants in the boxplots.

To estimate changes in the size of category-selective ROIs (Fig. 2D, Fig. S7) we used LMMs specified as: ROI size in mm^3^ ~ age in months + (1 | participant). Similarly, to estimate changes in the surface area of category-selective ROIs (Fig. S8) we used LMMs specified as: ROI surface area in mm^2^ ~ age in months + (1 | participant).

To evaluate significance of the developmental changes in emerging and waning ROIs, we used two separate LMMs: (i) selectivity changes across development (Fig. 3-left) were modeled as t-value ~ age in months + (1| participant), and (ii) changes in responses across development (Fig. 3-right) as % signal (b)~ age in months + (1 | participant). Percent signal refers to the change in response for a certain condition relative to baseline. Selectivity refers to contrasting responses to different categories (see above definition of contrasts and thresholds). Selectivity and response values were obtained for each voxel and LMMs were fit on the average value in an ROI. Boxplots in Fig. 3 show mean responses for each of the 10 categories in 5-9-year-olds and 13-17-year-olds. They are used for visualizing response amplitudes (units of % signal) in different age groups, but not to evaluate statistical significance, which was done using the LMM. The selection of age groups in the boxplots is validated by plotting the LMM prediction for the mean age in years of participants in each age group (compare black diamonds to measured median).

LMM analyses related to Fig. 4 tested if changes in limb-, face-, and word-selectivity in the developing ROIs were related to each other. For each emerging and waning ROI, we tested if selectivity for one category (e.g., word-selectivity in the emerging pOTS-words) was predicted by selectivity to the other categories (e.g., limb- and face-selectivity in the emerging pOTS-words). We also tested if an LMM with two predictors (e.g., predicting word-selectivity from both limb- and face-selectivity) is a better model than an LMM with one predictor (e.g., predicting word-selectivity just from face-selectivity) using a likelihood ratio test. If the likelihood ratio test confirmed that both predictors contributed significantly to the model fit, both predictors were included in the analysis. Parameters of LMMs and likelihood ratio tests are reported in supplemental Tables S10-11. Tables are grouped by analysis type.

The reported statistical tests are two-tailed. False-discovery rate (FDR) correction following the procedure by Benjamini and Hochberg^20^ as implemented in MATLAB version 2017b (The MathWorks, Inc.) was applied to correct for multiple comparisons related to the same analysis.

## Supporting information

corticalRecycling_supplements

## Data availability

The data that support the findings of this study are available from the corresponding author upon request.

## Code availability

Code will be made available with publication at: https://github.com/VPNL/Recycling.

## Acknowledgements

We thank Laura Villalobos, Erica Yeawon, Hwang, Savana Huskins, Alema Fitisemanu, and Philip Eykamp for manually editing gray-white matter brain segmentations. We thank Caitlyn Estrada and Nancy Lopez-Alvarez for help with data collection, and Rachel Hinds for help with data entry and management. We thank Jon Winawer for his constructive review of our manuscript.

## Funding

This work was supported by a fellowship of the German National Academic Foundation awarded to MN (NO 1448/1-1); NIH grant 2RO1 EY 022318 to KGS. NIH training grant 5T32EY020485 (VN); NSF Graduate Research Development Program (DGE-114747) and Ruth L. Kirschstein National Research Service Award (F31EY027201) to JG. The funders had no role in study design, data collection and analysis, decision to publish or preparation of the manuscript.

## Author contributions

M.N. collected data, developed and coded the analysis pipeline, analysed the data, and wrote the manuscript. V.N. and J.G. designed the experiment, collected data, and contributed to the manuscript. A.A.R. collected the data, contributed to data analysis, and contributed to the manuscript. D.F. and H.K, collected the data, and contributed to the manuscript. KGS designed the experiment, contributed to the analysis pipeline and data analysis, and wrote the manuscript.

## Competing interests

The authors declare no competing interests.

## Notes

### Competing Interest Statement

The authors have declared no competing interest.

## References

1. Kanwisher, N., McDermott, J. & Chun, M. M. The fusiform face area: a module in human extrastriate cortex specialized for face perception. J Neurosci 17, 4302–4311 (1997).

2. Peelen, M. V. & Downing, P. E. Selectivity for the human body in the fusiform gyrus. J. Neurophysiol. 93, 603–608 (2005).

3. Epstein, R. & Kanwisher, N. A cortical representation of the local visual environment. Nature 392, 598–601 (1998).

4. Cohen, L. et al. The visual word form area. Spatial and temporal characterization of an initial stage of reading in normal subjects and posterior split-brain patients. Brain 123, 291–307 (2000).

5. Deen, B. et al. Organization of high-level visual cortex in human infants. Nat. Commun. 8, 13995 (2017).

6. de Heering, A. & Rossion, B. Rapid categorization of natural face images in the infant right hemisphere. Elife 4, 1–14 (2015).

7. Livingstone, M. S. et al. Development of the macaque face-patch system. Nat. Commun. 8, 1–12 (2017).

8. Golarai, G. et al. Differential development of high-level visual cortex correlates with category-specific recognition memory. Nat. Neurosci. 10, 512–522 (2007).

9. Dehaene-Lambertz, G., Monzalvo, K. & Dehaene, S. The emergence of the visual word form: Longitudinal evolution of category-specific ventral visual areas during reading acquisition. PLoS Biol. 16, 1–34 (2018).

10. Srihasam, K., Vincent, J. L. & Livingstone, M. S. Novel domain formation reveals proto-architecture in inferotemporal cortex. Nat. Neurosci. 17, 1776–1783 (2014).

11. Gomez, J., Natu, V., Jeska, B., Barnett, M. & Grill-Spector, K. Development differentially sculpts receptive fields across early and high-level human visual cortex. Nat. Commun. 9, 1–12 (2018).

12. Nordt, M. et al. Learning to Read Increases the Informativeness of Distributed Ventral Temporal Responses. Cereb. Cortex 1–16 (2019). doi:10.1093/cercor/bhy178

13. Behrmann, M. & Plaut, D. C. A vision of graded hemispheric specialization. Ann. N. Y. Acad. Sci. 1359, 30–46 (2015).

14. Dehaene, S., Cohen, L., Morais, J. & Kolinsky, R. Illiterate to literate: behavioural and cerebral changes induced by reading acquisition. Nat. Rev. Neurosci. 16, 234–244 (2015).

15. Levy, I., Hasson, U., Avidan, G., Hendler, T. & Malach, R. Center-periphery organization of human object areas. Nat Neurosci 4, 533–539 (2001).

16. Cantlon, J. F., Pinel, P., Dehaene, S. & Pelphrey, K. A. Cortical representations of symbols, objects, and faces are pruned back during early childhood. Cereb. Cortex 21, 191–199 (2011).

17. Dehaene, S. et al. How Learning to Read Changes and Language. Science 1359, 1359–64 (2010).

18. Frost, M. A. & Goebel, R. Measuring structural-functional correspondence: Spatial variability of specialised brain regions after macro-anatomical alignment. Neuroimage 59, 1369–1381 (2012).

19. Weiner, K. S. et al. The mid-fusiform sulcus: A landmark identifying both cytoarchitectonic and functional divisions of human ventral temporal cortex. Neuroimage 84, 453–465 (2014).

20. Benjamini, Y. & Hochberg, Y. Controlling the False Discovery Rate: A Practical and Powerful Approach to Multiple Testing. J. R. Stat. Soc. Ser. B 57, 289–3300 (1995).

21. Scherf, K. S., Behrmann, M., Humphreys, K. & Luna, B. Visual category-selectivity for faces, places and objects emerges along different developmental trajectories. Dev. Sci. 10, (2007).

22. Peelen, M. V., Glaser, B., Vuilleumier, P. & Eliez, S. Differential development of selectivity for faces and bodies in the fusiform gyrus. Dev. Sci. 12, 16–25 (2009).

23. Ben-Shachar, M. The development of cortical sensitivity to visual word forms-Supplementary Information. 2387–2399 (2011).

24. Golarai, G., Liberman, A., Yoon, J. M. & Grill-Spector, K. Differential development of the ventral visual cortex extends through adolescence. Front. Hum. Neurosci. 3, 80 (2010).

25. Kobatake, E., Wang, G. & Tanaka, K. Effects of shape-discrimination training on the selectivity of inferotemporal cells in adult monkeys. J. Neurophysiol. 80, 324–330 (1998).

26. Arcaro, M. J., Schade, P. F., Vincent, J. L., Ponce, C. R. & Livingstone, M. S. Seeing faces is necessary for face-domain formation. Nat. Neurosci. 20, 1404–1412 (2017).

27. Fausey, C. M., Jayaraman, S. & Smith, L. B. From faces to hands: Changing visual input in the first two years. Cognition 152, 101–107 (2016).

28. Frank, M. C., Vul, E. & Saxe, R. Measuring the Development of Social Attention Using Free-Viewing. Infancy 17, 355–375 (2012).

29. Long, B., Kachergis, G., Agrawal, K., & Frank, M. C. Detecting social information in a dense database of infants’ natural visual experience. (2020).

30. Liszkowski, U., Carpenter, M. & Tomasello, M. Pointing out new news, old news, and absent referents at 12 months of age. Dev. Sci. 10, 1–7 (2007).

31. Haber, N., Mrowca, D., Wang, S., Fei-Fei, L. & Yamins, D. L. K. Learning to play with intrinsically-motivated, self-aware agents. Adv. Neural Inf. Process. Syst. 2018-Decem, 8388–8399 (2018).

32. Khaligh-Razavi, S. M. & Kriegeskorte, N. Deep Supervised, but Not Unsupervised, Models May Explain IT Cortical Representation. PLoS Comput. Biol. 10, (2014).

33. Zhuang, Chengxu; Zhai, Alex Lin; Yamins, D. Local Aggregation for Unsupervised Learning of Visual Embeddings. in Proceedings of the IEEE/CVF International Conference on Computer Vision (2019).

34. Gomez, J. et al. Microstructural proliferation in human cortex is coupled with the development of face processing. Science (80-.). 355, (2017).

35. Natu, V. S. et al. Development of Neural Sensitivity to Face Identity Correlates with Perceptual Discriminability. J. Neurosci. 36, 10893–10907 (2016).

36. Wandell, B. A., Rauschecker, A. M. & Yeatman, J. D. Learning to see words. Annu. Rev. Psychol. 63, 31–53 (2012).

37. Feldstein Ewing, S. W., Bjork, J. M. & Luciana, M. Implications of the ABCD study for developmental neuroscience. Dev. Cogn. Neurosci. 32, 161–164 (2018).

38. Constantino, J. N. et al. Infant viewing of social scenes is under genetic control and is atypical in autism. Nature 547, 340–344 (2017).

39. Duchaine, B. C. & Nakayama, K. Developmental prosopagnosia: a window to content-specific face processing. Curr. Opin. Neurobiol. 16, 166–173 (2006).

40. Amedi, A., Raz, N., Pianka, P., Malach, R. & Zohary, E. Early ‘visual’ cortex activation correlates with superior verbal memory performance in the blind. Nat. Neurosci. 6, 758–766 (2003).

41. Liu, T. T. et al. Successful Reorganization of Category-Selective Visual Cortex following Occipito-temporal Lobectomy in Childhood. Cell Rep. 24, 1113–1122.e6 (2018).

42. Norton, E. S., Beach, S. D. & Gabrieli, J. D. E. Neurobiology of dyslexia. Curr. Opin. Neurobiol. 30, 73–78 (2015).

43. Srihasam, K., Mandeville, J. B., Morocz, I. A., Sullivan, K. J. & Livingstone, M. S. Behavioral and Anatomical Consequences of Early versus Late Symbol Training in Macaques. Neuron 73, 608–619 (2012).

44. Büchel, C., Price, C. & Friston, K. A multimodal language region in the ventral visual pathway. Nature 394, 274–277 (1998).

45. Emmorey, K., McCullough, S. & Weisberg, J. Neural correlates of fingerspelling, text, and sign processing in deaf American sign language–English bilinguals. Lang. Cogn. Neurosci. 30, 749–767 (2015).

46. Bi, Y., Wang, X. & Caramazza, A. Object Domain and Modality in the Ventral Visual Pathway. Trends Cogn. Sci. 20, 282–290 (2016).

47. Reich, L., Szwed, M., Cohen, L. & Amedi, A. A ventral visual stream reading center independent of visual experience. Curr. Biol. 21, 363–368 (2011).

48. Dehaene, S. & Cohen, L. Cultural recycling of cortical maps. Neuron 56, 384–398 (2007).

49. Lucas, T. H., McKhann, G. M. & Ojemann, G. A. Functional separation of languages in the bilingual brain: A comparison of electrical stimulation language mapping in 25 bilingual patients and 117 monolingual control patients. J. Neurosurg. 101, 449–457 (2004).

50. Green, D. W., Crinion, J. & Price, C. J. Convergence, Degeneracy, and Control. Lang. Learn. 56, 99–125 (2006).

51. Amalric, M. & Dehaene, S. Origins of the brain networks for advanced mathematics in expert mathematicians. Proc. Natl. Acad. Sci. 113, 4909 LP–4917 (2016).

52. Kersey, A. J. & Cantlon, J. F. Neural Tuning to Numerosity Relates to Perceptual Tuning in 3-6-Year-Old Children. J. Neurosci. 37, 512 LP–522 (2017).

53. Pica, P., Lemer, C., Izard, V. & Dehaene, S. Exact and approximate arithmetic in an Amazonian indigene group. Science (80-.). 306, 499–503 (2004).

54. Natu, V. S. et al. Apparent thinning of human visual cortex during childhood is associated with myelination. Proc. Natl. Acad. Sci. U. S. A. 116, 20750–20759 (2019).

55. Mezer, A. et al. Quantifying the local tissue volume and composition in individual brains with magnetic resonance imaging. Nat. Med. 19, 1667–1672 (2013).

56. Grill-Spector, K. & Kanwisher, N. Visual recognition: as soon as you know it is there, you know what it is. Psychol Sci 16, 152–160 (2005).

57. Weiner, K. S. & Grill-Spector, K. Sparsely-distributed organization of face and limb activations in human ventral temporal cortex. Neuroimage 52, 1559–1573 (2010).

58. Weiner, K. S., Sayres, R., Vinberg, J. & Grill-Spector, K. fMRI-adaptation and category selectivity in human ventral temporal cortex: Regional differences across time scales. J. Neurophysiol. 103, 3349–3365 (2010).

59. Stigliani, A., Weiner, K. S. & Grill-Spector, K. Temporal Processing Capacity in High-Level Visual Cortex Is Domain Specific. J. Neurosci. 35, 12412–12424 (2015).

60. Kanwisher, N. Domain specificity in face perception. Nat. Neurosci. 3, 759–763 (2000).

61. Reuter, M., Schmansky, N. J., Rosas, H. D. & Fischl, B. Within-subject template estimation for unbiased longitudinal image analysis. Neuroimage 61, 1402–1418 (2012).

62. Haxby, J. V et al. Distributed and overlapping representations of faces and objects in ventral temporal cortex. Science (80-.). 293, 2425–2430 (2001).

63. Glasser, M. F. et al. A multi-modal parcellation of human cerebral cortex. Nature 536, 171–178 (2016).

