## Supplementary material for "Cortical recycling in high-level visual cortex during childhood development": corticalRecycling_supplements

### Supplementary Figures

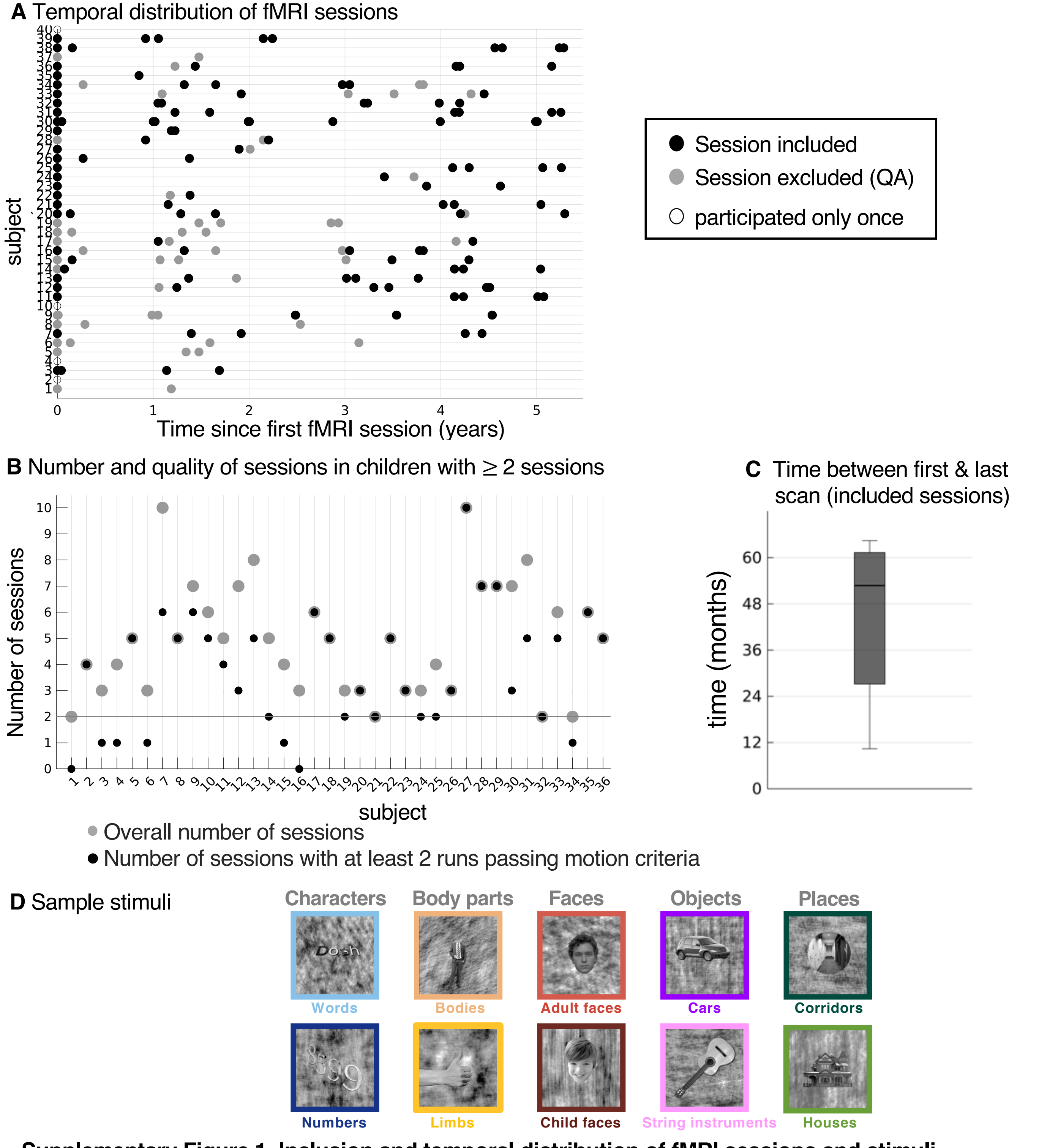


**Supplementary Figure 1. Inclusion and temporal distribution of fMRI sessions and stimuli**
(**A**) The temporal distribution of fMRI sessions is plotted relative to each subject's first session. *Filled black dots*: included sessions. *Filled gray dots:* sessions that were excluded due to quality assurance (QA). *Black outlined dots:* sessions of children who participated only once. (**B**) Overview of data from children who participated more than once. *Gray dots*: number of total sessions per child. Not included are 2 sessions which were excluded due to technical error. *Black dots*: number of sessions passing motion criteria. *Horizontal black line:* cut-off for inclusion in the present study (>=2 sessions passing motion criteria). (**C**) The boxplot shows the time between the first and last included fMRI session in all included subjects (n=29). (**D**) Sample stimuli for the 10 categories which can be grouped into five domains. Each domain comprises two categories: characters (words, numbers), body parts (headless bodies, limbs), faces (adult faces, child faces), objects (cars, string instruments), and places (corridors, houses).

**A** Lateral VTC

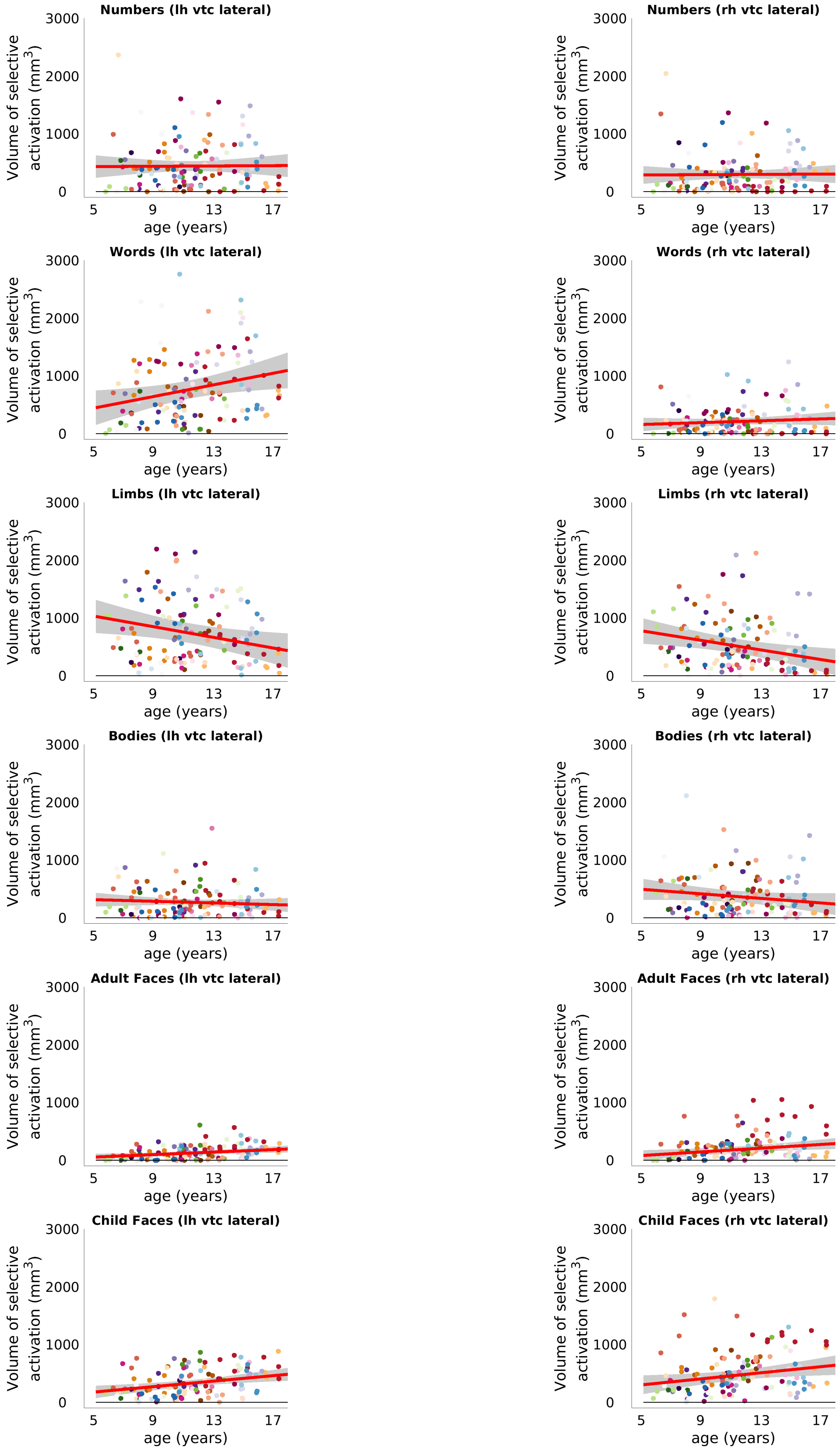

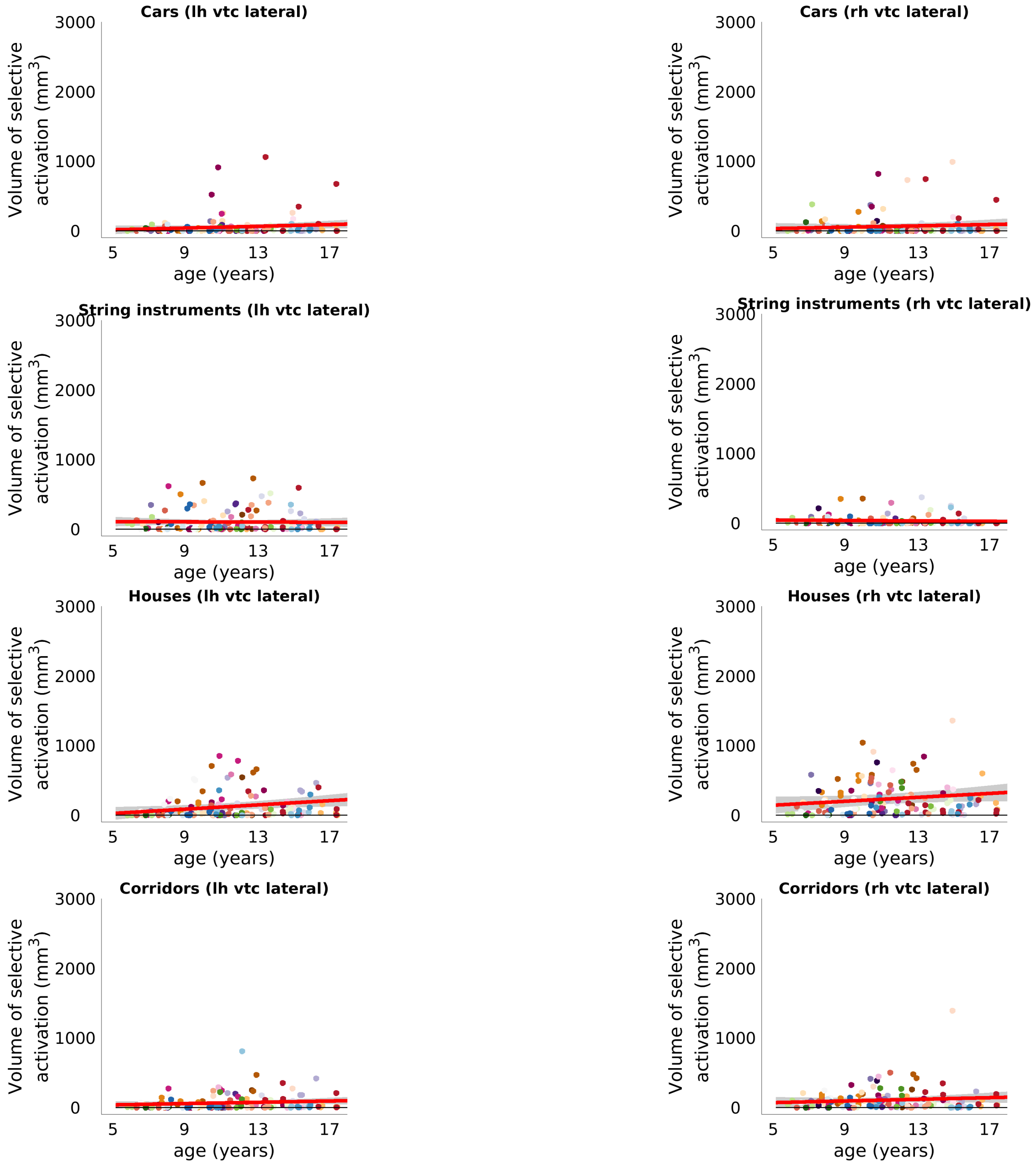
Cars (lh vtc lateral)
Cars (rh vtc lateral)
String instruments (lh vtc lateral)
String instruments (rh vtc lateral)
Houses (lh vtc lateral)
Houses (rh vtc lateral)
Corridors (lh vtc lateral)
Corridors (rh vtc lateral)
Volume of selective activation (mm$^3$)
age (years)
3000
2000
1000
0
5
9
13
17

**B** Medial VTC

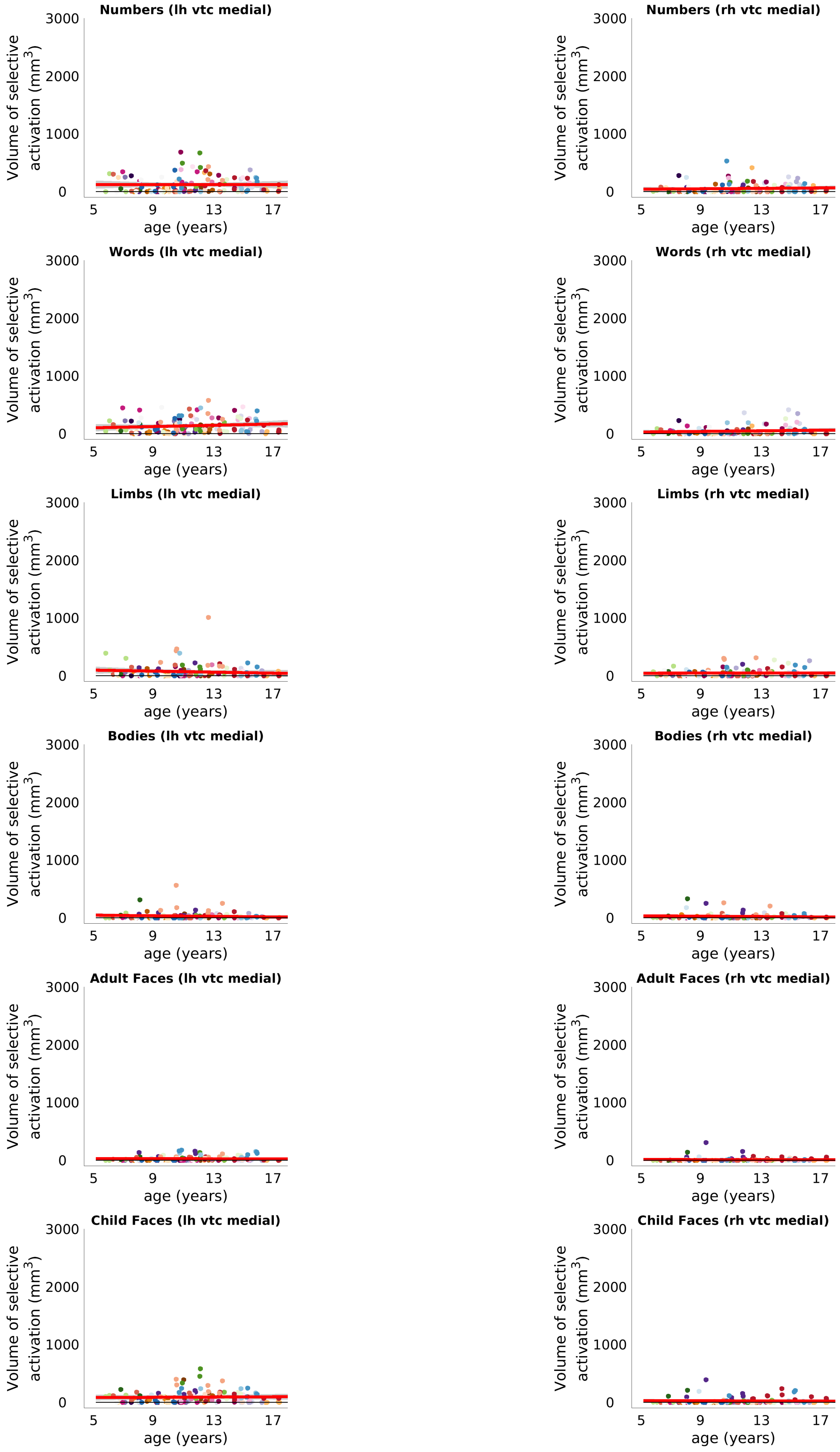
Numbers (lh vtc medial)
Numbers (rh vtc medial)
Words (lh vtc medial)
Words (rh vtc medial)
Limbs (lh vtc medial)
Limbs (rh vtc medial)
Bodies (lh vtc medial)
Bodies (rh vtc medial)
Adult Faces (lh vtc medial)
Adult Faces (rh vtc medial)
Child Faces (lh vtc medial)
Child Faces (rh vtc medial)
Volume of selective activation (mm$^3$)
0
1000
2000
3000
age (years)
5
9
13
17

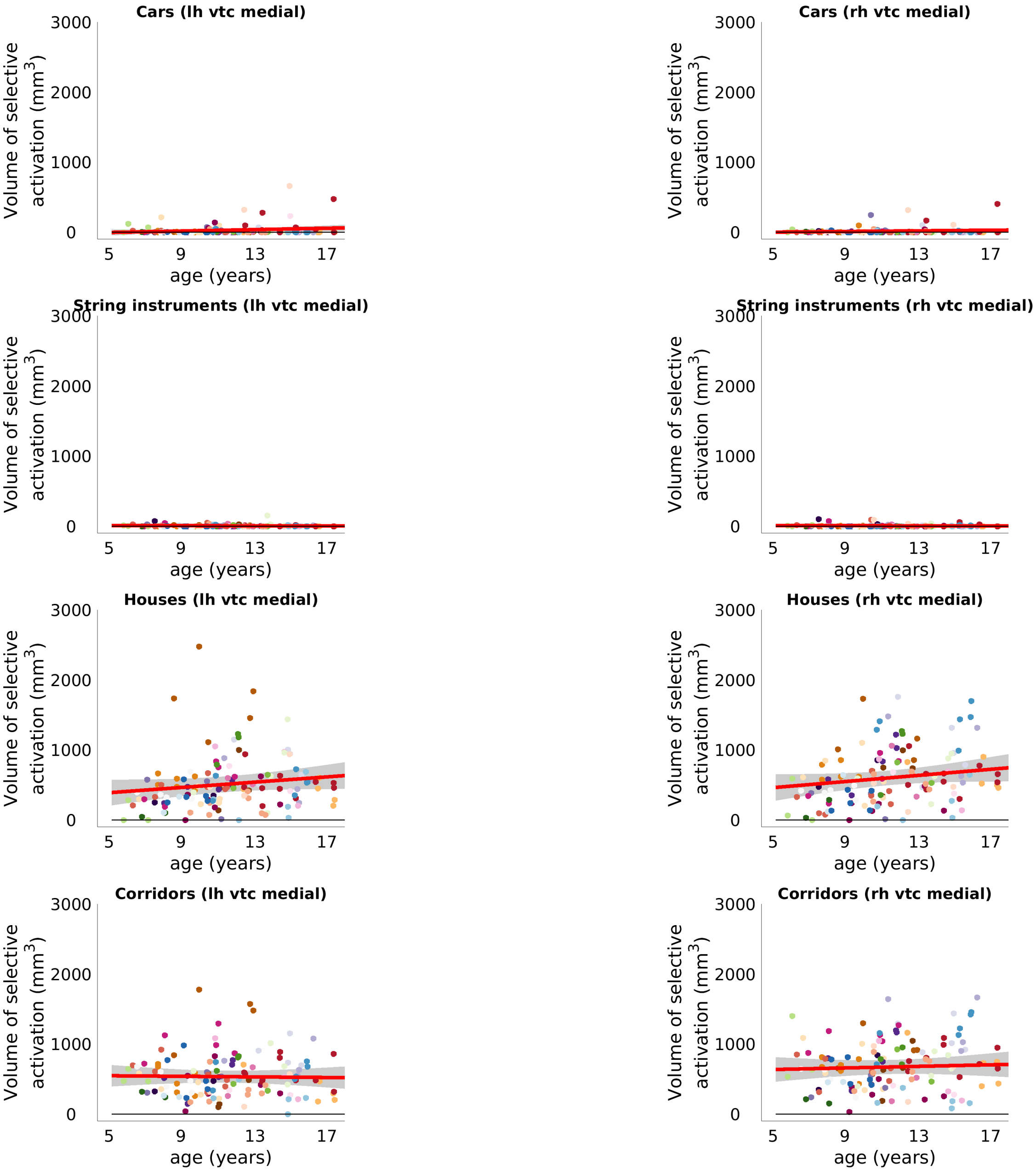


**Supplementary Figure 2. Development of the volume of selective activation for each category in lateral and medial VTC.** Each dot represents one session. Sessions belonging to the same subject are coded by color. The red line indicates the LMM predicting volume of selective activation by age. The gray shaded area represents the 95% CI. (**A**) Lateral VTC. Categories showing significant development are: words (left), limbs (bilateral), adult faces (bilateral), child faces (bilateral) and houses (left). Statistics are reported in Table S1. Note, as examination of activations in individual subjects did not reveal consistent clusters of activation for houses in left lateral VTC in contrast to those for faces, limbs or words – no house ROI was included in follow-up analyses. (**B**) Medial VTC. There were no significant effects in medial VTC. Statistics are reported in Table S2. Related to Fig. 1.

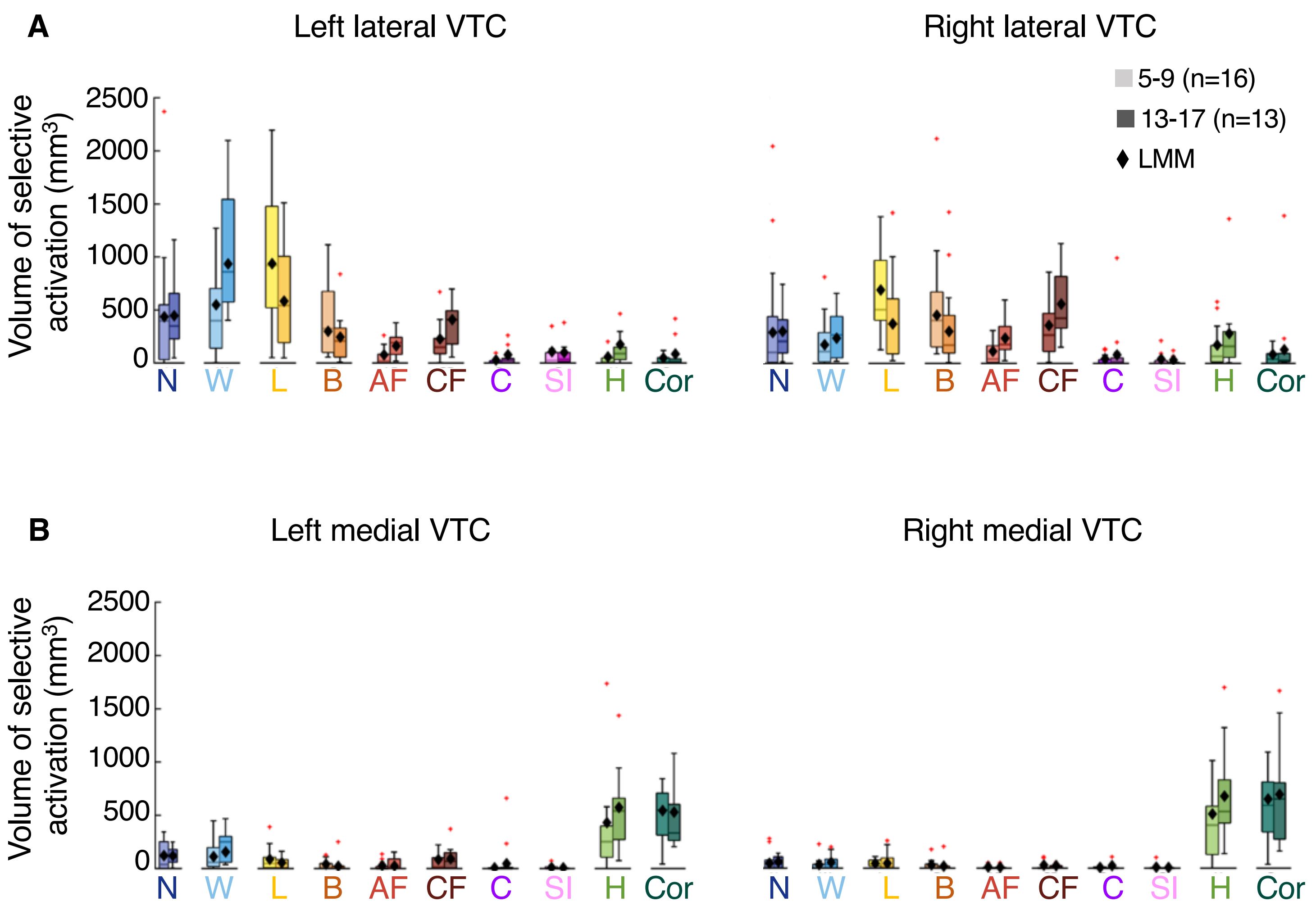


**Supplementary Figure 3. Changes in category-selective activation in lateral and medial VTC.** (**A**) Boxplots showing the volume of selective activation for all 10 categories for 5-9-year-olds (n=16) and 13-17-year-olds (n=13) in left and right lateral VTC. One functional session per child is included per boxplot. *Black diamonds*: LMM prediction for the mean age of each age group. (**B**) Same as (A) but for medial VTC. N = numbers; W = words; L = limbs; B = bodies; AF = adult faces; CF = child faces; C = cars; SI = string instruments; H = houses; Cor = corridors. Related to Fig 1D.

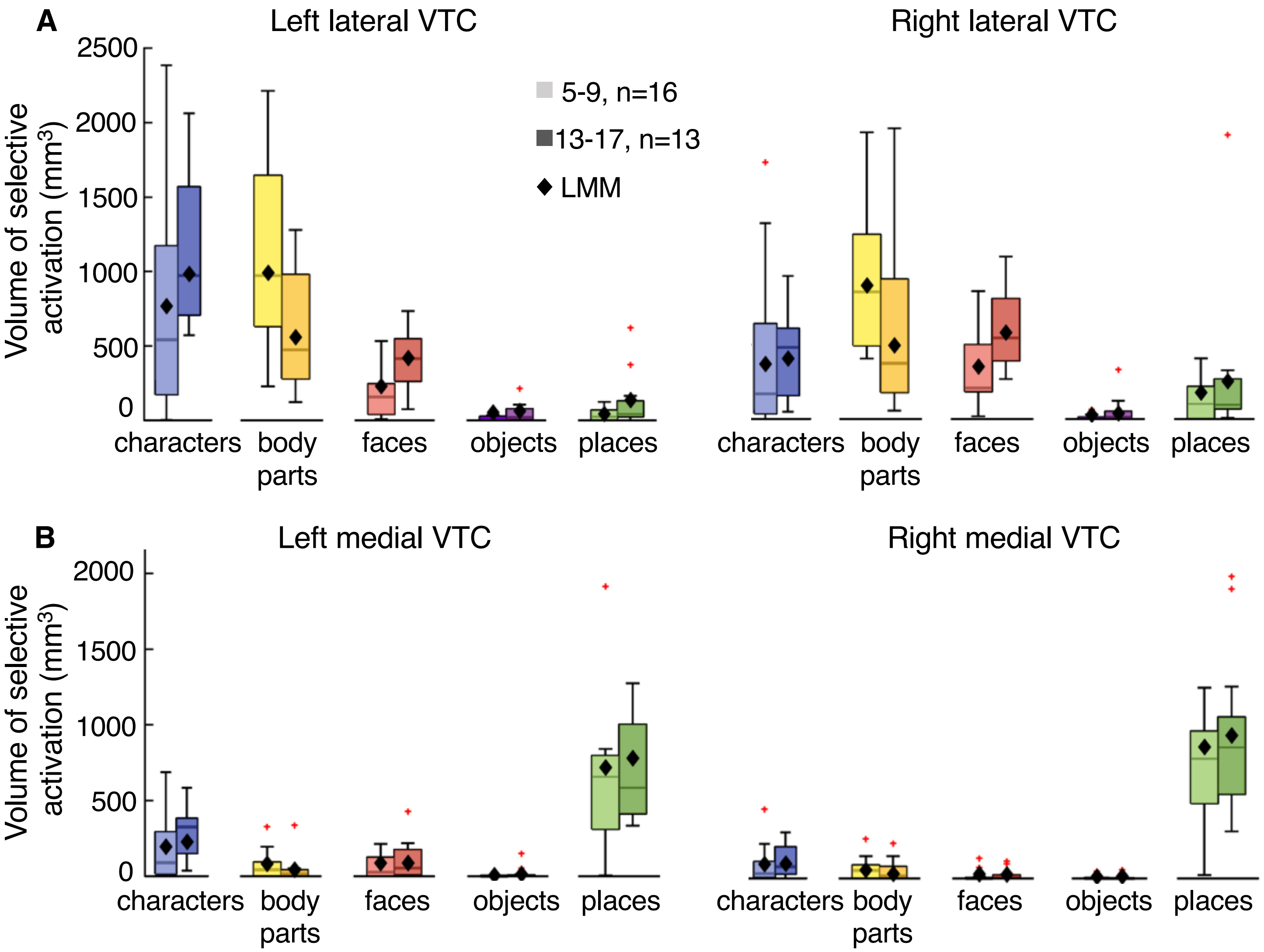


**Supplementary Figure 4. Changes in the volume of domain-selective activation in lateral and medial VTC.**
**(A)** Boxplots showing volume of selective activation for 5 domains in the left and right lateral VTC of 5-9-year-olds (n=16) and 13-17-year-olds (n=13) children. One functional session per child is included per boxplot. Lighter colors indicate younger ages. *Black diamonds*: LMM prediction for the mean age of each age group. For domain contrasts, each domain is contrasted with all other domains (selectivity: t-value>3, voxel-level). LMMs (n=128 sessions) reveal significant bilateral development for body parts and faces. Specifically, the volume of body part-selective activation decreased in the left (slope=-4.63 (CI:-7.6,-1.7) [$mm^3$/month], t(126)=-3.10, p=0.016, FDR-corrected) and right hemisphere (slope=-4.31 (CI:-6.9,-1.7) [$mm^3$/month], t(126)=-3.27, p=0.014, FDR-corrected), as volume of face-selective activation increased bilaterally (left: slope=2.03 (CI: 0.87,3.18) [$mm^3$/month], t(126)=3.48, p=0.014; right=2.45 (CI: 0.81,4.10) [$mm^3$/month], t(126)=2.95, p=0.019, FDR-corrected). No other domain showed significant development, but there was a trend for increase in volume of place-selective activation in the left hemisphere (slope=0.10 (CI: 0.21,1.78) [$mm^3$/month], t(126)=2.52, p=0.053, FDR-corrected). **(B)** Same as (A) but for medial VTC. No domain showed significant development in medial VTC. Related to Fig 1.

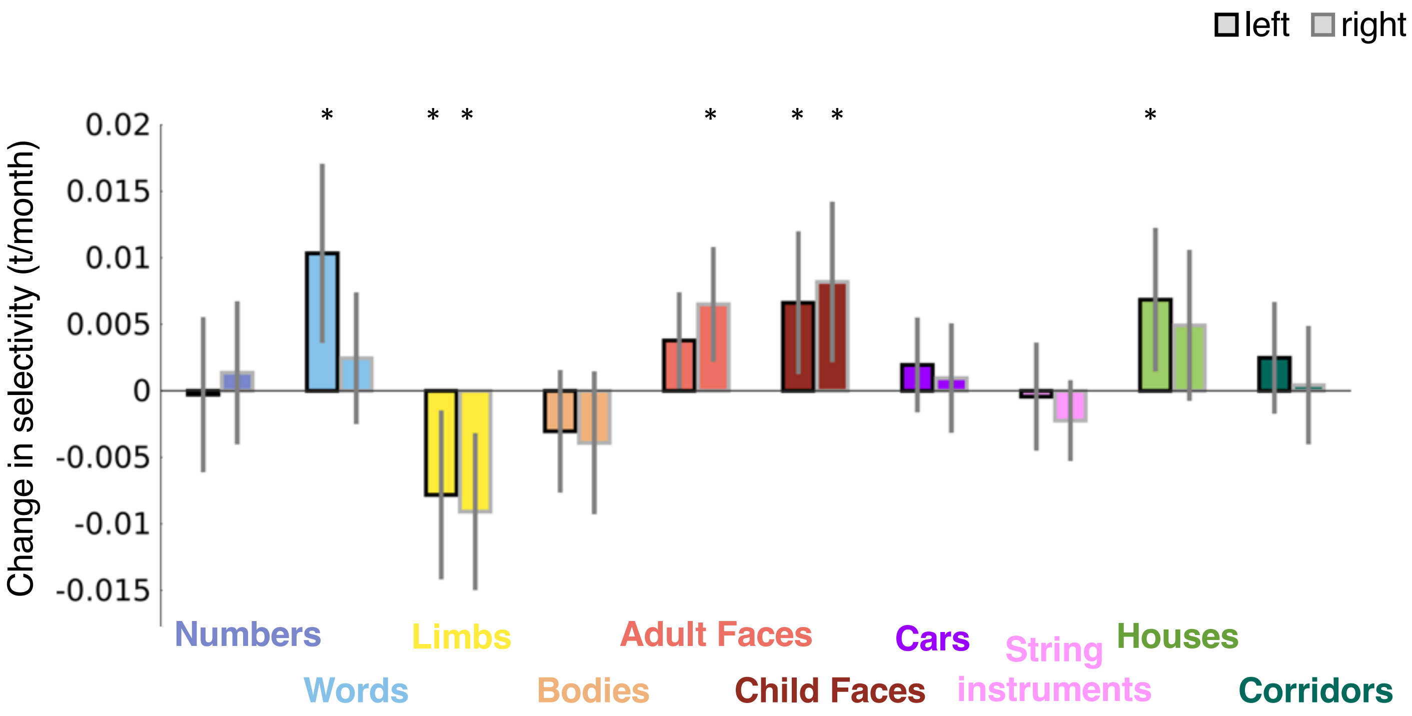


**Supplementary Figure 5. Development of selectivity in the 20% most selective voxels of lateral VTC.**

Slopes of LMMs indicate the change in selectivity (t-value/month) in the 20% most selective voxels for each category contrast in left and right lateral VTC (n=128 sessions, 29 children). Note that the number of included voxels (corresponding to the 20% most selective) per participant does not change across sessions, as all functional sessions of the same participant are aligned to the same brain anatomy (individual subject template).This analysis reveals significant developmental increases in selectivity for words in the left lateral VTC (left: slope= 0.010 (CI:0.004,0.017) (t/month), t(126)=3.04, $p_{FDR}$=0.023), a significant decrease for limbs in both hemispheres (left: slope=-0.008 (CI:-0.014,-0.002) (t/month), t(126)=-2.44, $p_{FDR}$=0.047; right=-0.009 (CI:-0.015,-0.003) (t/month), t(126)=-3.05, $p_{FDR}$=0.023), significant increases for adult faces in the right hemisphere (right: slope=0.007 (CI:0.002,0.011) (t/month), t(126)=2.98, $p_{FDR}$=0.023), child faces in both hemispheres (left: slope=0.007 (CI:0.0012,0.012) (t/month), t(126)=2.43, $p_{FDR}$=0.047; right: slope=0.008 (CI:0.002,0.014) (t/month), t(126)=2.68, $p_{FDR}$=0.041), and houses in the left lateral VTC (left: slope=0.007 (CI:0.0014,0.012) (t/month), t(126)=2.51, $p_{FDR}$=0.047). *Error bars:* 95% CI of the slope. *Asterisks*: effects surviving FDR correction ($p_{FDR}$<0.05). Related to Fig. 1C.

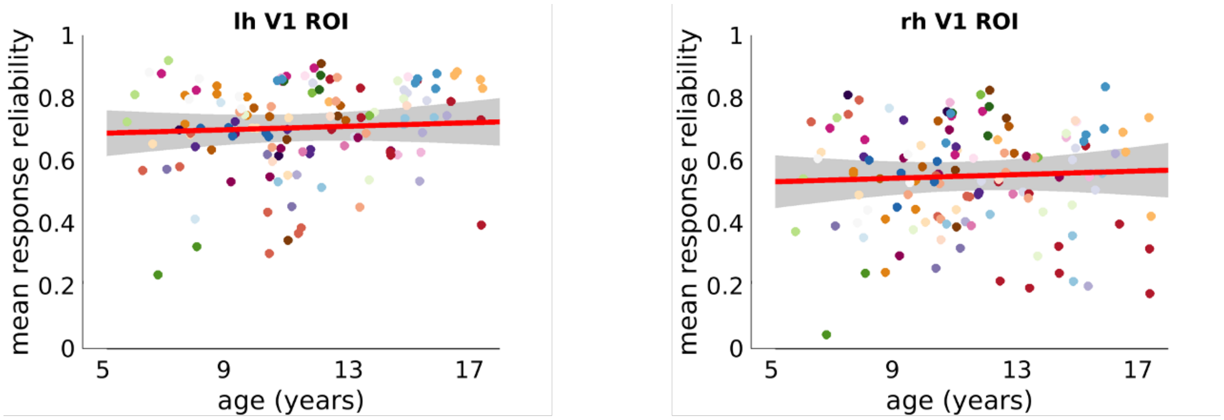


**Supplementary Figure 6. The reliability of V1 responses does not change with age**

We measured the reliability of distributed responses for items of the same categories across two runs in each session. We then calculated the mean response reliability as the mean of response reliability across all 10 categories. I.e., this is the mean of the on-diagonal of the representational similarity matrix for the 10 categories based on multivoxel patterns for the two runs included in each session (see Methods). Each dot is a session and dots are colored by participants.

Scatter plots show the response reliability in V1 by age. *Red line:* Linear mixed model (LMM) prediction of response reliability by age using a random intercept model with the grouping variable subject (response reliability ~ age +(1|subject)). *Shaded gray:* 95% confidence interval (CI). Analyses reveal that response reliability in V1 is independent of age as age is not a significant predictor (left: $\beta_{age}$=0.0002 (CI:-0.0005,0.001), $t(126)=0.59$, $p=0.56$; right: $\beta_{age}$=0.0002 (CI:-0.0007,0.001), $t(126)=0.51$, $p=0.61$).

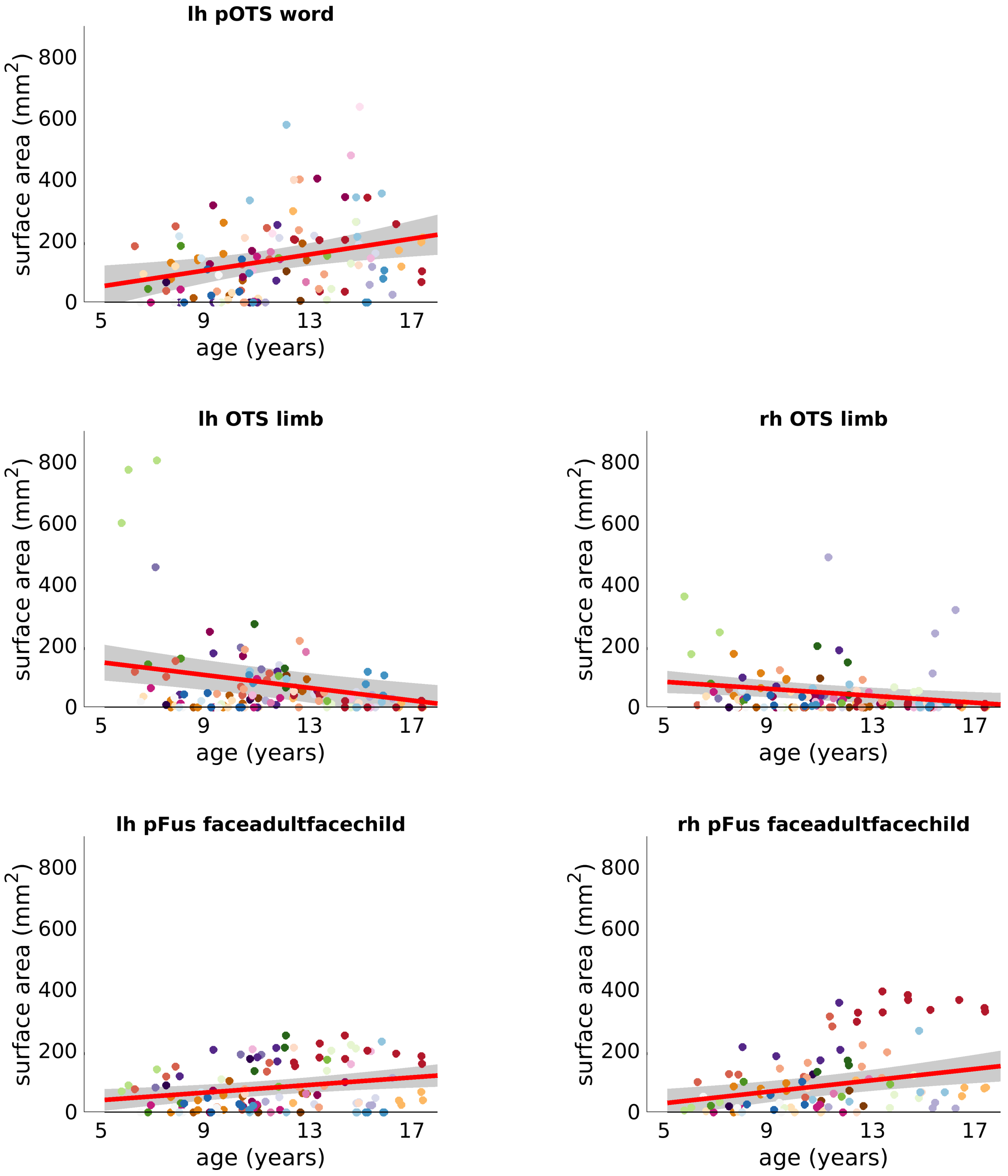


**Supplementary Figure 7. Differential development of ROI surface area.**

Surface area of word-, limb- and face-selective regions by age. Each dot is a session and dots are colored by participant. *Red line:* LMM prediction of surface area of ROIs by age. Shaded *gray:* 95% confidence interval (CI).

This analysis replicates the results presented in Fig. 2: There is a significant increase in the surface area of left pOTS-words ($\beta_{age}$=1.07 (CI:0.35,1.79), t(117)=2.94, p=0.004) and left and right pFus-faces (left: $\beta_{age}$=0.51 (CI:0.17,0.85), t(118)=2.94, p=0.004; right: $\beta_{age}$=0.77 (CI:0.32,1.22), t(96)=3.38, p=0.001), as well as a significant decrease in surface area of left and right OTS-limbs (left: $\beta_{age}$=-0.86 (CI:-1.32,-0.39), t(124)=-3.65, p=0.0004; right: $\beta_{age}$=-0.47 (CI:-0.81, -0.13), t(124)=-2.72, p=0.0075).

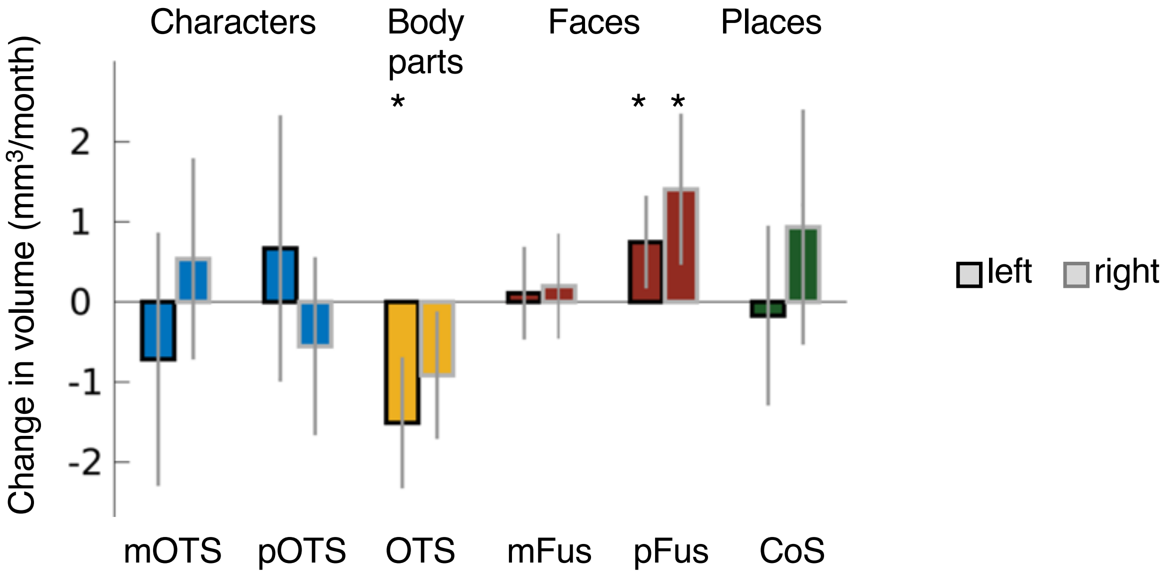


**Supplementary Figure 8. Differential development of domain-selective ROIs.**

Slopes of LMMs indicating the change in ROI volume for character-, body part-, and face- and place-selective regions when using domain-contrasts to define ROIs. Results of this analysis extend findings related to Fig. 2. They show that:

(i) The increase in word-selective volume in the left hemisphere is category-specific and does not extend to characters (words + numbers > others, slope=0.67 (CI:-0.99,2.33), $t(96)=0.80$, $p_{FDR}=0.57$).

(ii) The decrease in limb-selective activation extends to the combined contrast (limbs + headless bodies > others) in the left hemisphere (slope=-1.51 (CI:-2.33,-0.69) ($mm^3$/month), $t(124)=-3.65$, $p_{FDR}=0.0046$) and trends in the right hemisphere (slope=-0.91 (CI:-1.71,-0.12) ($mm^3$/month), $t(123)=-2.27$, $p_{FDR}=0.07$).

(iii) While mFus-faces does not show significant development, the posterior pFus-faces develops significantly in both hemispheres (left: 0.75 (CI:0.17,1.33) ($mm^3$/month), $t(118)=2.55$, $p_{FDR}=0.048$; right: slope=1.41 (CI:0.46,2.35) ($mm^3$/month), $t(96)=2.96$, $p_{FDR}=0.023$, note that for face-selective regions the same contrast was used in Fig. 2D).

(iv) Place-selective CoS-places does not develop significantly, replicating the non-significant increase of place-selective volume in medial VTC in Fig. 1C.

*Error bars*: 95% CI of the slope. *Asterisks:* effects surviving FDR correction ($p_{FDR}<0.05$). Related to Fig. 2.

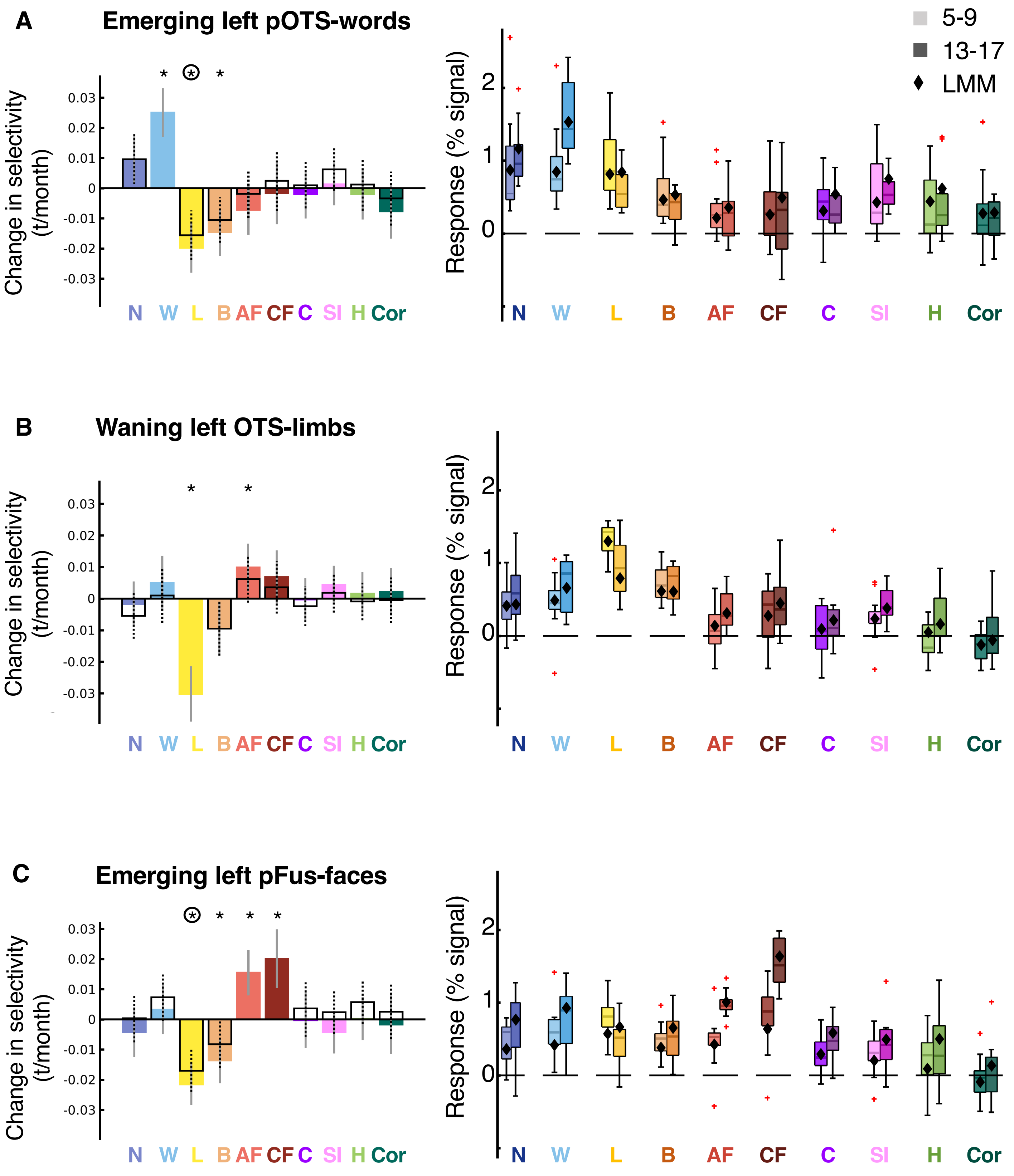

A
Emerging left pOTS-words
B
Waning left OTS-limbs
C
Emerging left pFus-faces
Change in selectivity (t/month)
Response (% signal)
5-9
13-17
LMM
N W L B AF CF C SI H Cor

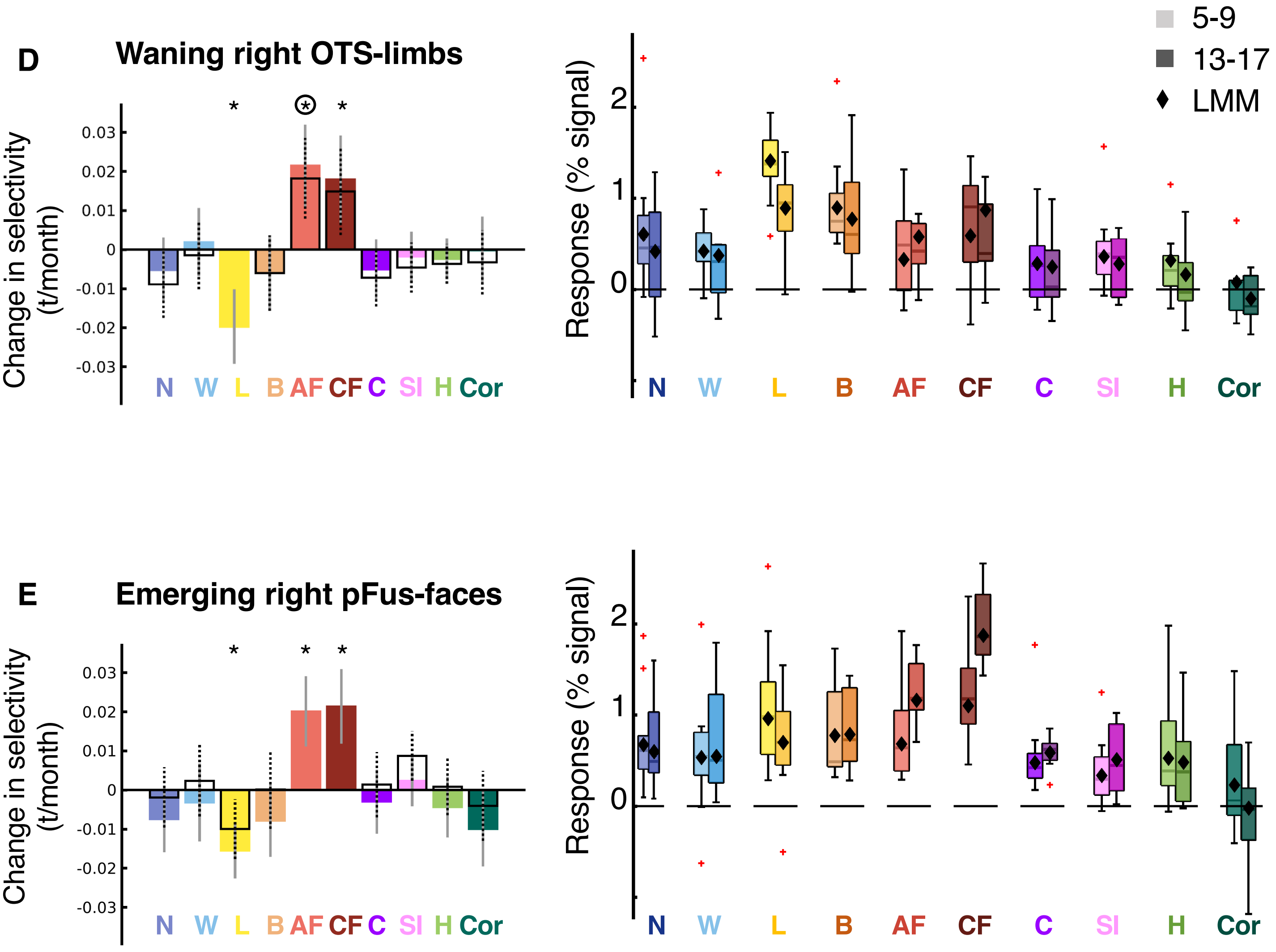


**Supplementary Figure 9. Functional changes underlying the development of category-selective ROIs.** *Left panel: Colored bars:* Slopes of LMMs indicating changes in selectivity by age for all 10 categories in emerging and waning ROIs. *Open bars:* LMM slopes for contrasts in which the ROI-defining category is not included. *Asterisks*: significant development after FDR-correction (p<0.05) for colored bars, *circles*: significant after FDR-correction for open bars. *Right panel:* Response amplitudes for 5-9-year-olds and 13-17-year-olds. Lighter colors indicate younger ages. One functional session per child is included per boxplot. *Black diamonds*: LMM prediction for the response at the mean age of each age group. *Red crosses:* outliers. (**A**) Left emerging pOTS-words. Left panel: n=24 (112 sessions). Right panel: 5-9-year-olds (n=12); 13-17-year-olds (n=13). (**B**) Left waning OTS-limbs. Left panel: n=26, 122 sessions. Right panel: 5-9-year-olds: n=15; 13-17-year-olds: n=12. (**C**) Left emerging pFus-faces. Left panel: n=22, 105 sessions. Right panel: 5-9-year-olds: n=13; 13-17-year-olds: n=11). (**D**) Right waning OTS-limbs. Left panel: n=21, 100 sessions. Right panel: 5-9-year-olds: n=12; 13-17-year-olds: n=10. (**E**) Right emerging pFus-faces. Left panel: n=21, 96 sessions. Right panel: 5-9-year-olds: n=12; 13-17-year-olds: n=8. As we observed a significant decrease in limb-selectivity in emerging parts of word- and face-selective regions and word-, face- and limb-selective regions neighbor, we tested if emerging parts of word- and face-selective regions overlap with waning parts of the limb-selective regions. However, the overlap between the developing parts of the ROIs (difference between initial and end ROIs) assessed by the dice coefficient (DC) was small (overlap between developing parts of OTS-limbs and pFus-faces, left: DC=0.025±0.01 (mean± SD), n=22; right: DC=0.026±0.01, n=18; overlap between developing parts of left OTS-limbs and pOTS-words: DC=0.006±SD=0.004, n=21). Related to Fig 3.

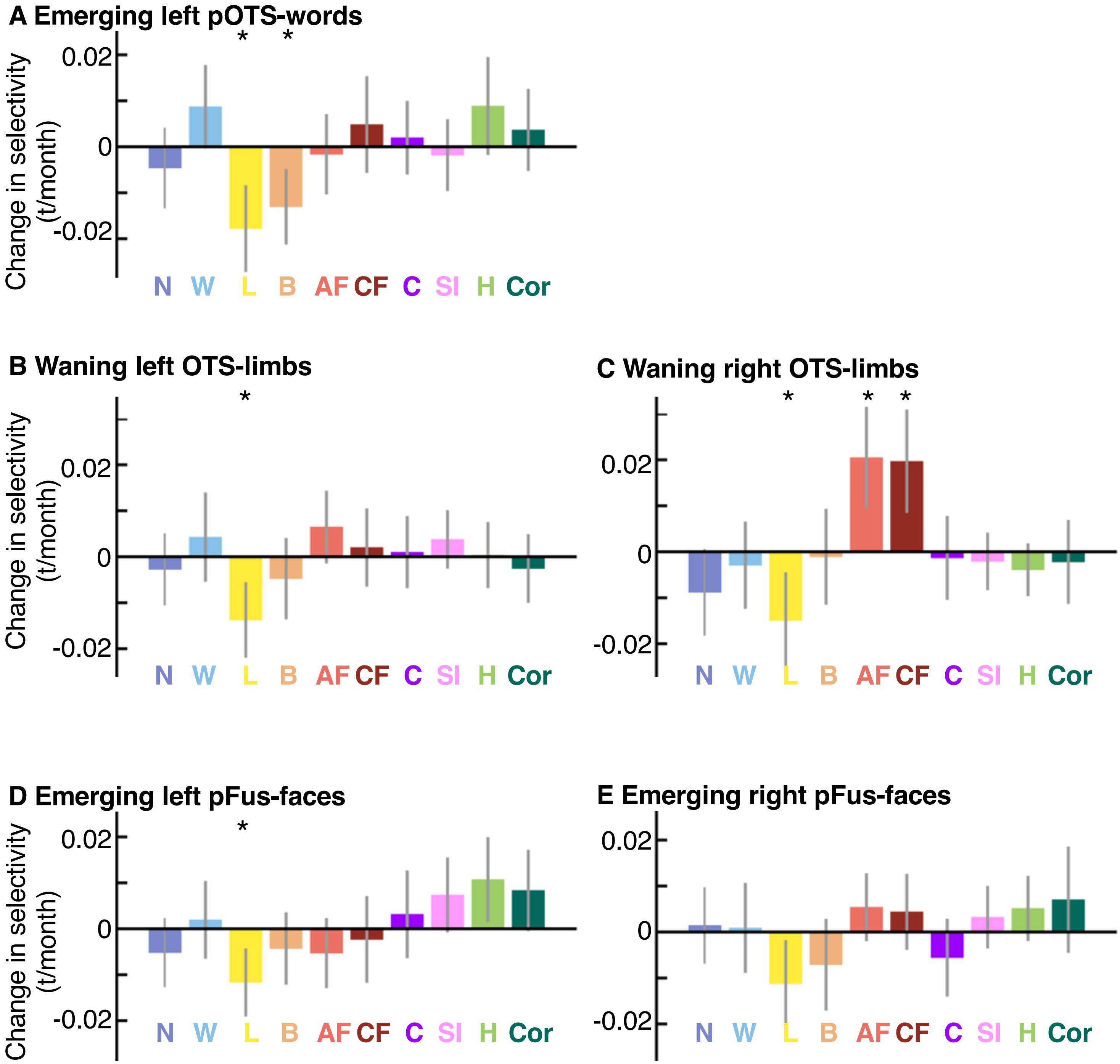


**Supplementary Figure 10. Development of selectivity in independent ring-shaped ROIs.**
Bar plots show slopes of LMMs for changes in selectivity by age in independent ring-shaped ROIs. (**A**) In the left pOTS-words-ring ROI there was a significant decrease for both limb- (slope=-0.018 (CI:-0.027,-0.008) [t/month], t(95)=-3.75, $p_{FDR}$=0.010) and body-selectivity (slope=-0.013 (CI:-0.02,-0.005) [t/month], t(95)=-3.16, $p_{FDR}$=0.019). (**B**) In the left OTS-limbs-ring ROI limb-selectivity decreased significantly (slope=-0.014 (CI:-0.02,-0.006) [t/month], t(118)=-3.32, $p_{FDR}$=0.015). (**C**) In the right OTS-limbs-ring ROI limb-selectivity decreased (slope=-0.015 (CI:-0.026,-0.004) [t/month], t(87)=-2.83, $p_{FDR}$=0.042) while face-selectivity increased (adult faces: slope=0.021 (CI:0.009,0.032) [t/month], t(87)=3.68, $p_{FDR}$=0.010; child faces: slope=0.02 (CI:0.008,0.031) [t/month], t(87)=3.48, $p_{FDR}$=0.013). (**D**) In the left pFus-faces-ring ROI selectivity to limbs decreased significantly (slope=-0.012 (CI:-0.019,-0.004) [t/month], t=-3.13, $p_{FDR}$=0.019). (**E**) In the right pFus-faces-ring ROI no effect survived FDR-correction. *Error bars:* 95% CI of the slopes. *Asterisks*: significant after FDR-correction ($p_{FDR}$<0.05). N = numbers; W = words; L = limbs; B = bodies; AF = adult faces; CF = child faces; C = cars; SI = string instruments; H = houses; Cor = corridors. Related to Fig. 3.

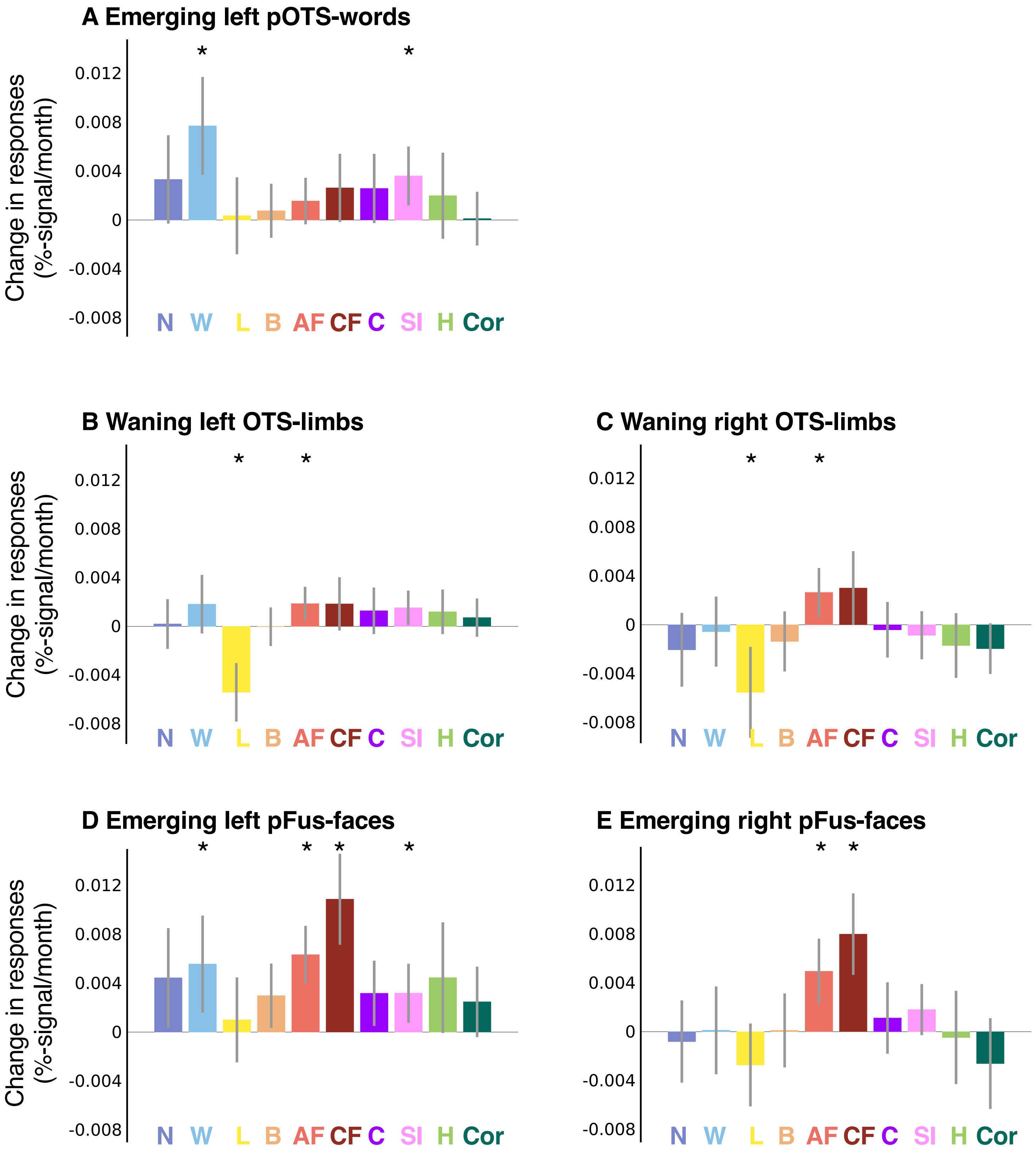


**Supplementary Figure 11. Development of response amplitudes in waning and emerging parts of ROIs.**

Bar plots show slopes of LMMs for changes in response amplitude (%-signal) by age. *Error bars:* 95% CI of the slopes. *Asterisks*: significant after FDR-correction ($p_{FDR}$<0.05). Statistics are reported in Table S9. N = numbers; W = words; L = limbs; B = bodies; AF = adult faces; CF = child faces; C = cars; SI = string instruments; H = houses; Cor = corridors. Related to Fig. 3.

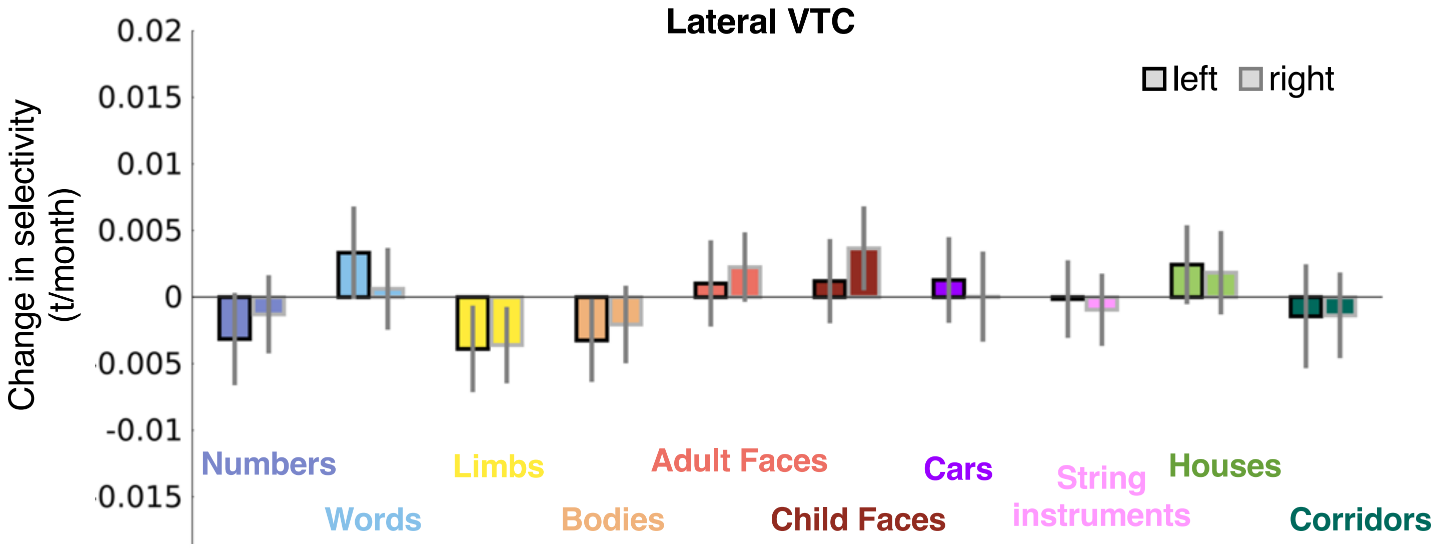


**Supplementary Figure 12. No significant development of limb-selectivity in the remaining lateral VTC after excluding the voxels selective for categories showing significant development**. Slopes of LMMs (n=128 sessions) indicating the change in selectivity (t/month) in the remainder of lateral VTC voxels after excluding the voxels that were selective to the categories showing development (adult faces, child faces, limbs, words, houses). No effects survive FDR-correction. *Error bars:* 95% CI of the slope. Related to Fig. 3.

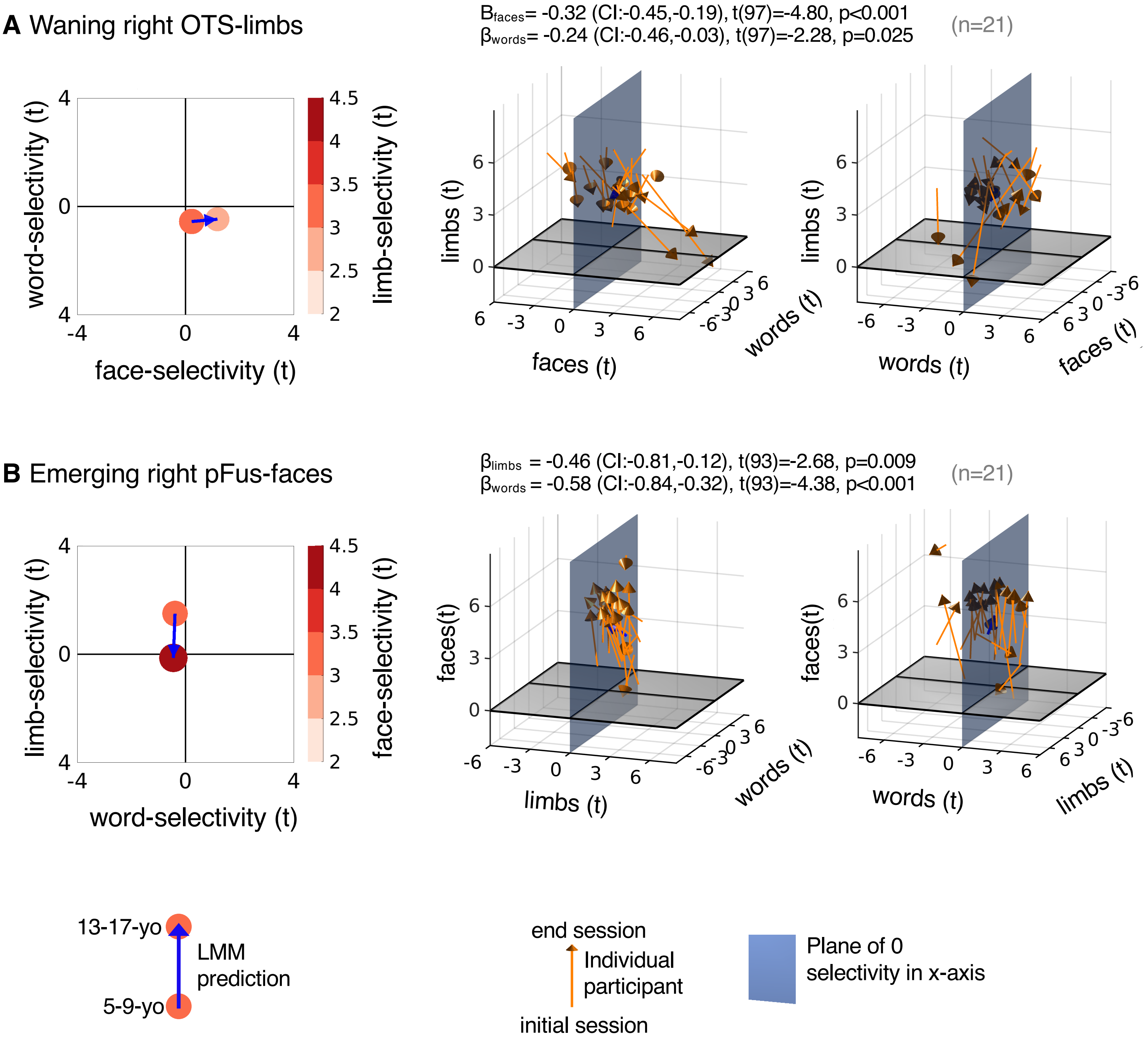


**Supplementary Figure 13. Developmental changes in word-, face-, and limb-selectivity are also linked in the right hemisphere.**
(**A**) Limb-selectivity vs face- and word-selectivity in the waning right OTS-limbs. (**B**) Face-selectivity vs. limb- and word-selectivity in the emerging right pFus faces. *Left:* Model prediction for 5-9-year-olds and 13-17-year-olds for the selectivity that defines the ROI as a function of the selectivity to the other two variables. *Middle:* Individual participant data visualized in 3D. In each panel the variable on the z-axis is related to the x- and y-variables. LMM $\beta s$, 95%-CIs, t-values, df, and p-values are shown on top. Full statistics are reported in Table S11. *Orange arrows:* Individual child data. *Blue arrows:* LMM, same as left panel. *Right:* Rotated version of the plots in the middle column to increase visibility of changes along the horizontal axes. Related to **Fig. 4.**

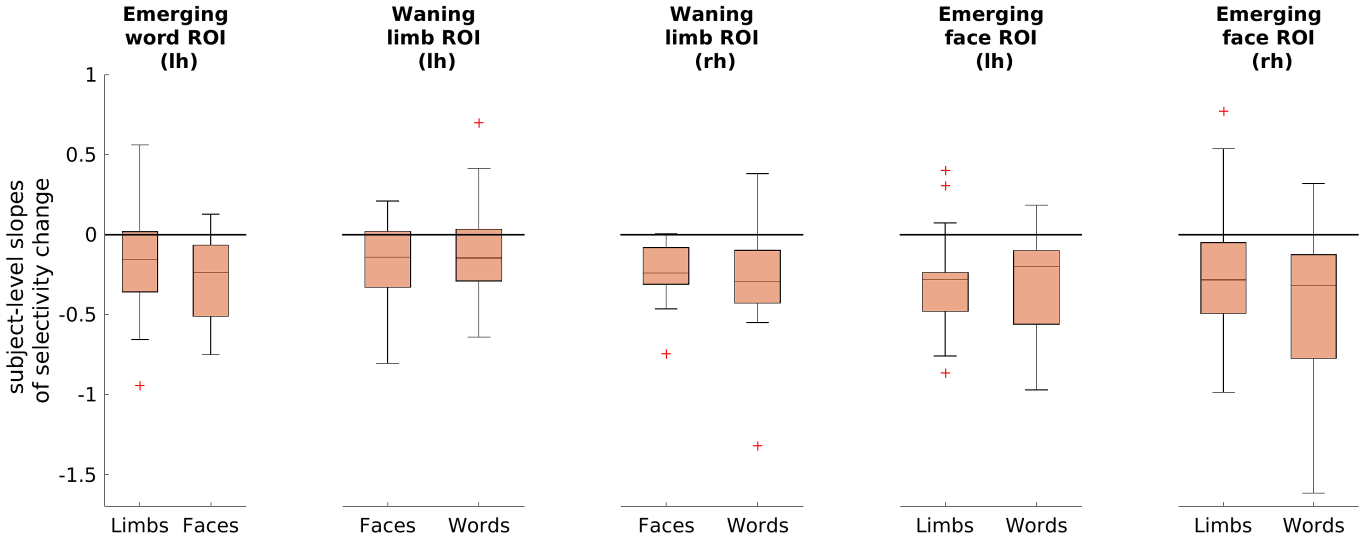


**Supplementary Figure 14. Link of developmental changes in word-, face-, and limb-selectivity on the voxel-level.** An interesting question is whether the observed developmental effects occur at a finer spatial scale. Thus, we tested if the link between development of face-, word, and limb-selectivity is also evident at the voxel-level (see Methods). We ran LMMs relating selectivity to one category as a function of the two other categories voxels in each participant's waning and emerging ROIs. LMMs were conducted separately for each participant and ROI and each point in the LMM is a voxel. Boxplots show the 75% and 25% percentiles (colored areas), median (horizontal lines) and range (whiskers) of individual subject LMM $\beta s$ based on voxel data in emerging and waning ROIs. Negative $\beta s$ indicate that increases in voxel selectivity to the category defining the ROI is related to decreases in voxel-selectivity to the other categories (and vice versa). *Red crosses*: outliers (values more than 1.5 times the interquartile range away from the bottom or top of the box). Results suggest that cortical recycling also occurs at the voxel level: we find an overall negative relationship between selectivity to the preferred category and the other two categories across voxels of the emerging and waning ROIs within individual subjects as the median of the individual subject LMM slopes is negative in all developing ROIs. While these effects are not significant in all individual participants, they suggest that the negative relationship between selectivity to the preferred category and the other two categories can also be measured at the voxel level. Related to Fig. 4.

**Supplementary Tables**

**Supplementary Table 1.**

Fixed effects parameters of LMMs predicting volume of category-selective activation in lateral VTC ROIs. Related to Fig 1.

| ROI | Contrast | parameter | β | CI | df | t | *p* | *p FDR* |
|---|---|---|---|---|---|---|---|---|
| Left lateral VTC | Numbers | intercept | 426.58 | 104,749 | 126 | 2.62 | 0.01 | |
| | | age | 0.11 | -2.15,2.38 | 126 | 0.10 | 0.92 | 0.92 |
| | Words | intercept | 200.41 | -280, 681 | 126 | 0.82 | 0.41 | |
| | | age | 4.14 | 0.81,7.48 | 126 | 2.46 | 0.015 | 0.044 |
| | Limbs | intercept | 1254.60 | 806,1702 | 126 | 5.54 | $1.7x10^{-7}$ | |
| | | age | -3.79 | -6.84,-0.74 | 126 | -2.46 | 0.015 | 0.044 |
| | Headless bodies | intercept | 348.69 | 151,546 | 126 | 3.50 | $6.53x10^{-4}$ | |
| | | age | -0.589 | -1.98,0.80 | 126 | -0.84 | 0.40 | 0.50 |
| | Adult faces | intercept | 2.5 | -89,94 | 126 | 0.05 | 0.96 | |
| | | age | 0.89 | 0.26,1.52 | 126 | 2.79 | 0.006 | 0.040 |
| | Child faces | intercept | 62.89 | -112,237 | 126 | 0.71 | 0.48 | |
| | | age | 1.95 | 0.74,3.17 | 126 | 3.18 | 0.002 | 0.036 |
| | Cars | intercept | -15.05 | -120,89.9 | 126 | -0.28 | 0.78 | |
| | | age | 0.52 | -0.22,1.25 | 126 | 1.39 | 0.17 | 0.31 |
| | String instruments | intercept | 112.65 | 0.10,225 | 126 | 1.98 | 0.05 | |
| | | age | -0.07 | -0.86,0.72 | 126 | -0.17 | 0.86 | 0.92 |
| | Houses | intercept | -48.32 | -193,96 | 126 | -0.66 | 0.51 | |
| | | age | 1.26 | 0.27,2.25 | 126 | 2.51 | 0.01 | 0.044 |
| | Corridors | intercept | 16.27 | -71,104 | 126 | 0.37 | 0.71 | |
| | | age | 0.38 | -0.23,0.998 | 126 | 1.23 | 0.22 | 0.37 |
| Right lateral VTC | Numbers | intercept | 281.78 | 26,537 | 126 | 2.18 | 0.03 | |
| | | age | 0.11 | -1.69,1.90 | 126 | 0.12 | 0.91 | 0.92 |
| | Words | intercept | 119.95 | -69,309 | 126 | 1.25 | 0.21 | |
| | | age | 0.66 | -0.66,1.99 | 126 | 0.99 | 0.32 | 0.46 |
| | Limbs | intercept | 978.09 | 632,1325 | 126 | 5.59 | $1.37x10^{-7}$ | |
| | | age | -3.42 | -5.79,-1.04 | 126 | -2.85 | 0.005 | 0.04 |
| | Headless bodies | intercept | 588.93 | 299,879 | 126 | 4.02 | $1.0x10^{-4}$ | |
| | | age | -1.625 | -3.63,0.38 | 126 | -1.61 | 0.11 | 0.22 |
| | Adult faces | intercept | 3.73 | -140,147 | 126 | 0.05 | 0.96 | |
| | | age | 1.32 | 0.34,2.29 | 126 | 2.68 | 0.008 | 0.042 |
| | Child faces | intercept | 174.96 | -86,436 | 126 | 1.33 | 0.187 | |
| | | age | 2.16 | 0.35,3.97 | 126 | 2.37 | 0.02 | 0.049 |
| | Cars | intercept | 9.67 | -120,139 | 126 | 0.15 | 0.88 | |
| | | age | 0.40 | -0.51,1.31 | 126 | 0.87 | 0.39 | 0.50 |
| | String instruments | intercept | 51.87 | 0.16,104 | 126 | 1.99 | 0.05 | |
| | | age | -0.13 | -0.49,0.24 | 126 | -0.69 | 0.49 | 0.58 |
| | Houses | intercept | 75.57 | -114,265 | 126 | 0.79 | 0.43 | |
| | | age | 1.16 | -0.123,2.45 | 126 | 1.79 | 0.08 | 0.17 |
| | Corridors | intercept | 42.67 | -88,173 | 126 | 0.65 | 0.52 | |
| | | age | 0.49 | -0.43,1.4 | 126 | 1. | 0.29 | 0.45 |

**Supplementary Table 2.**

Fixed effects parameters of LMMs predicting volume of category-selective activation in medial VTC ROIs. Related to Fig. 1.

| ROI | Contrast | parameter | β | CI | df | t | *p* | *p FDR* |
|---|---|---|---|---|---|---|---|---|
| Left medial VTC | Numbers | intercept | 121.4 | 6.07,237 | 126 | 2.08 | 0.04 | |
| | | age | -0.009 | -0.82,0.80 | 126 | -0.02 | 0.98 | 0.98 |
| | words | intercept | 75.21 | -29,180 | 126 | 1.43 | 0.16 | |
| | | age | 0.45 | -0.28,1.18 | 126 | 1.21 | 0.23 | 0.73 |
| | limbs | intercept | 118.10 | 20.6,216 | 126 | 2.40 | 0.02 | |
| | | age | -0.36 | -1.04,0.32 | 126 | -1.05 | 0.29 | 0.73 |
| | bodies | intercept | 57.91 | 5.46,110 | 126 | 2.19 | 0.03 | |
| | | age | -0.21 | -0.58,0.15 | 126 | -1.15 | 0.25 | 0.73 |
| | Adult faces | intercept | 32.33 | 3.15,61.5 | 126 | 2.19 | 0.03 | |
| | | age | -0.05 | -0.24,0.14 | 126 | -0.50 | 0.62 | 0.88 |
| | Child faces | intercept | 76.70 | 2.20,151 | 126 | 2.04 | 0.04 | |
| | | age | 0.09 | -0.39,0.57 | 126 | 0.38 | 0.71 | 0.88 |
| | Cars | intercept | -27.86 | -98,42.26 | 126 | -0.79 | 0.43 | |
| | | age | 0.41 | -0.08,0.90 | 126 | 1.67 | 0.10 | 0.73 |
| | String instruments | intercept | 11.47 | -1.69,24.6 | 126 | 1.73 | 0.09 | |
| | | age | -0.04 | -0.13,0.06 | 126 | -0.76 | 0.45 | 0.82 |
| | Houses | intercept | 299.84 | 15.2,584 | 126 | 2.08 | 0.04 | |
| | | age | 1.55 | -0.38,3.47 | 126 | 1.59 | 0.11 | 0.73 |
| | Corridors | intercept | 558.91 | 315,803 | 126 | 4.54 | $1.3x10^{-5}$ | |
| | | age | -0.17 | -1.85,1.51 | 126 | -0.20 | 0.84 | 0.93 |
| Right medial VTC | Numbers | intercept | 32.53 | -32.5,97.5 | 126 | 0.99 | 0.32 | |
| | | age | 0.15 | -0.31,0.61 | 126 | 0.66 | 0.51 | 0.85 |
| | words | intercept | 15.92 | -44,76 | 126 | 0.53 | 0.60 | |
| | | age | 0.22 | -0.20,0.63 | 126 | 1.03 | 0.31 | 0.73 |
| | limbs | intercept | 42.23 | -12,97 | 126 | 1.53 | 0.13 | |
| | | age | 0.02 | -0.36,0.41 | 126 | 0.13 | 0.90 | 0.95 |
| | bodies | intercept | 42.03 | -0.10,84.16 | 126 | 1.97 | 0.05 | |
| | | age | -0.14 | -0.43,0.16 | 126 | -0.91 | 0.37 | 0.73 |
| | Adult faces | intercept | 14.55 | -13.36,42.45 | 126 | 1.03 | 0.30 | |
| | | age | -0.03 | -0.22,0.16 | 126 | -0.31 | 0.75 | 0.89 |
| | Child faces | intercept | 33.50 | -11.7,78.72 | 126 | 1.47 | 0.15 | |
| | | age | -0.06 | -0.37,0.25 | 126 | -0.40 | 0.69 | 0.88 |
| | Cars | intercept | -10.02 | -49.47,29.43 | 126 | -0.50 | 0.62 | |
| | | age | 0.19 | -0.09,0.47 | 126 | 1.34 | 0.18 | 0.73 |
| | String instruments | intercept | 13.64 | -0.68,28 | 126 | 1.89 | 0.06 | |
| | | age | -0.05 | -0.15,0.05 | 126 | -0.96 | 0.34 | 0.73 |
| | Houses | intercept | 357.67 | 57.6,657.7 | 126 | 2.36 | 0.02 | |
| | | age | 1.80 | -0.27,3.87 | 126 | 1.72 | 0.09 | 0.73 |
| | Corridors | intercept | 610.52 | 334,887 | 126 | 4.37 | $2.55x10^{-5}$ | |
| | | age | 0.46 | -1.43,2.36 | 126 | 0.48 | 0.63 | 0.88 |

**Supplementary Table 3.**

Likelihood ratio tests determining if the number of selective voxels in VTC is better predicted if motion during scanning is included in the model. Related to Fig. 1.

| ROI | contrast | No. | Model | df | LRStat | *p* | *pFDR* |
|---|---|---|---|---|---|---|---|
| Left lateral VTC | numbers | 1 | Nr of Sel. voxels ~ age + (1\|subj) | 4 | | | |
| | | 2 | Nr of Sel. voxels ~ age + motion + (1\|subj) | 5 | 1.17 | 0.28 | 0.66 |
| | words | 1 | Nr of Sel. voxels ~ age + (1\|subj) | 4 | | | |
| | | 2 | Nr of Sel. voxels ~ age + motion + (1\|subj) | 5 | 0.39 | 0.53 | 0.79 |
| | limbs | 1 | Nr of Sel. voxels ~ age+ (1\|subj) | 4 | | | |
| | | 2 | Nr of Sel. voxels ~ age+ motion+ (1\|subj) | 5 | 0.83 | 0.36 | 0.66 |
| | bodies | 1 | Nr of Sel. voxels ~ age+ (1\|subj) | 4 | | | |
| | | 2 | Nr of Sel. voxels ~ age+ motion+ (1\|subj) | 5 | 0.009 | 0.93 | 0.93 |
| | adult faces | 1 | Nr of Sel. voxels ~ age + (1\|subj) | 4 | | | |
| | | 2 | Nr of Sel. voxels ~ age+ motion + (1\|subj) | 5 | 0.12 | 0.73 | 0.89 |
| | child faces | 1 | Nr of Sel. voxels ~ age + (1\|subj) | 4 | | | |
| | | 2 | Nr of Sel. voxels ~ age + motion + (1\|subj) | 5 | 0.21 | 0.64 | 0.86 |
| | cars | 1 | Nr of Sel. voxels ~ age + (1\|subj) | 4 | | | |
| | | 2 | Nr of Sel. voxels ~ age + motion + (1\|subj) | 5 | 3.02 | 0.08 | 0.66 |
| | string instr. | 1 | Nr of Sel. voxels ~ age+ (1\|subj) | 4 | | | |
| | | 2 | Nr of Sel. voxels ~ age+ motion+ (1\|subj) | 5 | 3.43 | 0.06 | 0.66 |
| | houses | 1 | Nr of Sel. voxels ~ age+ (1\|subj) | 4 | | | |
| | | 2 | Nr of Sel. voxels ~ age+ motion+ (1\|subj) | 5 | 0.007 | 0.93 | 0.93 |
| | corridors | 1 | Nr of Sel. voxels ~ age+ (1\|subj) | 4 | | | |
| | | 2 | Nr of Sel. voxels ~ age+ motion+ (1\|subj) | 5 | 2.13 | 0.14 | 0.66 |
| Right lateral VTC | numbers | 1 | Nr of Sel. voxels ~ age + (1\|subj) | 4 | | | |
| | | 2 | Nr of Sel. voxels ~ age + motion + (1\|subj) | 5 | 0.01 | 0.92 | 0.93 |
| | words | 1 | Nr of Sel. voxels ~ age + (1\|subj) | 4 | | | |
| | | 2 | Nr of Sel. voxels ~ age + motion + (1\|subj) | 5 | 1.23 | 0.27 | 0.66 |
| | limbs | 1 | Nr of Sel. voxels ~ age+ (1\|subj) | 4 | | | |
| | | 2 | Nr of Sel. voxels ~ age+ motion+ (1\|subj) | 5 | 0.10 | 0.75 | 0.89 |
| | bodies | 1 | Nr of Sel. voxels ~ age + (1\|subj) | 4 | | | |
| | | 2 | Nr of Sel. voxels ~ age+ motion + (1\|subj) | 5 | 0.61 | 0.43 | 0.72 |
| | adult faces | 1 | Nr of Sel. voxels ~ age + (1\|subj) | 4 | | | |
| | | 2 | Nr of Sel. voxels ~ age + motion + (1\|subj) | 5 | 0.82 | 0.36 | 0.66 |
| | child faces | 1 | Nr of Sel. voxels ~ age + (1\|subj) | 4 | | | |
| | | 2 | Nr of Sel. voxels ~ age + motion + (1\|subj) | 5 | 0.86 | 0.35 | 0.66 |
| | cars | 1 | Nr of Sel. voxels ~ age+ (1\|subj) | 4 | | | |
| | | 2 | Nr of Sel. voxels ~ age+ motion+ (1\|subj) | 5 | 1.28 | 0.26 | 0.66 |
| | string instr. | 1 | Nr of Sel. voxels ~ age + (1\|subj) | 4 | | | |
| | | 2 | Nr of Sel. voxels ~ age+ motion + (1\|subj) | 5 | 1.60 | 0.21 | 0.66 |
| | houses | 1 | Nr of Sel. voxels ~ age+ (1\|subj) | 4 | | | |
| | | 2 | Nr of Sel. voxels ~ age+ motion+ (1\|subj) | 5 | 0.97 | 0.33 | 0.66 |
| | corridors | 1 | Nr of Sel. voxels ~ age + (1\|subj) | 4 | | | |
| | | 2 | Nr of Sel. voxels ~ age+ motion + (1\|subj) | 5 | 0.35 | 0.55 | 0.79 |

**Supplementary Table 4.**

Fixed effects parameters of LMMs predicting volume of category-selective activation in lateral VTC ROIs with the additional factor tSNR. Related to Fig. 1.

| ROI | contrast | *p*Value of LR-Test $Model_{age}$ vs. $model_{Age\_and\_tSNR}$ | parameter | β | CI | df | t | *p* | *P FDR* |
|---|---|---|---|---|---|---|---|---|---|
| Lateral VTC lh | numbers | 0.10 | Intercept | 30.89 | -531,593 | 125 | 0.11 | 0.91 | |
| | | | age | 0.2 | -2.11,2.51 | 125 | 0.17 | 0.86 | 0.86 |
| | | | tSNR | 5.08 | -0.78,10.94 | 125 | 1.72 | 0.089 | |
| | words | 0.02 | Intercept | -579 | $-1.4x10^{+03}$,219 | 125 | -1.44 | 0.153 | |
| | | | age | 4.48 | 1.20,7.76 | 125 | 2.7 | 0.008 | 0.03 |
| | | | tSNR | 9.76 | 1.69,17.82 | 125 | 2.39 | 0.018 | |
| | limbs | $8.98x10^{-04}$ | Intercept | 278 | -440,995 | 125 | 0.77 | 0.45 | |
| | | | age | -3.33 | -6.3,-0.4 | 125 | -2.24 | 0.027 | 0.068 |
| | | | tSNR | 12.13 | 5.08,19.19 | 125 | 3.4 | $8.95x10^{-04}$ | |
| | bodies | 0.02 | Intercept | 30.98 | -308,370 | 125 | 0.18 | 0.86 | |
| | | | age | -0.64 | -2.03,0.76 | 125 | -0.9 | 0.37 | 0.49 |
| | | | tSNR | 4.28 | 0.72,7.85 | 125 | 2.38 | 0.02 | |
| | Adult faces | 0.15 | Intercept | -87 | -240,65.3 | 125 | -1.13 | 0.26 | |
| | | | age | 0.93 | 0.30,1.56 | 125 | 2.94 | 0.0039 | 0.02 |
| | | | tSNR | 1.12 | -0.41,2.65 | 125 | 1.45 | 0.15 | |
| | Child faces | 0.03 | Intercept | -190.99 | -482,100 | 125 | -1.3 | 0.2 | |
| | | | age | 2.05 | 0.85,3.25 | 125 | 3.39 | $9.48x10^{-04}$ | 0.006 |
| | | | tSNR | 3.2 | 0.2,6.2 | 125 | 2.14 | 0.03 | |
| | Cars | 0.18 | Intercept | -110 | -286,65 | 125 | -1.25 | 0.22 | |
| | | | age | 0.51 | -0.23,1.24 | 125 | 1.37 | 0.17 | 0.35 |
| | | | tSNR | 1.27 | -0.61,3.15 | 125 | 1.34 | 0.18 | |
| | String instr. | 0.03 | Intercept | -51.15 | -237,135 | 125 | -0.54 | 0.59 | |
| | | | age | -0.09 | -0.86,0.69 | 125 | -0.22 | 0.83 | 0.86 |
| | | | tSNR | 2.18 | 0.19,4.17 | 125 | 2.17 | 0.03 | |
| | Houses | 0.37 | Intercept | 40.42 | -201,282 | 125 | 0.33 | 0.74 | |
| | | | age | 1.21 | 0.22,2.2 | 125 | 2.41 | 0.017 | 0.049 |
| | | | tSNR | -1.09 | -3.49,1.3 | 125 | -0.9 | 0.37 | |
| | Corridors | 0.54 | Intercept | 53.78 | -95.7,203.2 | 125 | 0.71 | 0.48 | |
| | | | age | 0.38 | -0.24,0.99 | 125 | 1.22 | 0.23 | 0.38 |
| | | | tSNR | -0.49 | -2.06,1.09 | 125 | -0.61 | 0.54 | |
| Lateral VTC rh | numbers | 0.16 | Intercept | 26.25 | -409.9,462.4 | 125 | 0.12 | 0.91 | |
| | | | age | 0.2 | -1.64,2.04 | 125 | 0.22 | 0.83 | 0.86 |
| | | | tSNR | 3.46 | -1.17,8.09 | 125 | 1.48 | 0.14 | |
| | words | 0.41 | Intercept | 9.35 | -309,327 | 125 | 0.06 | 0.95 | |
| | | | age | 0.72 | -0.62,2.06 | 125 | 1.06 | 0.29 | 0.45 |
| | | | tSNR | 1.47 | -1.85,4.79 | 125 | 0.88 | 0.38 | |
| | limbs | 0.02 | Intercept | 432.91 | -120,986 | 125 | 1.55 | 0.12 | |
| | | | age | -2.99 | -5.33,-0.66 | 125 | -2.54 | 0.012 | 0.04 |
| | | | tSNR | 6.96 | 1.3,12.6 | 125 | 2.43 | 0.016 | |
| | bodies | 0.01 | Intercept | 122 | -345,590 | 125 | 0.52 | 0.61 | |

| | | | | | | | | |
|---|---|---|---|---|---|---|---|---|
| | | age | -1.3 | -3.28,0.67 | 125 | -1.3 | 0.19 | 0.35 |
| | | tSNR | 6.04 | 1.24,10.84 | 125 | 2.49 | 0.01 | |
| Adult faces | $3.84x10^{-05}$ | Intercept | -372 | -589,-154 | 125 | -3.39 | $9.32x10^{-04}$ | |
| | | age | 1.6 | 0.69,2.52 | 125 | 3.46 | $7.39x10^{-04}$ | 0.006 |
| | | tSNR | 4.82 | 2.61,7.04 | 125 | 4.31 | $3.29x10^{-05}$ | |
| Child faces | $4.72x10^{-08}$ | Intercept | -723.36 | $-1.1x10^{+03}$, -346 | 125 | -3.79 | $2.30x10^{-04}$ | |
| | | age | 2.78 | 1.19,4.38 | 125 | 3.46 | $7.44x10^{-04}$ | 0.006 |
| | | tSNR | 11.63 | 7.7,15.6 | 125 | 5.86 | $3.83x10^{-08}$ | |
| Cars | 0.40 | Intercept | 83.84 | -132.2,299.9 | 125 | 0.77 | 0.44 | |
| | | age | 0.36 | -0.55,1.27 | 125 | 0.78 | 0.44 | 0.55 |
| | | tSNR | -0.97 | -3.25,1.30 | 125 | -0.85 | 0.40 | |
| String instr. | 0.23 | Intercept | 9.87 | -75.9,95.6 | 125 | 0.23 | 0.82 | |
| | | age | -0.11 | -0.48,0.25 | 125 | -0.62 | 0.54 | 0.63 |
| | | tSNR | 0.57 | -0.4,1.5 | 125 | 1.21 | 0.23 | |
| Houses | 0.51 | Intercept | -4.43 | -312,304 | 125 | -0.03 | 0.98 | |
| | | age | 1.22 | -0.08,2.52 | 125 | 1.86 | 0.06 | 0.14 |
| | | tSNR | 1.02 | -2.09,4.14 | 125 | 0.65 | 0.52 | |
| Corridors | 0.73 | Intercept | 74.14 | -143,291 | 125 | 0.68 | 0.50 | |
| | | age | 0.47 | -0.45,1.38 | 125 | 1.01 | 0.32 | 0.45 |
| | | tSNR | -0.41 | -2.69,1.88 | 125 | -0.35 | 0.73 | |

Note, LR-Test: Likelihood Ratio test.

**Supplementary Table 5.**

Fixed effects parameters of LMMs predicting volume of category-selective activation in medial VTC ROIs with the additional factor tSNR. Related to Fig. 1.

| ROI | contrast | *p*Val of LR-Test $Model_{age}$ vs $model_{Age\_and\_tSNR}$ | parameter | β | CI | df | t | *p* | *P FDR* |
|---|---|---|---|---|---|---|---|---|---|
| medial VTC lh | numbers | $1.81x10^{-04}$ | Intercept | -232 | -443,-21 | 125 | -2.18 | 0.03 | |
| | | | age | 0.29 | -0.49,1.06 | 125 | 0.73 | 0.47 | 0.67 |
| | | | tSNR | 4.84 | 2.35, 7.33 | 125 | 3.85 | $1.87x10^{-04}$ | |
| | words | 0.24 | Intercept | -29.63 | -232,173 | 125 | -0.29 | 0.77 | |
| | | | age | 0.54 | -0.21,1.29 | 125 | 1.43 | 0.15 | 0.56 |
| | | | tSNR | 1.43 | -0.95,3.81 | 125 | 1.19 | 0.24 | |
| | limbs | 0.15 | Intercept | 2.74 | -180,186 | 125 | 0.03 | 0.98 | |
| | | | age | -0.26 | -0.94,0.43 | 125 | -0.74 | 0.46 | 0.67 |
| | | | tSNR | 1.57 | -0.55,3.68 | 125 | 1.46 | 0.15 | |
| | bodies | 0.46 | Intercept | 89.26 | -9.56,188.08 | 125 | 1.79 | 0.08 | |
| | | | age | -0.24 | -0.61,0.13 | 125 | -1.28 | 0.20 | 0.56 |
| | | | tSNR | -0.42 | -1.56,0.71 | 125 | -0.74 | 0.46 | |
| | Adult faces | 0.06 | Intercept | -6.98 | -57.8,43.8 | 125 | -0.27 | 0.79 | |
| | | | age | -0.01 | -0.21,0.18 | 125 | -0.12 | 0.91 | 0.91 |
| | | | tSNR | 0.53 | -0.03,1.09 | 125 | 1.88 | 0.06 | |
| | Child faces | 0.047 | Intercept | -28.87 | -156,98.7 | 125 | -0.45 | 0.65 | |
| | | | age | 0.2 | -0.29,0.69 | 125 | 0.81 | 0.42 | 0.67 |
| | | | tSNR | 1.4 | 0.01,2.8 | 125 | 2.0 | 0.048 | |
| | Cars | 0.56 | Intercept | -61.36 | -195,73 | 125 | -0.91 | 0.37 | |
| | | | age | 0.44 | -0.06,0.94 | 125 | 1.76 | 0.08 | 0.54 |
| | | | tSNR | 0.46 | -1.1,2.02 | 125 | 0.58 | 0.56 | |
| | String instr. | 0.18 | Intercept | 26.45 | 2.15,50.74 | 125 | 2.15 | 0.03 | |
| | | | age | -0.04 | -0.13,0.05 | 125 | -0.85 | 0.40 | 0.67 |
| | | | tSNR | -0.23 | -0.52,0.07 | 125 | -1.52 | 0.13 | |
| | Houses | $2.62x10^{-04}$ | Intercept | -478 | -968,11.7 | 125 | -1.93 | 0.06 | |
| | | | age | 2.31 | 0.44,4.18 | 125 | 2.45 | 0.02 | 0.18 |
| | | | tSNR | 10.44 | 4.9,15.9 | 125 | 3.75 | $2.68x10^{-04}$ | |
| | Corridors | $1.13x10^{-04}$ | Intercept | -167 | -595,261 | 125 | -0.77 | 0.44 | |
| | | | age | 0.49 | -1.12,2.11 | 125 | 0.60 | 0.55 | 0.73 |
| | | | tSNR | 9.83 | 4.9,14.7 | 125 | 3.98 | $1.18x10^{-04}$ | |
| medial VTC rh | numbers | 0.26 | Intercept | -30.15 | -169,109 | 125 | -0.43 | 0.67 | |
| | | | age | 0.11 | -0.37,0.6 | 125 | 0.46 | 0.65 | 0.76 |
| | | | tSNR | 1.17 | -0.68,3.03 | 125 | 1.25 | 0.21 | |
| | words | 0.57 | Intercept | -13.23 | -133, 106 | 125 | -0.22 | 0.83 | |
| | | | age | 0.24 | -0.19,0.67 | 125 | 1.11 | 0.27 | 0.56 |
| | | | tSNR | 0.45 | -1.12,2.02 | 125 | 0.57 | 0.57 | |
| | limbs | 0.24 | Intercept | -15.51 | -126,95 | 125 | -0.28 | 0.78 | |
| | | | age | 0.07 | -0.32,0.46 | 125 | 0.35 | 0.73 | 0.81 |
| | | | tSNR | 0.89 | -0.58,2.36 | 125 | 1.19 | 0.24 | |
| | bodies | 0.39 | Intercept | 74.48 | -9.6,158.57 | 125 | 1.75 | 0.08 | |

| | | | | | | | | |
|---|---|---|---|---|---|---|---|---|
| | | age | -0.16 | -0.46,0.14 | 125 | -1.08 | 0.28 | 0.56 |
| | | tSNR | -0.49 | -1.60,0.62 | 125 | -0.88 | 0.38 | |
| Adult faces | 0.68 | Intercept | 4.66 | -50.09,59.40 | 125 | 0.17 | 0.87 | |
| | | age | -0.022 | -0.22,0.18 | 125 | -0.22 | 0.83 | 0.87 |
| | | tSNR | 0.15 | -0.56,0.86 | 125 | 0.41 | 0.68 | |
| Child faces | 0.68 | Intercept | 49.05 | -37.7,135.8 | 125 | 1.12 | 0.27 | |
| | | age | -0.08 | -0.39,0.24 | 125 | -0.48 | 0.64 | 0.76 |
| | | tSNR | -0.23 | -1.35,0.88 | 125 | -0.42 | 0.68 | |
| Cars | 0.87 | Intercept | -3.5 | -85.76,78.75 | 125 | -0.08 | 0.93 | |
| | | age | 0.18 | -0.10,0.47 | 125 | 1.27 | 0.21 | 0.56 |
| | | tSNR | -0.1 | -1.21,1.01 | 125 | -0.18 | 0.86 | |
| String instr. | 0.13 | Intercept | 34.28 | 4.74,63.83 | 125 | 2.30 | 0.02 | |
| | | age | -0.07 | -0.17,0.04 | 125 | -1.29 | 0.2 | 0.56 |
| | | tSNR | -0.31 | -0.70,0.08 | 125 | -1.56 | 0.12 | |
| Houses | $4.65x10^{-04}$ | Intercept | -504 | $-1.1x10^{+03}$,57 | 125 | -1.78 | 0.08 | |
| | | age | 2.45 | 0.42,4.48 | 125 | 2.39 | 0.02 | 0.18 |
| | | tSNR | 13.26 | 6.0,20.5 | 125 | 3.6 | $4.37x10^{-04}$ | |
| Corridors | $2.11x10^{-04}$ | Intercept | -219 | -727,288 | 125 | -0.86 | 0.39 | |
| | | age | 1.14 | -0.70,2.98 | 125 | 1.22 | 0.22 | 0.56 |
| | | tSNR | 12.66 | 6.12,19.2 | 125 | 3.83 | $1.99x10^{-04}$ | |

**Supplementary Table 6.**

Fixed effects parameters of LMMs predicting ROI volume. Related to Fig. 2.

| ROI | parameter | β | CI | df | t | *p* | *p FDR* |
|---|---|---|---|---|---|---|---|
| Left mOTS-words | intercept | 130.61 | -39.93,301 | 101 | 1.52 | 0.13 | |
| | age | -0.07 | -1.22,1.08 | 101 | -0.12 | 0.90 | 0.90 |
| Right mOTS-words | intercept | 8.93 | -95.1,113 | 33 | 0.17 | 0.86 | |
| | age | 0.25 | -0.43,0.92 | 33 | 0.74 | 0.46 | 0.66 |
| Left pOTS-words | intercept | -19.96 | -205,165 | 117 | -0.21 | 0.83 | |
| | age | 1.71 | 0.45,2.98 | 117 | 2.68 | 0.009 | 0.02 |
| Right pOTS-words | intercept | 21.33 | -73.9,116.6 | 71 | 0.45 | 0.66 | |
| | age | 0.37 | -0.32,1.06 | 71 | 1.07 | 0.29 | 0.48 |
| Left OTS-limbs | intercept | 369.43 | 217,522 | 124 | 4.80 | $4.40x10^{-6}$ | |
| | age | -1.70 | -2.59,-0.81 | 124 | -3.78 | $2.40x10^{-4}$ | 0.002 |
| Right OTS-limbs | intercept | 204.46 | 105,304 | 124 | 4.07 | $8.41x10^{-5}$ | |
| | age | -0.89 | -1.53,-0.24 | 124 | -2.73 | 0.007 | 0.02 |
| Left mFus-faces | intercept | 99.77 | 7.39,192 | 105 | 2.14 | 0.03 | |
| | age | 0.11 | -0.47,0.69 | 105 | 0.38 | 0.71 | 0.79 |
| Right mFus-faces | intercept | 90.54 | -6.41,187.48 | 100 | 1.85 | 0.07 | |
| | age | 0.20 | -0.46,0.85 | 100 | 0.60 | 0.55 | 0.69 |
| Left pFus-faces | intercept | 23.36 | -64.37,111.1 | 118 | 0.53 | 0.60 | |
| | age | 0.75 | 0.17,1.33 | 118 | 2.55 | 0.01 | 0.02 |
| Right pFus-Faces | intercept | -23.39 | -163,116 | 96 | -0.33 | 0.74 | |
| | age | 1.41 | 0.46,2.35 | 96 | 2.96 | 0.004 | 0.019 |

**Supplementary Table 7.**

Fixed effects parameters of LMMs predicting changes in selectivity by age in developing parts of ROIs. Listed by figure order. Related to Fig. 3 and Fig. S9.

| ROI | Contrast | parameter | β | CI | df | t | *p* | *p FDR* |
|---|---|---|---|---|---|---|---|---|
| Emerging lh pOTS-word | numbers | intercept | 1.19 | -0.04,2.41 | 110 | 1.92 | 0.06 | |
| | | *age* | 0.010 | 0.001,0.018 | 110 | 2.29 | 0.02 | 0.07 |
| | words | intercept | -0.09 | -1.27,1.094 | 110 | -0.14 | 0.89 | |
| | | *age* | 0.025 | 0.017,0.033 | 110 | 6.17 | $1.15x10^{-8}$ | $1.92x10^{-7}$ |
| | limbs | intercept | 3.84 | 2.59,5.09 | 110 | 6.09 | $1.67x10^{-8}$ | |
| | | *age* | -0.020 | -0.028,-0.012 | 110 | -4.74 | $6.42x10^{-6}$ | $8.03x10^{-5}$ |
| | bodies | intercept | 1.44 | 0.28,2.60 | 110 | 2.46 | 0.02 | |
| | | *age* | -0.015 | -0.022,-0.007 | 110 | -3.68 | $3.66x10^{-4}$ | 0.002 |
| | Adult faces | intercept | -1.25 | -2.51, 0.02 | 110 | -1.96 | 0.05 | |
| | | *age* | -0.007 | -0.016,0.001 | 110 | -1.68 | 0.10 | 0.23 |
| | Child faces | intercept | -1.13 | -2.69,0.43 | 110 | -1.44 | 0.15 | |
| | | *age* | -0.002 | -0.012,0.009 | 110 | -0.30 | 0.76 | 0.83 |
| | Cars | intercept | -0.73 | -1.90,0.44 | 110 | -1.23 | 0.22 | |
| | | *age* | -0.002 | -0.010,0.006 | 110 | -0.50 | 0.62 | 0.77 |
| | String instr. | intercept | -0.27 | -1.30,0.77 | 110 | -0.51 | 0.61 | |
| | | *age* | 0.001 | -0.006,0.008 | 110 | 0.36 | 0.72 | 0.81 |
| | Houses | intercept | -0.48 | -1.72,0.75 | 110 | -0.78 | 0.44 | |
| | | *age* | -0.002 | -0.010,0.007 | 110 | -0.46 | 0.65 | 0.77 |
| | Corridors | intercept | -0.91 | -2.28,0.45 | 110 | -1.33 | 0.19 | |
| | | *age* | -0.008 | -0.017,0.002 | 110 | -1.66 | 0.10 | 0.23 |
| Waning lh OTS-limbs | numbers | intercept | 0.58 | -0.47,1.63 | 120 | 1.09 | 0.28 | |
| | | *age* | -0.002 | -0.009,0.006 | 120 | -0.47 | 0.64 | 0.77 |
| | words | intercept | 0.30 | -0.97,1.57 | 120 | 0.47 | 0.64 | |
| | | *age* | 0.005 | -0.004,0.014 | 120 | 1.14 | 0.26 | 0.44 |
| | limbs | intercept | 7.39 | 6.09,8.68 | 120 | 11.30 | $1.31x10^{-20}$ | |
| | | *age* | -0.030 | -0.039,-0.021 | 120 | -6.83 | $3.77x10^{-10}$ | $1.88x10^{-8}$ |
| | bodies | intercept | 3.21 | 1.97,4.46 | 120 | 5.12 | $1.18x10^{-6}$ | |
| | | *age* | -0.010 | $-0.018,-9.2x10^{-04}$ | 120 | -2.19 | 0.03 | 0.09 |
| | Adult faces | intercept | -2.26 | -3.35,-1.16 | 120 | -4.09 | $7.85x10^{-5}$ | |
| | | *age* | 0.010 | 0.002,0.017 | 120 | 2.59 | 0.01 | 0.04 |
| | Child faces | intercept | -1.08 | -2.31,0.15 | 120 | -1.74 | 0.08 | |
| | | *age* | 0.007 | -0.002,0.015 | 120 | 1.62 | 0.11 | 0.24 |
| | Cars | intercept | -1.19 | -2.16,-0.23 | 120 | -2.44 | 0.02 | |
| | | *age* | $-2.71x10^{-4}$ | -0.007,0.007 | 120 | -0.08 | 0.94 | 0.97 |
| | String instr. | intercept | -1.05 | -1.91, -0.19 | 120 | -2.41 | 0.02 | |
| | | *age* | 0.004 | -0.002,0.010 | 120 | 1.45 | 0.15 | 0.31 |
| | Houses | intercept | -1.76 | -2.72,-0.79 | 120 | -3.61 | $4.47x10^{-4}$ | |
| | | *age* | 0.002 | -0.005,0.008 | 120 | 0.48 | 0.63 | 0.77 |
| | Corridors | intercept | -2.90 | -3.98, -1.82 | 120 | -5.32 | $4.80x10^{-7}$ | |
| | | *age* | 0.002 | -0.005,0.01 | 120 | 0.57 | 0.57 | 0.77 |
| Emerging lh pFus-faces | numbers | intercept | 0.98 | -0.23,2.19 | 103 | 1.61 | 0.11 | |
| | | *age* | -0.004 | -0.013,0.004 | 103 | -1.00 | 0.32 | 0.51 |

| | | | | | | | |
|---|---|---|---|---|---|---|---|
| | words | intercept | 0.43 | -0.75, 1.61 | 103 | 0.72 | 0.47 | |
| | | *age* | 0.003 | -0.005,0.011 | 103 | 0.79 | 0.43 | 0.66 |
| | limbs | intercept | 3.30 | 2.28,4.32 | 103 | 6.41 | $4.55x10^{-9}$ | |
| | | *age* | -0.022 | -0.028,-0.015 | 103 | -6.24 | $9.79x10^{-9}$ | $1.92x10^{-7}$ |
| | bodies | intercept | 1.71 | 0.60,2.82 | 103 | 3.04 | 0.003 | |
| | | *age* | -0.014 | -0.021,-0.006 | 103 | -3.57 | $5.48x10^{-4}$ | 0.002 |
| | Adult faces | intercept | -0.47 | -1.55,0.62 | 103 | -0.85 | 0.39 | |
| | | *age* | 0.016 | 0.008,0.023 | 103 | 4.09 | $8.51x10^{-5}$ | $4.06x10^{-4}$ |
| | Child faces | intercept | 0.68 | -0.73,2.09 | 103 | 0.96 | 0.34 | |
| | | *age* | 0.020 | 0.010,0.030 | 103 | 4.11 | $8.06x10^{-5}$ | $4.06x10^{-4}$ |
| | Cars | intercept | -0.86 | -2.19,0.47 | 103 | -1.29 | 0.20 | |
| | | *age* | $-2.70x10^{-4}$ | -0.010,0.009 | 103 | -0.06 | 0.95 | 0.97 |
| | String instr. | intercept | -0.82 | -1.87,0.23 | 103 | -1.55 | 0.13 | |
| | | *age* | -0.004 | -0.011,0.003 | 103 | -1.18 | 0.24 | 0.43 |
| | Houses | Intercept | -1.48 | -2.50,-0.45 | 103 | -2.86 | 0.005 | |
| | | *age* | $2.59x10^{-4}$ | -0.007,0.007 | 103 | 0.07 | 0.94 | 0.97 |
| | Corridors | intercept | -3.05 | -4.45,-1.66 | 103 | -4.34 | $3.38x10^{-5}$ | |
| | | *age* | -0.002 | -0.011,0.008 | 103 | -0.35 | 0.73 | 0.81 |
| Waning rh OTS-limbs | numbers | intercept | 0.69 | -0.58,1.96 | 98 | 1.08 | 0.28 | |
| | | *age* | -0.005 | -0.014,0.003 | 98 | -1.24 | 0.22 | 0.40 |
| | words | intercept | -0.74 | -2.04,0.56 | 98 | -1.13 | 0.26 | |
| | | *age* | 0.002 | -0.007,0.011 | 98 | 0.43 | 0.67 | 0.78 |
| | limbs | intercept | 5.80 | 4.41,7.18 | 98 | 8.31 | $5.51x10^{-13}$ | |
| | | *age* | -0.020 | -0.029,-0.010 | 98 | -4.10 | $8.57x10^{-5}$ | $4.06x10^{-4}$ |
| | bodies | intercept | 3.22 | 1.81,4.63 | 98 | 4.53 | $1.65x10^{-5}$ | |
| | | *age* | -0.006 | -0.016,0.004 | 98 | -1.24 | 0.22 | 0.40 |
| | Adult faces | intercept | -2.88 | -4.45,-1.31 | 98 | -3.64 | $4.43x10^{-4}$ | |
| | | *age* | 0.022 | 0.011,0.032 | 98 | 4.09 | $8.93x10^{-5}$ | $4.06x10^{-4}$ |
| | Child faces | intercept | -1.28 | -2.98,0.41 | 98 | -1.50 | 0.14 | |
| | | *age* | 0.018 | 0.007,0.029 | 98 | 3.19 | 0.002 | 0.007 |
| | Cars | intercept | -0.58 | -1.67,0.52 | 98 | -1.04 | 0.30 | |
| | | *age* | -0.005 | -0.013,0.003 | 98 | -1.31 | 0.19 | 0.39 |
| | String instr. | intercept | -0.79 | -1.71,0.12 | 98 | -1.72 | 0.09 | |
| | | *age* | -0.002 | -0.008,0.005 | 98 | -0.55 | 0.59 | 0.77 |
| | Houses | Intercept | -1.14 | -1.89,-0.39 | 98 | -3.02 | 0.003 | |
| | | *age* | -0.002 | -0.008,0.003 | 98 | -0.91 | 0.366 | 0.57 |
| | Corridors | intercept | -2.69 | -3.94,-1.43 | 98 | -4.26 | $4.77x10^{-5}$ | |
| | | *age* | $-1.45x10^{-4}$ | -0.009,0.009 | 98 | -0.03 | 0.97 | 0.97 |
| Emerging rh pFus-faces | numbers | intercept | 0.66 | -0.53,1.85 | 94 | 1.10 | 0.28 | |
| | | *age* | -0.007 | -0.016,0.001 | 94 | -1.73 | 0.09 | 0.23 |
| | words | intercept | -0.20 | -1.60,1.20 | 94 | -0.30 | 0.77 | |
| | | *age* | -0.003 | -0.013,0.007 | 94 | -0.63 | 0.53 | 0.73 |
| | limbs | intercept | 2.75 | 1.74,3.76 | 94 | 5.40 | $4.94x10^{-7}$ | |
| | | *age* | -0.016 | -0.023,-0.008 | 94 | -4.31 | $4.02x10^{-5}$ | $2.87x10^{-4}$ |
| | bodies | intercept | 1.65 | 0.30,2.99 | 94 | 2.43 | 0.02 | |
| | | *age* | -0.008 | -0.017,0.002 | 94 | -1.67 | 0.098 | 0.23 |

| | | | | | | | |
|---|---|---|---|---|---|---|---|
| Adult faces | intercept | -0.81 | -2.08,0.45 | 94 | -1.29 | 0.20 | |
| | *age* | 0.020 | 0.011,0.029 | 94 | 4.43 | $2.55x10^{-5}$ | $2.12x10^{-4}$ |
| Child faces | intercept | 0.71 | -0.63,2.05 | 94 | 1.06 | 0.30 | |
| | *age* | 0.021 | 0.012,0.031 | 94 | 4.44 | $2.47x10^{-5}$ | $2.12x10^{-4}$ |
| Cars | intercept | -0.40 | -1.54,0.74 | 94 | -0.69 | 0.49 | |
| | *age* | -0.003 | -0.011,0.005 | 94 | -0.72 | 0.47 | 0.68 |
| String instr. | intercept | -1.68 | -2.59,-0.76 | 94 | -3.63 | $4.69x10^{-4}$ | |
| | *age* | 0.002 | -0.004,0.009 | 94 | 0.72 | 0.47 | 0.68 |
| Houses | Intercept | -0.52 | -1.60,0.57 | 94 | -0.94 | 0.35 | |
| | *age* | -0.004 | -0.012,0.003 | 94 | -1.12 | 0.27 | 0.44 |
| Corridors | intercept | -1.51 | -2.87,-0.16 | 94 | -2.22 | 0.03 | |
| | *age* | -0.010 | -0.020,$-3.53x10^{-4}$ | 94 | -2.06 | 0.04 | 0.12 |

**Supplementary Table 8.**

Fixed effects parameters of LMMs predicting changes in selectivity by age in developing parts of ROIs excluding the ROI-defining category from the contrasting categories. Listed by figure order. Related to Fig. 3 and Fig. S9.

| ROI | Contrast | parameter | β | CI | df | t | *p* | *p FDR* |
|---|---|---|---|---|---|---|---|---|
| Emerging lh pOTS-word | numbers | intercept | 1.19 | -0.04,2.41 | 110 | 1.92 | 0.057 | |
| | | *age* | 0.01 | 0.001,0.018 | 110 | 2.29 | 0.024 | 0.128 |
| | limbs | intercept | 3.65 | 2.41,4.89 | 110 | 5.85 | $5.16 \times 10^{-08}$ | |
| | | *age* | -0.016 | -0.024,-0.007 | 110 | -3.78 | $2.59 \times 10^{-04}$ | 0.0056 |
| | bodies | intercept | 1.405 | 0.29,2.52 | 110 | 2.50 | 0.01 | |
| | | *age* | -0.01 | -0.018,-0.003 | 110 | -2.78 | 0.007 | 0.064 |
| | Adult faces | intercept | -1.5 | -2.70,-0.29 | 110 | -2.47 | 0.015 | |
| | | *age* | -0.002 | -0.01,0.006 | 110 | -0.47 | 0.639 | 0.83 |
| | Child faces | intercept | -1.32 | -2.83,0.18 | 110 | -1.74 | 0.085 | |
| | | *age* | 0.003 | -0.008,0.013 | 110 | 0.49 | 0.622 | 0.83 |
| | Cars | intercept | -0.71 | -1.84,0.42 | 110 | -1.24 | 0.22 | |
| | | *age* | $9.9 \times 10^{-4}$ | -0.007, 0.009 | 110 | 0.25 | 0.8 | 0.88 |
| | String instr. | intercept | -0.46 | -1.5, 0.58 | 110 | -0.88 | 0.38 | |
| | | *age* | 0.006 | $-8.0 \times 10^{-4}$, 0.013 | 110 | 1.76 | 0.08 | 0.25 |
| | Houses | intercept | -0.56 | -1.83,0.71 | 110 | -0.87 | 0.38 | |
| | | *age* | 0.001 | -0.007,0.010 | 110 | 0.27 | 0.78 | 0.88 |
| | Corridors | intercept | -1.09 | -2.43,0.25 | 110 | -1.61 | 0.11 | |
| | | *age* | -0.003 | -0.012,0.006 | 110 | -0.75 | 0.45 | 0.76 |
| Waning lh OTS-limbs | numbers | intercept | 1.54 | 0.48,2.59 | 120 | 2.89 | 0.005 | |
| | | *age* | -0.006 | -0.013,0.002 | 120 | -1.51 | 0.13 | 0.34 |
| | words | intercept | 1.26 | -0.02,2.53 | 120 | 1.95 | 0.054 | |
| | | *age* | 0.001 | -0.008,0.010 | 120 | 0.24 | 0.81 | 0.88 |
| | bodies | intercept | 3.21 | 1.97,4.46 | 120 | 5.12 | $1.18 \times 10^{-6}$ | |
| | | *age* | -0.01 | $-0.018,-9.2 \times 10^{-4}$ | 120 | -2.19 | 0.03 | 0.14 |
| | Adult faces | intercept | -1.27 | -2.35,-0.18 | 120 | -2.31 | 0.02 | |
| | | *age* | 0.006 | -0.001,0.014 | 120 | 1.64 | 0.10 | 0.30 |
| | Child faces | intercept | -0.25 | -1.46,0.96 | 120 | -0.41 | 0.68 | |
| | | *age* | 0.004 | -0.005,0.012 | 120 | 0.86 | 0.39 | 0.76 |
| | Cars | intercept | -0.49 | -1.40,0.41 | 120 | -1.08 | 0.28 | |
| | | *age* | -0.002 | -0.009,0.004 | 120 | -0.74 | 0.46 | 0.76 |
| | String instr. | intercept | -0.21 | -1.09,0.68 | 120 | -0.46 | 0.64 | |
| | | *age* | 0.002 | -0.004,0.008 | 120 | 0.59 | 0.56 | 0.83 |
| | Houses | intercept | -1.04 | -1.89,-0.19 | 120 | -2.43 | 0.017 | |
| | | *age* | $-9.1 \times 10^{-4}$ | -0.007,0.005 | 120 | -0.30 | 0.76 | 0.88 |
| | Corridors | intercept | -2.14 | -3.17,-1.11 | 120 | -4.12 | $6.95 \times 10^{-5}$ | |
| | | *age* | $-5.1 \times 10^{-4}$ | -0.008,0.007 | 120 | -0.14 | 0.89 | 0.93 |
| Emerging lh pFus-faces | numbers | intercept | 1.06 | -0.10,2.23 | 103 | 1.81 | 0.07 | |
| | | *age* | $3.8 \times 10^{-4}$ | -0.008,0.008 | 103 | 0.09 | 0.92 | 0.95 |
| | words | intercept | 0.52 | -0.63,1.67 | 103 | 0.89 | 0.38 | |
| | | *age* | 0.007 | $-5.3 \times 10^{-4}$, 0.015 | 103 | 1.85 | 0.07 | 0.22 |
| | limbs | intercept | 3.31 | 2.26,4.37 | 103 | 6.23 | $1.03 \times 10^{-8}$ | |

| | | | | | | | | |
|---|---|---|---|---|---|---|---|---|
| | | *age* | -0.017 | -0.024,-0.01 | 103 | -4.76 | 6.26x10$^{-6}$ | 2.69x10$^{-4}$ |
| | bodies | intercept | 1.78 | 0.63,2.92 | 103 | 3.07 | 0.003 | |
| | | *age* | -0.008 | -0.016,-3.8x10$^{-4}$ | 103 | -2.08 | 0.04 | 0.17 |
| | Cars | intercept | -0.73 | -2.08,0.62 | 103 | -1.08 | 0.28 | |
| | | *age* | 0.004 | -0.006,0.013 | 103 | 0.76 | 0.45 | 0.76 |
| | String instr. | intercept | -0.97 | -2.03,0.09 | 103 | -1.82 | 0.07 | |
| | | *age* | 0.002 | -0.005,0.01 | 103 | 0.65 | 0.51 | 0.82 |
| | Houses | Intercept | -1.74 | -2.79,-0.68 | 103 | -3.27 | 0.001 | |
| | | *age* | 0.006 | -0.002,0.013 | 103 | 1.56 | 0.12 | 0.32 |
| | Corridors | intercept | -3.03 | -4.34,-1.71 | 103 | -4.56 | 1.4x10$^{-5}$ | |
| | | *age* | 0.003 | -0.007,0.012 | 103 | 0.56 | 0.58 | 0.83 |
| Waning rh OTS-limbs | numbers | intercept | 1.58 | 0.27,2.90 | 98 | 2.39 | 0.02 | |
| | | *age* | -0.009 | -0.018,-2.7x10$^{-4}$ | 98 | -2.05 | 0.04 | 0.17 |
| | words | intercept | 0.10 | -1.21,1.40 | 98 | 0.15 | 0.88 | |
| | | *age* | -0.001 | -0.01,0.007 | 98 | -0.32 | 0.75 | 0.88 |
| | bodies | intercept | 3.22 | 1.81,4.63 | 98 | 4.53 | 1.65x10$^{-5}$ | |
| | | *age* | -0.006 | -0.016,0.004 | 98 | -1.24 | 0.22 | 0.47 |
| | Adult faces | intercept | -1.92 | -3.48,-0.37 | 98 | -2.46 | 0.016 | |
| | | *age* | 0.018 | 0.008,0.029 | 98 | 3.50 | 6.95x10$^{-4}$ | 0.01 |
| | Child faces | intercept | -0.48 | -2.16,1.19 | 98 | -0.57 | 0.57 | |
| | | *age* | 0.015 | 0.004,0.026 | 98 | 2.65 | 0.009 | 0.067 |
| | Cars | intercept | 0.10 | -0.96,1.15 | 98 | 0.18 | 0.86 | |
| | | *age* | -0.007 | -0.015,1.25x10$^{-4}$ | 98 | -1.95 | 0.054 | 0.19 |
| | String instr. | intercept | 0.047 | -0.89,0.98 | 98 | 0.10 | 0.92 | |
| | | *age* | -0.005 | -0.011,0.002 | 98 | -1.42 | 0.16 | 0.36 |
| | Houses | Intercept | -0.66 | -1.39,0.07 | 98 | -1.80 | 0.07 | |
| | | *age* | -0.004 | -0.009,0.001 | 98 | -1.42 | 0.16 | 0.36 |
| | Corridors | intercept | -1.90 | -3.10,-0.70 | 98 | -3.14 | 0.002 | |
| | | *age* | -0.003 | -0.012,0.005 | 98 | -0.78 | 0.44 | 0.76 |
| Emerging rh pFus-faces | numbers | intercept | 0.62 | -0.47,1.71 | 94 | 1.13 | 0.26 | |
| | | *age* | -0.002 | -0.010,0.006 | 94 | -0.49 | 0.63 | 0.83 |
| | words | intercept | -0.26 | -1.55,1.03 | 94 | -0.40 | 0.69 | |
| | | *age* | 0.002 | -0.007,0.012 | 94 | 0.52 | 0.61 | 0.83 |
| | limbs | intercept | 2.73 | 1.64,3.82 | 94 | 4.98 | 2.83x10$^{-6}$ | |
| | | *age* | -0.01 | -0.018,-0.002 | 94 | -2.60 | 0.01 | 0.067 |
| | bodies | intercept | 1.46 | -0.002,2.93 | 94 | 1.98 | 0.0503 | |
| | | *age* | 1.56x10$^{-4}$ | -0.010,0.01 | 94 | 0.031 | 0.98 | 0.98 |
| | Cars | intercept | -0.30 | -1.47,0.87 | 94 | -0.51 | 0.61 | |
| | | *age* | 0.001 | -0.007,0.010 | 94 | 0.33 | 0.74 | 0.88 |
| | String instr. | intercept | -1.72 | -2.61,-0.83 | 94 | -3.85 | 2.16x10$^{-4}$ | |
| | | *age* | 0.009 | 0.0024,0.015 | 94 | 2.74 | 0.007 | 0.06 |
| | Houses | Intercept | -0.64 | -1.66,0.38 | 94 | -1.25 | 0.21 | |
| | | *age* | 8.2x10$^{-4}$ | -0.006,0.008 | 94 | 0.22 | 0.82 | 0.88 |
| | Corridors | intercept | -1.64 | -2.90,-0.38 | 94 | -2.58 | 0.01 | |
| | | *age* | -0.004 | -0.013,0.005 | 94 | -0.90 | 0.37 | 0.76 |

**Supplementary Table 9.**

Fixed effects parameters of LMMs predicting changes in response values by age in developing parts of ROIs. Listed by figure order. Related to Fig. 3 and Fig. S9.

| ROI | Contrast | parameter | β | CI | df | t | *p* | *p FDR* |
|---|---|---|---|---|---|---|---|---|
| Emerging lh pOTS-word | numbers | intercept | 0.58 | 0.04,1.12 | 110 | 2.13 | 0.04 | |
| | | *age* | 0.003 | -2.9x10$^{-4}$,0.007 | 110 | 1.82 | 0.07 | 0.17 |
| | words | intercept | 0.17 | -0.42,0.76 | 110 | 0.57 | 0.57 | |
| | | *age* | 0.008 | 0.004,0.012 | 110 | 3.82 | 2.24x10$^{-4}$ | 0.002 |
| | limbs | intercept | 0.78 | 0.30,1.27 | 110 | 3.19 | 0.002 | |
| | | *age* | 3.54x10$^{-4}$ | -0.003,0.004 | 110 | 0.22 | 0.82 | 0.92 |
| | bodies | intercept | 0.40 | 0.06,0.74 | 110 | 2.35 | 0.02 | |
| | | *age* | 7.62x10$^{-4}$ | -0.001,0.003 | 110 | 0.683 | 0.50 | 0.64 |
| | Adult faces | intercept | 0.08 | -0.21,0.37 | 110 | 0.55 | 0.58 | |
| | | *age* | 0.002 | -3.4x10$^{-4}$, 0.004 | 110 | 1.62 | 0.11 | 0.21 |
| | Child faces | intercept | 0.03 | -0.40,0.45 | 110 | 0.13 | 0.90 | |
| | | *age* | 0.003 | -1.6x10$^{-4}$, 0.005 | 110 | 1.87 | 0.06 | 0.16 |
| | Cars | intercept | 0.08 | -0.34,0.51 | 110 | 0.39 | 0.70 | |
| | | *age* | 0.003 | -2.4x10$^{-4}$, 0.005 | 110 | 1.81 | 0.07 | 0.17 |
| | String instr. | intercept | 0.11 | -0.25,0.48 | 110 | 0.61 | 0.54 | |
| | | *age* | 0.004 | 0.001,0.006 | 110 | 2.98 | 0.004 | 0.025 |
| | Houses | intercept | 0.26 | -0.26,0.79 | 110 | 1.00 | 0.32 | |
| | | *age* | 0.002 | -0.002,0.006 | 110 | 1.12 | 0.27 | 0.39 |
| | Corridors | intercept | 0.27 | -0.06,0.59 | 110 | 1.62 | 0.11 | |
| | | *age* | 1.18x10$^{-4}$ | -0.002,0.002 | 110 | 0.11 | 0.92 | 0.97 |
| Waning lh OTS-limbs | numbers | intercept | 0.40 | 0.10,0.70 | 120 | 2.61 | 0.01 | |
| | | *age* | 1.93x10$^{-4}$ | -0.002,0.002 | 120 | 0.19 | 0.85 | 0.93 |
| | *words* | intercept | 0.34 | -0.02,0.69 | 120 | 1.86 | 0.06 | |
| | | *age* | 0.002 | -5.9x10$^{-4}$, 0.004 | 120 | 1.50 | 0.14 | 0.25 |
| | *limbs* | intercept | 1.76 | 1.40,2.11 | 120 | 9.74 | 7.04x10$^{-17}$ | |
| | | *age* | -0.005 | -0.008,-0.003 | 120 | -4.47 | 1.81x10$^{-5}$ | 2.26x10$^{-4}$ |
| | *bodies* | intercept | 0.62 | 0.39,0.85 | 120 | 5.34 | 4.49x10$^{-7}$ | |
| | | *age* | -2.99x10$^{-5}$ | -0.002,0.002 | 120 | -0.04 | 0.97 | 0.97 |
| | *Adult faces* | intercept | -0.02 | -0.22,0.18 | 120 | -0.17 | 0.86 | |
| | | *age* | 0.002 | 4.6x10$^{-4}$, 0.003 | 120 | 2.64 | 0.01 | 0.043 |
| | *Child faces* | intercept | 0.12 | -0.19,0.44 | 120 | 0.77 | 0.45 | |
| | | *age* | 0.002 | -3.4x10$^{-4}$, 0.004 | 120 | 1.67 | 0.10 | 0.20 |
| | *Cars* | intercept | -0.01 | -0.29,0.26 | 120 | -0.09 | 0.93 | |
| | | *age* | 0.001 | -6.4x10$^{-4}$,0.003 | 120 | 1.32 | 0.19 | 0.30 |
| | *String instr.* | intercept | 0.11 | -0.09,0.31 | 120 | 1.06 | 0.29 | |
| | | *age* | 0.002 | 1.2x10$^{-4}$, 0.003 | 120 | 2.15 | 0.03 | 0.11 |
| | *Houses* | intercept | -0.049 | -0.31,0.21 | 120 | -0.37 | 0.71 | |
| | | *age* | 0.001 | -6.4x10$^{-4}$,0.003 | 120 | 1.29 | 0.20 | 0.31 |
| | *Corridors* | intercept | -0.18 | -0.41,0.04 | 120 | -1.61 | 0.11 | |
| | | *age* | 7.14x10$^{-4}$ | -8.6x10$^{-4}$, 0.002 | 120 | 0.90 | 0.37 | 0.51 |
| Emerging lh pFus-faces | numbers | intercept | -0.02 | -0.61,0.57 | 103 | -0.07 | 0.95 | |
| | | *age* | 0.004 | 3.8x10$^{-4}$, 0.009 | 103 | 2.2 | 0.03 | 0.11 |

| | | | | | | | | |
|---|---|---|---|---|---|---|---|---|
| | *words* | intercept | -0.06 | -0.63,0.51 | 103 | -0.21 | 0.83 | |
| | | *age* | 0.006 | 0.002,0.01 | 103 | 2.78 | 0.006 | 0.04 |
| | *limbs* | intercept | 0.49 | -0.02,0.99 | 103 | 1.91 | 0.06 | |
| | | *age* | 0.001 | -0.003,0.005 | 103 | 0.57 | 0.57 | 0.71 |
| | *bodies* | intercept | 0.13 | -0.25,0.51 | 103 | 0.66 | 0.51 | |
| | | *age* | 0.003 | 3.5x10$^{-4}$, 0.006 | 103 | 2.25 | 0.03 | 0.1 |
| | *Adult faces* | intercept | -0.12 | -0.46,0.22 | 103 | -0.69 | 0.49 | |
| | | *age* | 0.006 | 0.004,0.009 | 103 | 5.33 | 5.92x10$^{-7}$ | 1.48x10$^{-5}$ |
| | *Child faces* | intercept | -0.30 | -0.84,0.25 | 103 | -1.08 | 0.28 | |
| | | *age* | 0.011 | 0.007,0.015 | 103 | 5.79 | 7.74x10$^{-8}$ | 3.87x10$^{-6}$ |
| | *Cars* | intercept | 0.02 | -0.36,0.40 | 103 | 0.11 | 0.91 | |
| | | *age* | 0.003 | 5.1x10$^{-4}$, 0.006 | 103 | 2.36 | 0.02 | 0.08 |
| | *String instr.* | intercept | -0.07 | -0.42,0.28 | 103 | -0.41 | 0.69 | |
| | | *age* | 0.003 | 7.7x10$^{-4}$, 0.006 | 103 | 2.62 | 0.01 | 0.04 |
| | *Houses* | Intercept | -0.29 | -0.94,0.36 | 103 | -0.88 | 0.38 | |
| | | *age* | 0.005 | -7.3x10$^{-5}$, 0.009 | 103 | 1.95 | 0.05 | 0.15 |
| | *Corridors* | intercept | -0.30 | -0.72,0.11 | 103 | -1.46 | 0.15 | |
| | | *age* | 0.003 | -4.0x10$^{-4}$, 0.005 | 103 | 1.71 | 0.09 | 0.19 |
| *Waning rh OTS-limbs* | numbers | intercept | 0.78 | 0.33,1.23 | 98 | 3.41 | 9.51x10$^{-4}$ | |
| | | *age* | -0.002 | -0.005,9.8x10$^{-4}$ | 98 | -1.34 | 0.18 | 0.30 |
| | *words* | intercept | 0.47 | 0.04,0.90 | 98 | 2.17 | 0.03 | |
| | | *age* | -5.53x10$^{-4}$ | -0.003,0.002 | 98 | -0.38 | 0.70 | 0.84 |
| | *limbs* | intercept | 1.88 | 1.34,2.42 | 98 | 6.90 | 5.13x10$^{-10}$ | |
| | | *age* | -0.006 | -0.009,-0.002 | 98 | -2.95 | 0.004 | 0.025 |
| | *bodies* | intercept | 1.01 | 0.65,1.38 | 98 | 5.51 | 2.91x10$^{-7}$ | |
| | | *age* | -0.001 | -0.004,0.001 | 98 | -1.09 | 0.28 | 0.39 |
| | *Adult faces* | intercept | 0.11 | -0.19,0.41 | 98 | 0.72 | 0.47 | |
| | | *age* | 0.003 | 6.4x10$^{-4}$,0.005 | 98 | 2.62 | 0.01 | 0.04 |
| | *Child faces* | intercept | 0.34 | -0.12,0.80 | 98 | 1.47 | 0.14 | |
| | | *age* | 0.003 | -2.3x10$^{-5}$,0.006 | 98 | 1.97 | 0.05 | 0.15 |
| | *Cars* | intercept | 0.32 | -0.01, 0.65 | 98 | 1.90 | 0.06 | |
| | | *age* | -4.05x10$^{-4}$ | -0.003,0.002 | 98 | -0.35 | 0.72 | 0.84 |
| | *String instr.* | intercept | 0.43 | 0.14,0.72 | 98 | 2.94 | 0.004 | |
| | | *age* | -8.54x10$^{-4}$ | -0.003,0.001 | 98 | -0.86 | 0.39 | 0.53 |
| | *Houses* | Intercept | 0.46 | 0.08,0.85 | 98 | 2.40 | 0.02 | |
| | | *age* | -0.002 | -0.004,9.6x10$^{-4}$ | 98 | -1.26 | 0.21 | 0.32 |
| | *Corridors* | intercept | 0.24 | -0.05,0.54 | 98 | 1.63 | 0.11 | |
| | | *age* | -0.002 | -0.004,1.3x10$^{-4}$ | 98 | -1.86 | 0.07 | 0.16 |
| *Emerging rh pFus-faces* | numbers | intercept | 0.74 | 0.26,1.22 | 94 | 3.08 | 0.003 | |
| | | *age* | -8.03x10$^{-4}$ | -0.004,0.003 | 94 | -0.47 | 0.64 | 0.78 |
| | *words* | intercept | 0.52 | 0.016,1.03 | 94 | 2.05 | 0.04 | |
| | | *age* | 1.18x10$^{-4}$ | -0.004,0.004 | 94 | 0.07 | 0.95 | 0.97 |
| | *limbs* | intercept | 1.18 | 0.70,1.66 | 94 | 4.91 | 3.85x10$^{-6}$ | |
| | | *age* | -0.003 | -0.006,6.8x10$^{-4}$ | 94 | -1.59 | 0.12 | 0.21 |
| | *bodies* | intercept | 0.77 | 0.34,1.20 | 94 | 3.52 | 6.61x10$^{-4}$ | |
| | | *age* | 1.07x10$^{-4}$ | -0.003,0.003 | 94 | 0.07 | 0.94 | 0.97 |

| | | | | | | | |
|---|---|---|---|---|---|---|---|
| *Adult faces* | intercept | 0.28 | -0.09,0.65 | 94 | 1.48 | 0.14 | |
| | *age* | 0.005 | 0.002,0.008 | 94 | 3.68 | $3.92x10^{-4}$ | 0.003 |
| *Child faces* | intercept | 0.45 | -0.03,0.92 | 94 | 1.87 | 0.06 | |
| | *age* | 0.008 | 0.005,0.011 | 94 | 4.75 | $7.42x10^{-6}$ | $1.24x10^{-4}$ |
| *Cars* | intercept | 0.39 | -0.02,0.80 | 94 | 1.87 | 0.06 | |
| | *age* | 0.001 | -0.002,0.004 | 94 | 0.76 | 0.45 | 0.59 |
| *String instr.* | intercept | 0.19 | -0.10,0.48 | 94 | 1.27 | 0.21 | |
| | *age* | 0.002 | $-2.8x10^{-4}$,0.004 | 94 | 1.72 | 0.09 | 0.19 |
| *Houses* | Intercept | 0.56 | 0.03,1.10 | 94 | 2.09 | 0.04 | |
| | *age* | $-4.67x10^{-4}$ | -0.004,0.003 | 94 | -0.24 | 0.81 | 0.92 |
| *Corridors* | intercept | 0.45 | -0.08,1.0 | 94 | 1.70 | 0.09 | |
| | *age* | -0.003 | -0.006,0.001 | 94 | -1.39 | 0.17 | 0.29 |

**Supplementary Table 10.**

Likelihood ratio tests determining if the selectivity of the dependent variable is better predicted by one or two predictors. Listed by figure order. Related to Fig. 4 and Fig. S13.

| ROI | No. | Model | df | LRStat | *p* |
|---|---|---|---|---|---|
| Emerging left pOTS-words | 1 | Words ~ limbs + (1\|subj) | 4 | | |
| | 2 | Words ~ limbs + faces + (1\|subj) | 5 | 26.3 | $2.92 \times 10^{-7}$ |
| Waning left OTS-limbs | 1 | Limbs ~ faces + (1\|subj) | 4 | | |
| | 2 | Limbs ~ faces + words (1\|subj) | 5 | 4.76 | 0.029 |
| Emerging left pFus-faces | 1 | Faces ~ limbs + (1\|subj) | 4 | | |
| | 2 | Faces ~ limbs + words + (1\|subj) | 5 | 6.23 | 0.013 |
| Waning right OTS-limbs | 1 | Limbs ~ faces + (1\|subj) | 4 | | |
| | 2 | Limbs ~~faces + words (1\|subj) | 5 | 5.03 | 0.025 |
| Emerging right pFus-faces | 1 | Faces ~ limbs + (1\|subj) | 4 | | |
| | 2 | Faces ~ limbs + words + (1\|subj) | 5 | 17.45 | $2.95 \times 10^{-5}$ |

**Supplementary Table 11.**

Fixed effects parameters for LMMs testing if selectivity to faces, words and limbs is linked in developing parts or ROIs. Listed by figure order. Related to Fig. 4 and Fig. S13.

| ROI | parameter | β | CI | df | t | *p* |
|---|---|---|---|---|---|---|
| Emerging left pOTS-words | Intercept | 3.18 | 2.80,3.56 | 109 | 16.64 | $1.081x10^{-31}$ |
| | limbs | -0.48 | -0.64,-0.31 | 109 | -5.78 | $7.267x10^{-8}$ |
| | faces | -0.36 | -0.49,-0.23 | 109 | -5.44 | $3.236x10^{-7}$ |
| Waning left OTS-limbs | Intercept | 3.28 | 2.89,3.68 | 119 | 16.43 | $2.23x10^{-32}$ |
| | words | -0.21 | -0.40,-0.02 | 119 | -2.20 | 0.030 |
| | faces | -0.32 | -0.49,-0.15 | 119 | -3.67 | 0.0004 |
| Emerging left pFus-faces | Intercept | 4.05 | 3.57,4.54 | 102 | 16.57 | $1.066x10^{-30}$ |
| | words | -0.33 | -0.58,-0.07 | 102 | -2.54 | 0.013 |
| | limbs | -0.47 | -0.73,-0.21 | 102 | -3.61 | 0.0005 |
| Waning right OTS-limbs | Intercept | 3.24 | 2.90,3.59 | 97 | 18.77 | $4.59x10^{-34}$ |
| | words | -0.24 | -0.46,-0.03 | 97 | -2.28 | 0.025 |
| | faces | -0.32 | -0.45,-0.19 | 97 | -4.80 | $5.74x10^{-6}$ |
| Emerging right pFus-faces | Intercept | 3.81 | 3.27,4.36 | 93 | 13.90 | $1.97x10^{-24}$ |
| | words | -0.58 | -0.84,-0.32 | 93 | -4.38 | $3.16x10^{-5}$ |
| | limbs | -0.46 | -0.81,-0.12 | 93 | -2.68 | 0.009 |

**Supplementary Table 12.**

Mean t-values of predictor values and predicted values of dependent variables in developing parts of ROIs. Listed by figure order. Related to Fig. 4 and Fig. S13.

| ROI | Predictor variables (PV) | Mean selectivity (t) of PV in 5-9yo, (SD) | Mean selectivity (t) of PV in 13-17yo, (SD) | Dependent variable (DV) | Predicted selectivity (t) of DV for 5-9yo | Predicted selectivity (t) of DV for 13-17yo |
|---|---|---|---|---|---|---|
| Emerging left pOTS-words | limbs | 2.20 (1.28) | 0.26 (0.95) | words | 2.77 | 4.01 |
| | faces | -1.76 (1.35) | -2.65 (2.11) | | | |
| Waning left OTS-limbs | faces | -1.23 (1.52) | -0.62 (1.85) | limbs | 3.54 | 3.21 |
| | words | 0.63 (1.40) | 1.27 (1.22) | | | |
| Emerging left pFus-faces | limbs | 1.78 (0.89) | -0.82 (0.86) | faces | 2.91 | 4.15 |
| | words | 0.95 (1.17) | 0.88 (1.29) | | | |
| Waning right OTS-limbs | faces | 0.23 (2.43) | 1.18 (3.35) | limbs | 3.31 | 2.98 |
| | words | -0.56 (1.20) | -0.47 (1.25) | | | |
| Emerging right pFus-faces | limbs | 1.50 (0.97) | -0.14 (1.13) | faces | 3.34 | 4.13 |
| | words | -0.39 (1.54) | -0.45 (1.57) | | | |
